# Single-nucleus resolution mapping of the adult *C. elegans* and its application to elucidate inter- and trans-generational response to alcohol

**DOI:** 10.1101/2022.07.21.500524

**Authors:** Lisa Truong, Yen-Wei Chen, Rio Barrere-Cain, Karissa Shuck, Wen Xiao, Max T. Levenson, Eduardo da Veiga Beltrame, Blake Panter, Ella Reich, Paul W. Sternberg, Xia Yang, Patrick Allard

**Affiliations:** Human Genetics Graduate Program, UCLA, Los Angeles, CA. 90095; Molecular Toxicology Inter-Departmental Program, UCLA, Los Angeles, CA. 90095; Institute for Society & Genetics, UCLA, Los Angeles, CA. 90095; Department of Microbiology, Immunology, and Molecular Genetics, UCLA, Los Angeles, CA. 90095; Biology and Biological Engineering, California Institute of Technology, Pasadena, CA. 91125; Integrative Biology and Physiology Department, UCLA, Los Angeles, CA. 90095; Molecular Biology Institute, UCLA, Los Angeles, CA. 90095

**Author notes:** Authors contributed equally to the work (Lisa Truong, Yen-Wei Chen). **Author Contributions**, LT, RBC, KS, WX, MTL, BP, and EL performed biological experiments and corresponding analyses, YWC performed bioinformatic analyses, PA and XY supervised experiments, PA and XY supervised analyses, EB devised a visualization approach that assisted cell type assignment and cluster identification which PS supervised, LT, RBC, YWC, XY, and PA wrote the manuscript.

**Keywords:** *C. elegans*, snRNA-seq, ethanol, alcohol

## Abstract

Single-cell RNA transcriptomic platforms have significantly contributed to our understanding of tissue heterogeneity as well as of developmental and cellular differentiation trajectories. They also provide an opportunity to map an organism’s response to environmental cues with high resolution and unbiasedly identify the cell types that are the most transcriptionally sensitive to exposures. Here, we applied single nucleus RNA-seq experimental and computational approaches to *C. elegans* to establish the transcriptome of the adult nematode and comprehensively characterize the transcriptional impact of ethanol as a model environmental exposure on the entire organism at cell type-resolution over several generations. Clustering, tissue and phenotype enrichment, and gene ontology analyses identified 31 clusters representing a diverse number of adult cell types, including those from syncytial and multi-nucleated tissues which are difficult to assess by single cell RNA-seq, such as the mitotic and meiotic germline, hypodermal cells, and the intestine. We applied this method to identify the impact of inter- and trans-generational exposure to two human-relevant doses of alcohol. Cell type proportions were not significantly altered by ethanol. However, Euclidean distance analysis identified several germline, striated muscle, and neuronal clusters as being major transcriptional targets of ethanol at both the F1 and F3 generations although the relative order of clusters changed between generations. The impact on germline clusters was further confirmed by phenotypic enrichment analysis as well as functional validation, namely a remarkable inter- and trans-generational increase in germline apoptosis, aneuploidy, and embryonic lethality. Together, snRNA-seq of the adult *C. elegans* represents a powerful approach for the detailed examination of an adult organism’s response to environmental cues.

## INTRODUCTION

In mammals, *in utero* exposure to alcohol is associated with an array of well-characterized morphological, neurological, and reproductive deficits in the F1 progeny that are grouped into symptoms of Fetal Alcohol Syndrome Disorders (FASD) [1]. The plurality of the conditions associated with FASD reflects the variety of organ systems and processes showing structural and functional anomalies following prenatal alcohol exposure, such as the reproductive system, the central nervous system, craniofacial morphogenesis, the heart, kidney, liver, and gastrointestinal system (reviewed in [2,3]). However, while *in utero* alcohol exposure clearly impacts the function of multiple organ systems, a comprehensive assessment of all organs, tissues, and cell types that are the most affected by alcohol remains lacking [4].

In addition to impacting the health of the F1 progeny, mounting evidence in various model systems, such as mice, rats, *Drosophila*, and *C. elegans*, indicates that at least some exposure-related adverse reproductive and neurobehavioral features also extend beyond the F1 and are detectable in F3 progeny [5–8]. For instance, a rat model of late gestational ethanol exposure demonstrated that not only F1 but also F2 and F3 individuals show an average 50% increase in ethanol intake [9]. Moreover, preconception exposure is sufficient to cause increased alcohol intake in the offspring, together with signs of spatial learning and memory deficits [10]. Notably, the impact of ethanol on alcohol and substance use across several generations is observed in the broader context of several established multi- and transgenerational models in which various cognitive, behavioral, or physical endpoints are altered (reviewed in [11,12]).

*C. elegans* is a simplified but highly advantageous model for studying the effects of alcohol and is the most used invertebrate species for modeling FASD (reviewed in [13]).Direct exposure to ethanol causes a variety of dose- and duration-dependent outcomes similar to those elicited in mammals such as growth and fertility impairments, neuro-depressive effects, increase alcohol preference and disinhibition, withdrawal, all supported by the involvement of similar cellular and neurological pathways [14–18]. The *C. elegans* genome is also equipped with the conserved gene families of alcohol and aldehyde dehydrogenases that provide the main metabolic activity towards ethanol which is first processed into acetaldehyde and subsequently into acetate [19]. Finally, its reproductive system, with two gonads opening into a common uterus where embryos initiate their development, provides a window for *in utero* exposure to alcohol.

Recently, the combination of single-cell RNA sequencing (scRNA-seq) technologies and the tractability of the model organism *C. elegans*, with its well-established differentiation lineages and timing, has enabled the layering of transcriptional data with developmental events at both embryonic and larval (L2) stages, as well as the mapping of the entire nervous system [20–22]. This has led, for example, to the identification of gene expression changes that track the development of 502 preterminal and terminal cell types in embryos [21] and the characterization of 27 distinct cell types at larval stages [20]. Furthermore, we and others have shown that *C. elegans* is also a powerful model for the study of multi- and trans-generational responses to environmental stimuli [23–28]. However, single cell transcriptomic approaches have yet to be applied to the characterization of environmental exposures, including alcohol, at the whole organism level and across generations.

Here, we used RNA-seq from single nuclei to maximize the isolation of diverse cell types, including from the approximately 30% of all somatic cells that are polyploid and from the mostly syncytial adult germline. We applied this approach to examine the transcriptional impact of parental (P0) exposure to two physiologically relevant doses of ethanol on the F1 offspring (inter-generational exposure) as well as on the F3 generation (trans-generational exposure). We show that single nucleus RNA-seq (snRNA-seq) identifies a large number of distinct cell types that resolve into well-characterized cellular and functional identities. We also demonstrate that this powerful method can provide insights into the effect of inter- and trans-generational exposure to ethanol at tissue and cell-type specific resolution and identify the cells and molecular pathways that are most impacted by such exposures.

## MATERIAL AND METHODS

### Culture conditions and strains

The strain JK560 *fog-1(q253)* was used for sequencing and single molecule fluorescence *in situ* hybridization (smFISH) experiments. N2 (wild type) worms were used for embryonic lethality and acridine orange apoptosis experiments. The strain TY2441 (*Pxol-1:: gfp; rol-6(pRF4)*) was used for X chromosome aneuploidy experiments. Worms were cultured on standard nematode growth medium (NGM) plates streaked with single colony OP50 *E. coli* and maintained at 20°C. The generation of worms to be collected for single-nucleus analysis was moved to 25°C at the L1 stage and grown at 25°C for 48 hours until the beginning of day 2 of adulthood. To collect L1 larvae from the F1 generation, worms were synchronized by bleaching P0 gravid adults and having their F1 embryos subsequently grown for 16 hours at 20°C at which time all F1 were at the L1 stage. To collect L1 larvae from the F3 generation, the L1 larvae were collected through filtration using a 10μm nylon mesh filter (EDM Millipore NY1102500 and EDM Millipore SX00025000) which only allows L1 stage worms to pass through. L1 worms for both generations were kept for 48 hours at 25°C and then washed with five rounds of M9 buffer and centrifuged at 1,300g for 1 minute to pellet worms between each wash. After the final wash, worms were spun in a rotator with 1mL of M9 for 30 minutes to remove OP50 from the worms’ gut. The worms were then allowed to settle by gravity for 5 minutes and the final compact worm pellet volume was adjusted to 30μL. The aforementioned strains were obtained from the *C. elegans* Genetics Center (CGC): JK560 *fog-1(q253)* I; TY2441 *yls34 (Pxol-1::gfp+rol-6 (pRF4))* (obtained by crossing *him-8(e1489)* out of TY2431); N2: wild-type.

### *C. elegans* ethanol exposure

For ethanol exposures, a population of gravid adult worms was bleached. The embryos obtained were plated on standard OP50 seeded NGM plates and allowed to grow to the L4 stage (approximately 50 hours post bleaching). Nematodes were exposed for 48 hours in a liquid culture containing M9 buffer solution, standard OP50 bacteria (10mg/mL), and ethanol at a final concentration of 0.05% or 0.50% in 15mL conical tubes. Following liquid exposure, the F1 progeny of the exposed P0 generation was split into two sets of plates, one for collecting the F1 adults and one for maintenance until the F3 generation. This scheme ensured the comparability of F1 and F3 snRNA-seq data.

### Single-nucleus dissociation

All single-nucleus dissociation steps were done at 4°C. A compact 30μL pellet of adult *JK560 C. elegans* (approximately 4,000 worms) was transferred to a prechilled Dounce homogenizer (Sigma Z378623-1EA) and homogenized for 10 strokes with 400μL of ice cold FA lysis buffer (50mM HEPES/NaOH pH 7.5, 1mM EDTA, 0.1% Triton X-100, 150mM NaCl, protease inhibitor 0.5X (Roche 11697498001), RNase inhibitor 0.2U/μL (Thermo Fisher 10777019), and RNase free water). Worms were homogenized for 10 strokes in a 1.5mL Wheaton Dounce homogenizer and for an additional 20 strokes (350μL FA buffer) with an Eppendorf Dounce homogenizer (Fisher 06-434) in a corkscrew fashion. Between each set of 10 homogenization strokes, debris was pelleted using 100g for 1 minute and supernatant containing nuclei was collected and pooled in a fresh 1.5mL low retention microcentrifuge tube.

After homogenization, the pooled supernatant containing nuclei was centrifuged at 100g for 1 minute to pellet remaining debris. The top 900μL of supernatant with nuclei was transferred to a fresh 1.5mL low retention microcentrifuge tube and washed once with 1% PBS BSA ((Thermo Fisher AM9624) that was filtered with a 0.22μm pressure filter (Thermo Scientific 03-377-26, Fisher SLGP033RS)). Nuclei were pelleted at 500g for 4 minutes, resuspended in 750-850μL 1% PBS-BSA, and then filtered using a 40μm Flowmi tip filter (Sigma-Aldrich BAH136800040-50EA).

Nuclei integrity was verified by staining single-nuclei isolations with DAPI and observing the nuclei under a fluorescent microscope. The nuclei did not have a frayed appearance and were compact. Nuclei extractions were performed at 4°C in a timely fashion to prevent cellular transcription during the dissociation process. On average, a total of 1,200 nuclei is obtained per batch of 4,000 worms. Flow cytometry was used to ensure optimal nuclei concentration (700-1200 events/μl).

### Single molecule fluorescence *in situ* hybridization

Single molecule fluorescence *in situ* hybridization (smFISH) was performed on F1 and F3 adults that were maintained, exposed, and filtered in a similar manner to animals used in the single-nucleus dissociation protocol. SmFISH was performed using the protocol developed by the Kimble Lab [29]. Probes were designed and ordered through Stellaris and are compiled in Table S1. Both probes were used at a final concentration of 0.25μM with approximately 100 dissected worms per condition. Samples were mounted on slides using fluoroshield with DAPI (Sigma-Aldrich F6057). Slides were imaged on the Leica SP8 confocal microscope. Fluorescence images were quantified using FISH-Quant v3 [30].

### Embryonic lethality assessment, *xol-1::*gfp analysis, and apoptosis assay

Embryonic lethality was performed on wild-type N2 F1 and F3 worms. At both generations, L4s were singled out and moved onto individual 33mm plates. Embryonic lethality was performed by monitoring the number of embryos produced each day and the subsequent larvae that hatched from these embryos for each individual worm spanning its entire reproductive lifespan. *Pxol-1::gfp* analysis was done by fluorescent microscopy on 24-hours post-L4 F1 and F3 adults and the occurrence of GFP+ embryos (expressing *Pxol-1::gfp*) was recorded. The proportion of XOL-1::GFP+ was calculated by dividing the number of worms with at least 1 GFP+ embryo by the total number of worms analyzed [31]. Apoptosis assays were performed on wild-type N2 worms by Acridine Orange staining of synchronized adult hermaphrodites collected at 20-24 hours post-L4 at the F1 and F3 generations as previously described [32,33].

**Expanded Material and Methods, including computational analyses, can be found in the supplemental information section.**

## RESULTS

### SnRNA-seq identifies a wide array of defined cell types in the adult *C. elegans*

Intact nuclei were isolated from adult *fog-1(q253) C. elegans* raised at the restrictive temperature of 25°C [34]. Since the focus of our study was on the characterization of adult tissues response to ethanol, this sperm-defective strain was used to prevent self-fertilization and the crowding of our snRNA-seq data with embryonic cell types (see material and methods section). Briefly, worms were synchronized and allowed to grow to day one of adulthood before mechanical nuclear extraction (Figure 1A). Nuclei concentration was determined using flow cytometry and nuclear integrity was assessed by high-resolution microscopy. Single nucleus RNA-seq library preparation was performed using the 10X Genomics Chromium system followed by 50 PE sequencing on the Illumina Novaseq 6000 platform. In total, we generated transcriptomic data for 81,267 nuclei, each with more than 500 transcripts derived from 31 groups collected in 5 distinct batches. On average, 2,181 unique molecular identifiers (UMIs) and 992 genes were detected per nucleus with high sequencing depth (90.3% average sequencing depth) (Figure S1).

**Figure 1:**
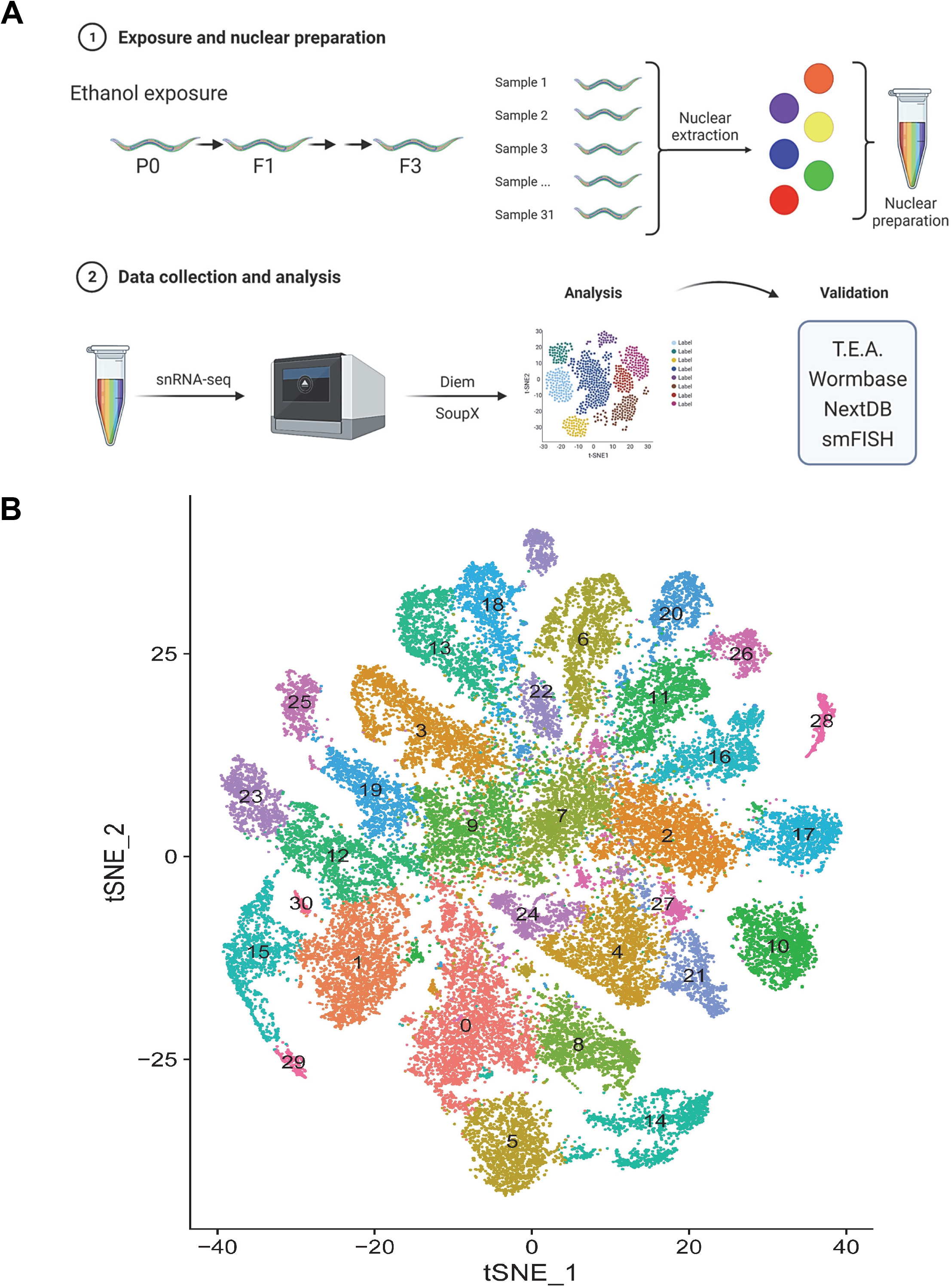
Adult *C. elegans* snRNA-seq sample preparation, analysis, and t-SNE projection. **A.** Experimental flow for single-nucleus isolation and snRNA-seq analysis. **B.** t-distributed stochastic neighbor embedding (t-SNE) plot of cells from all the samples with clustering through unsupervised Louvain clustering. Cluster molecular characterization and identity is presented in the corresponding dashboard (Supplemental file 1).

The snRNA-seq reads were demultiplexed and aligned to the ENSEMBL ce10 *C. elegans* transcriptome to generate gene expression matrices using CellRanger (10x Genomics) (see supplemental material and methods). To mitigate the inclusion of debris-contaminated droplets and to correct for ambient RNA contamination, we also applied DIEM [35] and SoupX [36], respectively. DIEM identifies and removes droplets containing high levels of extranuclear RNA through modeling semi-supervised expectation maximization and outperforms other methods in snRNA-seq [35]. We then combined DIEM with SoupX which models contamination levels of snRNA-seq with ambient RNA and corrects expression for the remaining droplets. Using these stringent pipelines, we retained transcriptomic data from 41,750 droplets representing a median of 1,627 UMIs and 1,007 genes. A total of 31 discrete clusters were identified following batch/group effect correction by canonical correlation analysis (CCA) in Seurat v3 followed by Louvain clustering algorithm [37,38]. Log-normalized expression levels in t-distributed stochastic neighbor embedding (t-SNE) plot projections were used to visualize cell clusters in two dimensions and dot heatmap were used to visualize marker expression across different cell types (Figure 1B and Figure 2).

**Figure 2:**
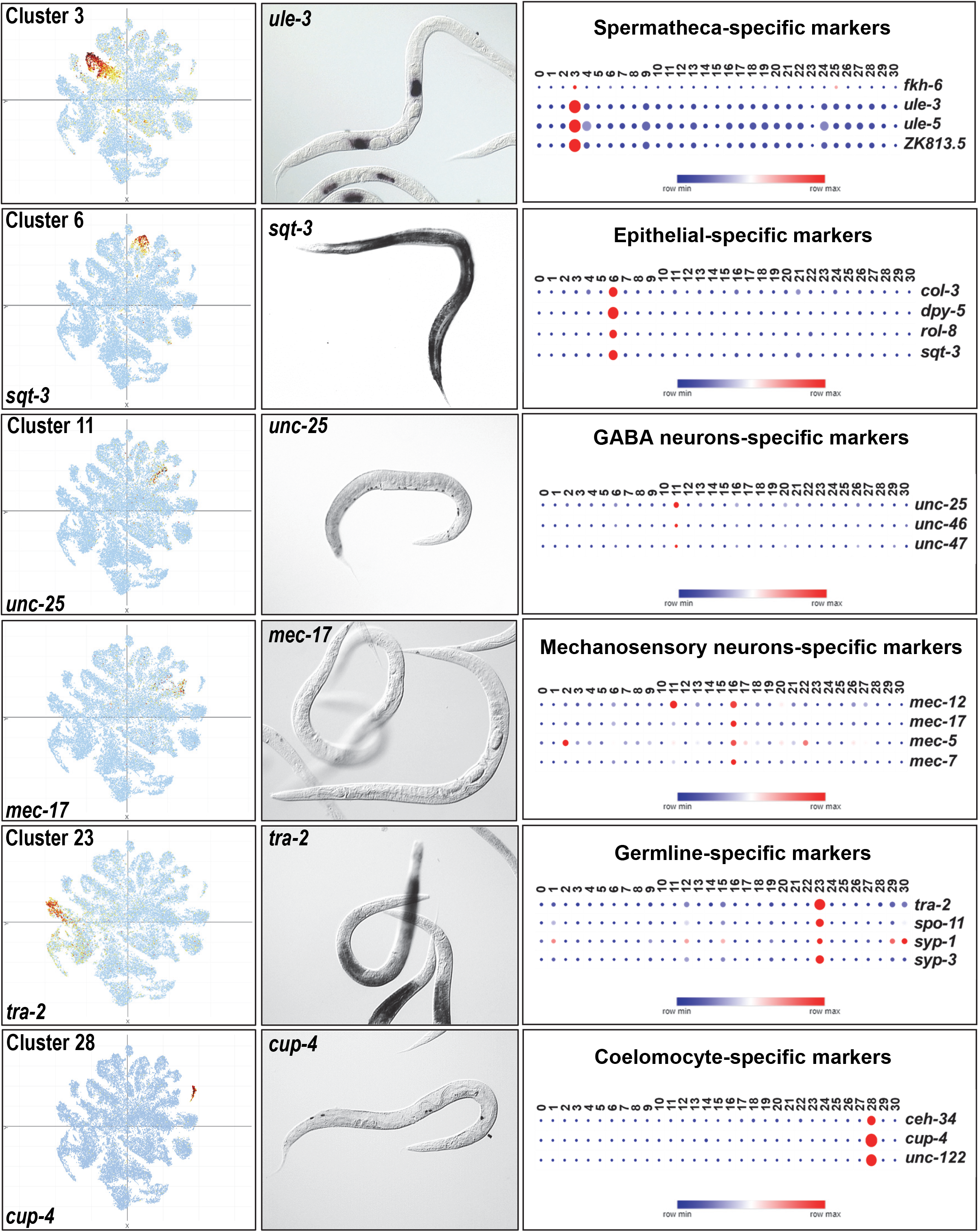
Cluster-resolved expression of cell type-specific markers. Example of cluster-specific expression of cell type specific markers for the spermatheca, epithelial cells, GABA and mechanosensory neurons, germline, and coelomocyte. Right column: cluster specific expression. Middle column: representative *in situ* expression data from NEXTDB for corresponding marker. Left column: gene expression dot plot of known cell type specific markers.

To facilitate unbiased cluster identification, gene enrichment (FDR<0.05) in each cluster was used to mine the Tissue enrichment, Gene Ontology, and Phenotype enrichment modules of WormBase [39] (Figure 1A). The output from these modules was cross-referenced with *in situ* expression data of the top enriched transcripts using the Nematode Expression Pattern Database (NEXTDB https://nematode.nig.ac.jp/) [40].

Comprehensive information for all 31 clusters that include top enriched and depleted genes, the 3 WormBase modules outputs, and representative examples from NEXTDB are presented in a dashboard (Supplemental data file 1). Examples are shown in Figure 2 alongside expression values of several known markers of each cell type showing notably high concordance. For example, cluster 23 is the only cluster showing a high expression level of the germline-specific transcripts *tra-2* [41], the meiotic-specific endonuclease *spo-11*, and the synaptonemal complex components *syp-1* and *syp-3* (as well as *syp-2* and *syp-4* not depicted here) [42,43]. In the adult *C. elegans* germline, the expression of *tra-2* is restricted to the most distal section of the germline and overlaps with that of the Notch signaling target *sygl-1* [29,44]. To test whether cluster 23 could correspond to the distal germline, we confirmed *sygl-1* expression by small molecule FISH (smFISH) as previously described [29] and compared its expression with its tSNE distribution (Figure S2). As expected, *sygl-1* showed a high expression level in the region corresponding to cluster 23 with some expression in other germline clusters which matched its smFISH distribution and published data [44] indicating that cluster 23 likely corresponds to the distal germline.

Similarly, cluster 3 specifically expresses all known markers of the spermatheca such as *fkh-6, ule-3, ule-5*, and also *ZK813.7* [45,46]. Marker analyses also revealed that sub-regionalization within clusters is apparent. For example, the three GABAergic neuron markers *unc-25, unc46*, and *unc-47* [47] resolve in one section of cluster 11 (Figure 2) which also expresses markers of cholinergic neurons (*e.g. cha-1* and *cho-1*) [48]. Also of note, based on WormBase’s TEA analysis, several tissue- and cell-types are represented by multiple clusters (*e.g*. the germline represented by 6 clusters, the intestine and epithelial system, represented by 3 cluster each) (Supplemental data file 1) likely reflective of the cellular heterogeneity underlying the composition of those tissues. Very few clusters (2/31), *i.e*. clusters 0 and 9, showed a lack of concordance between TEA, GO, and *in situ* data precluding assignment of a clear identity. All snRNA-seq data can be mined at: https://singlecell.broadinstitute.org/single_cell/reviewer_access/d94dacb1-9ab3-49f4-a5a0-392d8a0399ca (also see data availability section).

### SnRNA-seq reveals broad impacts of inter-generational exposure to ethanol

We first applied snRNA-seq to identify the organism-wide transcriptional outcome of a parental 48-hour (L4 to end of day 1 of adulthood) exposure to two concentrations of ethanol (0.05% and 0.5%) or water control on the F1 adult progeny. These doses were chosen to circumscribe the wide range of human blood alcohol concentrations associated with low (0.05%) or high alcohol (0.5%) use respectively [49]. We first compared cell type proportion in the F1 following parental ethanol exposure and observed that broadly similar cell type distributions were observed across all treatment conditions (Figure S3). However, we observed a significant number of Differentially Expressed Genes (DEGs) (FDR<0.05) between treatment conditions (Figure 3A). Across all F1 clusters from the 0.05% ethanol exposure condition, we identified a total of 1,223 DEGs, including 583 uniformly upregulated DEGs, 520 uniformly downregulated DEGs, and 120 DEGs that were differentially up or down regulated in cluster specific ways (*i.e*. upregulated in some clusters but downregulated in other clusters) (Table S2). Surprisingly, compared to 0.05%, exposure to the higher ethanol concentration of 0.5% resulted in fewer DEGs identified at the F1 (Table S3) with a total of 948 DEGs, including 430 uniformly upregulated DEGs, 407 uniformly downregulated DEGs, and 111 up- and down-regulated DEGs (Figure 3A).

**Figure 3:**
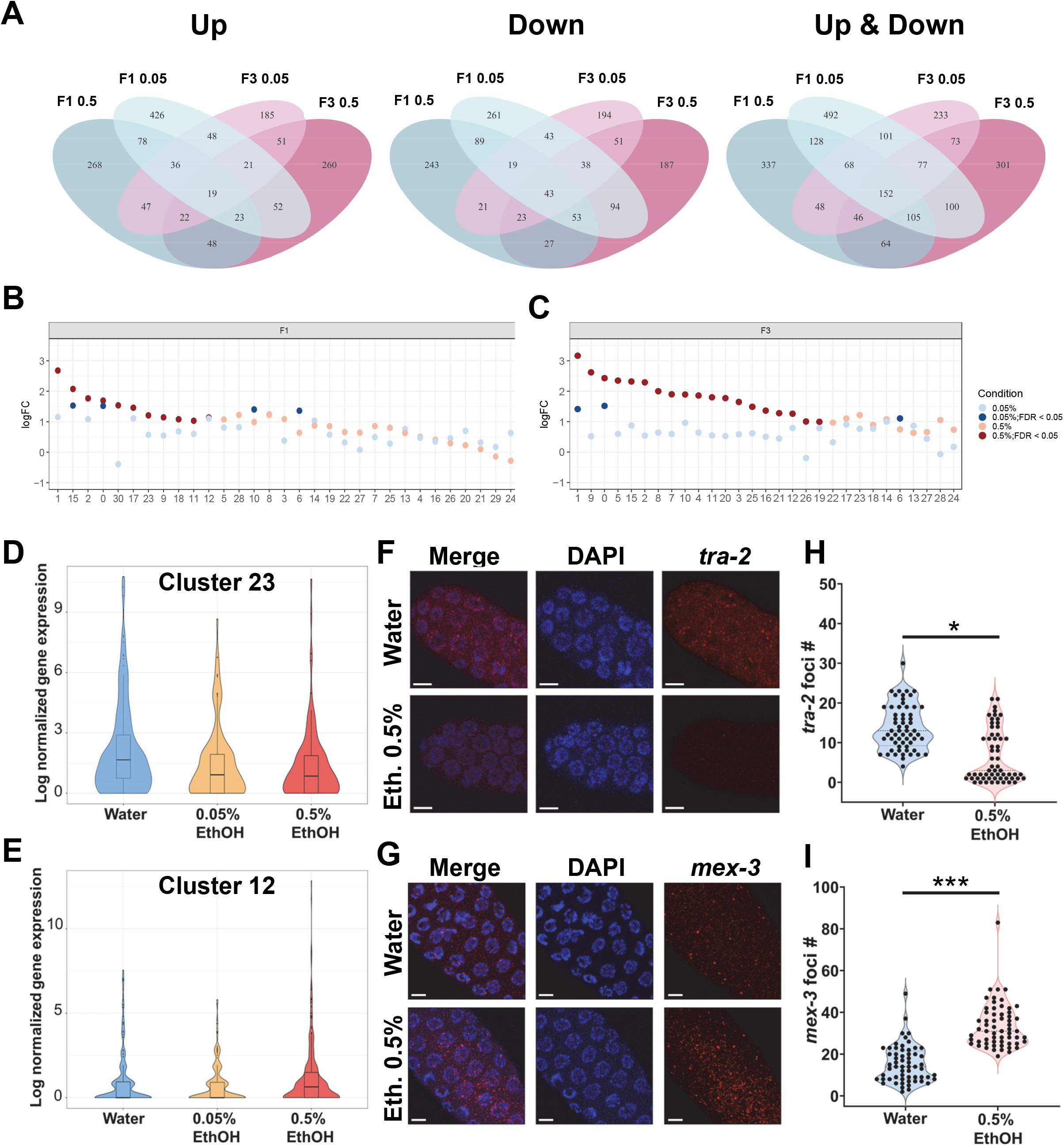
Global and cluster-specific alterations in gene expression by inter- or trans-generational ethanol exposure. **A.** Venn diagram based on the union of DEGs across all the cell types, separated by Up-regulated DEGs only (left), Downregulated genes only (right), and all DEGs (middle). **B, C.** Euclidean distance sensitivity analysis of all the cell clusters at the F1 generation (B) or F3 generation (C). X-axis indicates cluster number and y-axis indicates log fold change compared to Euclidean distance obtained by permuting treatment labels. Significance was assessed based on comparing Euclidean distance against 1,000 random permutated labels. **D, E.** Expression of *tra-2* in cluster 23 at the F1 (D) and *mex-3* in cluster 12 at the F3 (E). **F-I.** Validation of snRNA-seq data through single molecule fluorescence *in situ* hybridization (smFISH) by confocal imaging (F,G) followed by FISH-Quant (H,I). Scale bar 5 μm. 3 biological replicates, 2 worms per repeat, 10 nuclei per germline. *P<0.05, ***P<0.001. Welch’s t-test.

Gene Ontology of the union of all DEGs revealed the enrichment of some functional categories that align with alcohol metabolism such as the GO category “carboxylic acid metabolic process” driven by the presence in the DEG list of several aldehyde dehydrogenases (Table 1 and Table S4), which catalyze the final step of ethanol metabolism from acetaldehyde into acetate. However, a major target of ethanol across exposure conditions and clusters is the translation machinery as exemplified by the deregulation of many ribosomal components and representing 5 of the top 10 GO categories, including the top 3 (Table S4). The inhibition of translation and the downregulation of genes encoding ribosomal subunits are a well described and conserved impact of alcohol exposure *in vitro* and in a variety of species from bacteria to humans [50–55]. In addition, reproductive pathways were amongst the top shared across exposure conditions at the F1, such as “gamete generation”, “germ cell development”, and “embryo development ending in birth or egg hatching” pathways (Table S4).

**Table 1:**
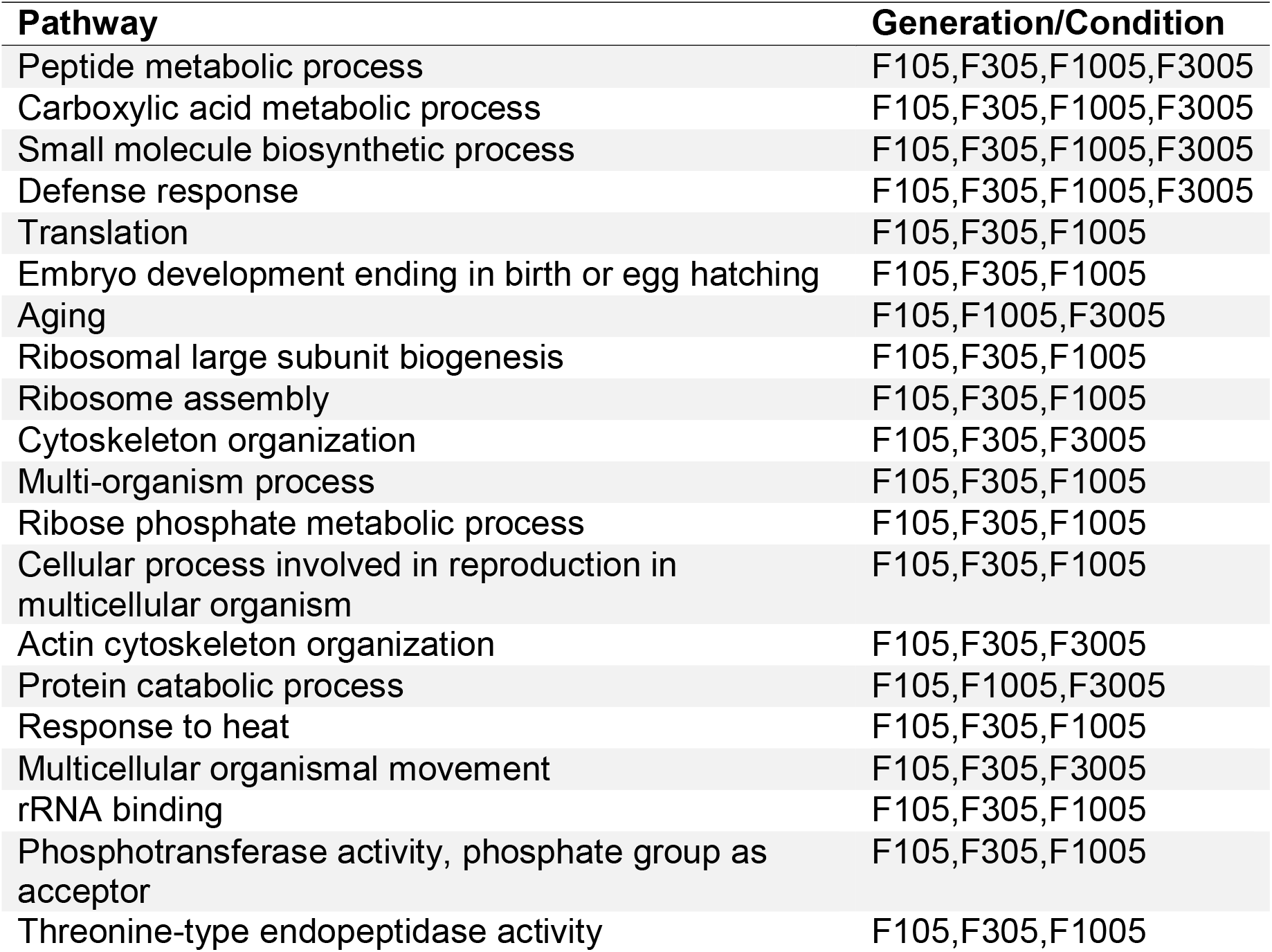
Shared pathways identified through DEG analysis across both exposures and generations. Top 20 shared pathways identified by the union of all cell type specific DEGs across the four conditions. F105 indicates 0.5% ethanol exposure at the F1 generation. F1005 indicates 0.05% ethanol exposure at the F1 generation. F305 indicates 0.5% ethanol exposure at the F3 generation. F3005 indicates 0.05% ethanol exposure at the F3 generation.

| **Pathway** | **Generation/Condition** |
| --- | --- |
| Peptide metabolic process | F105,F305,F1005,F3005 |
| Carboxylic acid metabolic process | F105,F305,F1005,F3005 |
| Small molecule biosynthetic process | F105,F305,F1005,F3005 |
| Defense response | F105,F305,F1005,F3005 |
| Translation | F105,F305,F1005 |
| Embryo development ending in birth or egg hatching | F105,F305,F1005 |
| Aging | F105,F1005,F3005 |
| Ribosomal large subunit biogenesis | F105,F305,F1005 |
| Ribosome assembly | F105,F305,F1005 |
| Cytoskeleton organization | F105,F305,F3005 |
| Multi-organism process | F105,F305,F1005 |
| Ribose phosphate metabolic process | F105,F305,F1005 |
| Cellular process involved in reproduction in multicellular organism | F105,F305,F1005 |
| Actin cytoskeleton organization | F105,F305,F3005 |
| Protein catabolic process | F105,F1005,F3005 |
| Response to heat | F105,F305,F1005 |
| Multicellular organismal movement | F105,F305,F3005 |
| rRNA binding | F105,F305,F1005 |
| Phosphotransferase activity, phosphate group as acceptor | F105,F305,F1005 |
| Threonine-type endopeptidase activity | F105,F305,F1005 |

### SnRNA-seq reveals tissue-specific DEGs from intergenerational (P0 to F1) exposure to ethanol

Next, we conducted cluster-specific DEG analysis to investigate cell type specific effects at the F1. Cluster-resolved DEG analysis indicated clearly distinct transcriptional responses to parental ethanol exposure between cell types. While some genes were consistently up-regulated (*atp-6, nduo-6*) or down-regulated (*vit-5*) across all clusters between ethanol and water treatment, most DEGs showed cell type-specific restriction as highlighted by the low overlap of the top DEGs per cluster (Figure S4, Table S2 and S3). To rank order the F1 clusters by sensitivity to ethanol exposure, we employed a Euclidean distance analysis [56,57], which estimates the degree of transcriptomic shifts between exposure and control groups (see Supplemental Material and Methods). Several clusters (1, 15, and 30) with an assigned germline identity based on TEA/GO/Phenotype enrichment analyses and NEXTDB (Supplemental File 1) showed some of the largest degree of transcriptomic shifts at the F1 generation under 0.5% ethanol exposure condition (Figure 3B). Other cluster categories that appeared most affected included clusters related to muscle function such as cluster 2 and cluster 17, both carrying striated muscle cell identity. The degree of transcriptomic shift was much less pronounced following 0.05% ethanol exposure compared to 0.5% ethanol, suggesting a dose-dependent transcriptomic response across cell types.

We hypothesized that while most DEGs are cell type-specific, genes implicated in ethanol metabolism may show a more uniform response across clusters. Thus, we investigated the expression of genes involved in ethanol metabolism, including 3 distinct alcohol dehydrogenase (*sodh-1*, H24K24.3, ZK829.7) and 10 aldehyde dehydrogenase (*alh-3, −4, −7* through −*13*), whose expression was detectable in our datasets (Figure S5). Contrary to our expectations, of the 13 genes examined, only 5 showed significant changes in expression (FDR < 0.05) and did so in a cluster- and dose-dependent fashion. For example, *sodh-1* was upregulated in cluster 13 and cluster 18 under the 0.05% exposure condition but was downregulated in cluster 2 and cluster 27 at 0.5%. Notably, the cell types showing the highest increase in ethanol metabolism genes were not the cell types that were the least sensitive to ethanol and vice versa, suggesting that the upregulation of ethanol metabolism genes in the F1 does not protect a tissue from the inter-generational impact of exposure (compare Figure 3B and S5).

At the F1, the majority of the clusters reaching statistical significance (FDR<0.05) after 0.5% ethanol exposure in our Euclidean Distance analysis displayed a germline identity, *e.g*. clusters 1, 12, 15, 23, 30. Thus, we next examined whether reproduction-related phenotypes were significantly over-represented in our dataset. We analyzed the top 10 most shared WormBase phenotypes across cell types with significantly altered Euclidean distance and identified several phenotypic terms related to reproduction (*e.g*. “diplotene region organization variant”, “pachytene region organization variant”, “germ cell compartment expansion variant”) that are mildly upregulated in germline cluster 1 but strongly and uniformly downregulated in germline cluster 12 (Figure 4A). Finally, we examined whether the DEGs across the sensitive cell types were enriched in specific phenotypes by comparing the DEGs’ phenotype enrichment outcome with all WormBase phenotypes. This analysis revealed that our dataset has a significantly higher proportion of phenotypes related to reproductive system development, cellular development, and morphology categories among both treatment groups (Figure 4B).

**Figure 4:**
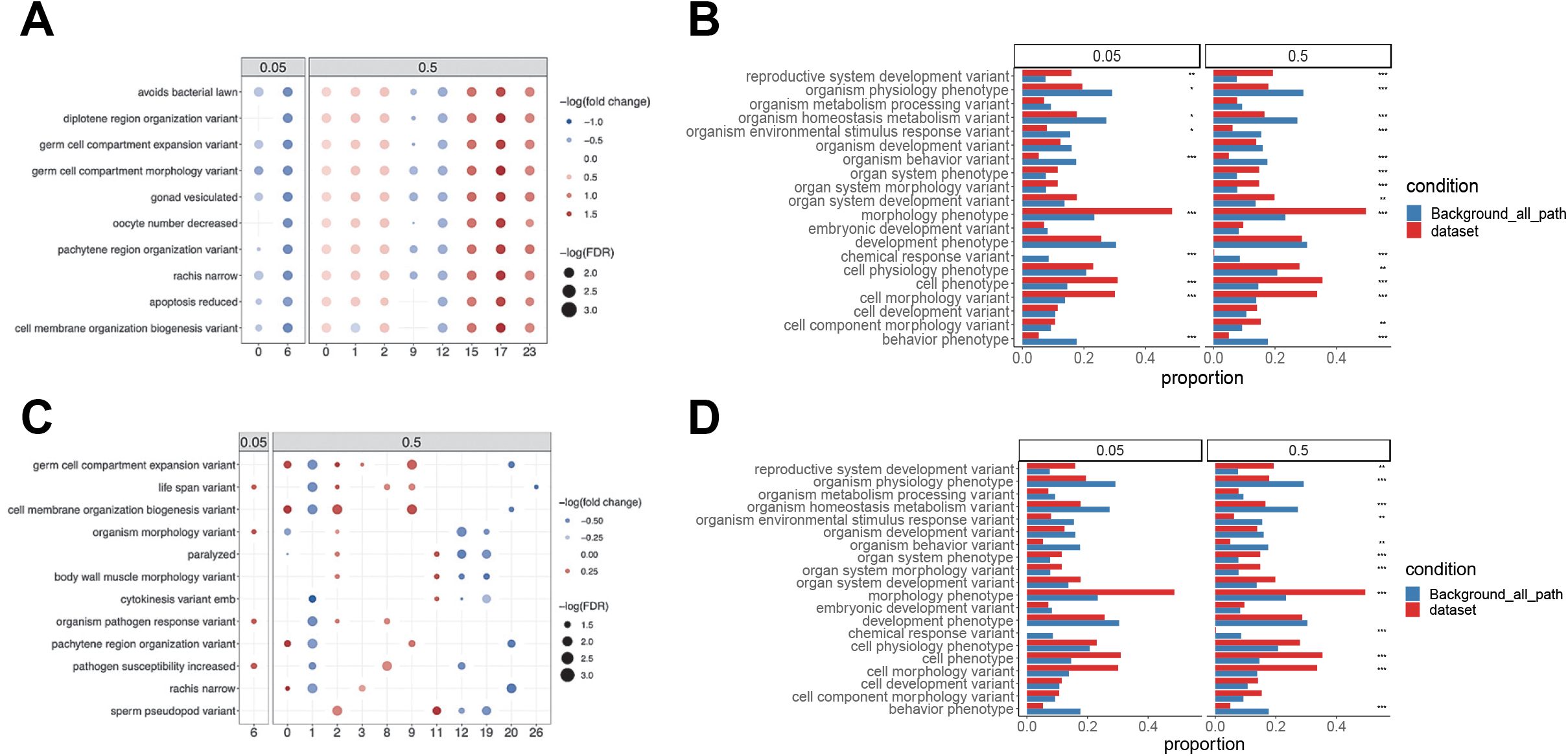
Phenotypic enrichment in response to inter- or trans-generational ethanol exposure. **A,C.** Dot heatmap of top shared WormBase phenotype across cell types with significantly altered Euclidean distance metric at the F1 (A) or F3 (C). Dot size corresponded to -log(FDR) obtained from enrichment analysis and dot color corresponded -log(median fold change) of overlapping genes in each pathway. **B,D**. Bar plot showing the proportion of top WormBase phenotype annotations from all enriched pathways (“dataset”) and WormBase phenotype database (“Background_all_path”) at the F1 (B) or F3 (D). For each WormBase phenotype from the original database we retrieved the corresponding WormBase phenotype annotations by querying EBI OLS API, followed by selecting the top 20 shared phenotypes. Proportions were calculated based on the proportion of annotations among all enriched pathways (“dataset”) and the WormBase phenotype database (“Background_all_path”). Fisher’s exact test was used to compare proportions between the two conditions in each annotation category.

Since the germline appears to be a major target for ethanol at the F1 and F3 generations, we validated the magnitude and directionality of the transcriptional impact of ethanol exposure in that tissue. For example, in our snRNA-seq dataset, *tra-2* showed a significant downregulation of its expression in cluster 23 in the F1 under 0.5% ethanol exposure (Figure 3D, Table S3, 0.60-fold change, adjusted P value = 0.003). We performed smFISH in dissected F1 germlines followed by quantification using FISH-quant (Figure 3F-I). We observed a congruent 0.47-fold downregulation of *tra-2* level (P<0.05). Conversely, at the F3 under 0.5% ethanol exposure condition, *mex-3* was significantly upregulated in our snRNA-seq dataset in germline cluster 12 (Figure 3E, Table S6, 1.58-fold change, adjusted P value = 0.008). Cluster 12 also as a meiotic germline identity based on GO enrichment (Supplemental file 1). SmFISH indicated an upregulation of *mex-3* in the mid-pachytene region of F3 *C. elegans* gonads (2.19-fold, P<0.001).

Together, these results indicate a strong intergenerational impact of alcohol in *C. elegans* on a variety of cell types including those that belong to the germline.

### Transgenerational (P0 to F3) impact of ethanol

We extended our snRNA-seq approach to the F3 generation to capture the transgenerational effect of P0 exposure to ethanol. Similarly to the F1, no overt impact on cell type distributions was observed (Figure S3). Across all clusters at the F3 stemming from a P0 0.05% ethanol exposure, a total of 798 unique Differentially Expressed Genes (DEGs), 366 unique upregulated DEGs, 369 unique downregulated DEGs satisfying an FDR<0.05 and 63 DEGs that were differentially up or down regulated in cluster specific ways (Figure 3A, Table S5). For 0.5% ethanol, a total of 918 unique Differentially Expressed Genes (DEGs) were identified comprising 402 unique upregulated DEGs, 422 unique downregulated DEGs satisfying an FDR<0.05 and 94 DEGs that were differentially up or down regulated in cluster specific ways (Figure 3A, Table S6).

Gene Ontology analysis of all F3 DEGs revealed the enrichment of some functional categories that align with alcohol metabolism. These were exemplified by GO categories such as “carboxylic acid metabolic process”, “drug metabolic process”, and “small molecule catabolic process” driven in part by the presence in our DEG list of alcohol dehydrogenase genes, *sodh-1* and *hphd-1*, which catalyze the first step of ethanol metabolism from ethanol to acetaldehyde, as well as aldehyde dehydrogenase genes, *alh-8* and *alh-13*, which catalyzes the second step of ethanol metabolism from acetaldehyde into acetate, in both exposure groups as compared to water at the F3 (Table 1 and Table S4). Other highly enriched GO terms were “structural molecule activity”, “cytoskeleton organization”, “translation” and reproductive GO categories “embryo development ending in birth or egg hatching” and “sexual reproduction”.

### Cell type-specific transgenerational response to ethanol

Next, we conducted cluster specific DEG analysis to investigate ethanol’s transgenerational effects. While the majority of top DEGs at the F1 were up-regulated, in comparison, the majority of the top DEGs found at the F3 were down-regulated, especially at the 0.5% ethanol exposure condition (Figure S6). These top DEGs showed remarkable cell type specificity as highlighted by their low overlap across clusters. We assessed the sensitivity of individual clusters by measuring their Euclidean distance (Figure 3C). Notably, compared to the F1 results, more clusters reached the cut-off of FDR<0.05 from 0.5% exposure at the F3. While changes in the relative order of the clusters were observed between F3 and F1, several germline clusters showed some of the largest transcriptomic shifts including cluster 1 which remained the cluster the most transcriptionally affected by ethanol (Figure 3C).

The examination of F3 DEGs through the WormBase phenotype enrichment tool revealed a strong alteration of different phenotypes under the 0.5% ethanol exposure condition, albeit in a less uniform fashion across clusters (Figure 4C,D). The phenotype pachytene region organization variant was downregulated in cluster 1 but not in other germline clusters, suggesting a lasting but weakened impact of ethanol exposure effect at the F3. A comparison of phenotype proportion between our dataset and all phenotypes, revealed a persistent enrichment in the F3 of the phenotypic categories described in the F1 including a higher proportion of reproductive system development variant at 0.5% ethanol. By contrast to the F1, no phenotypic category reached significance under the 0.05% ethanol exposure condition, confirming the weakened impact of ethanol at the F3.

### Functional outcomes of ethanol’s inter- and trans-generational transcriptional impacts

The presence of several germline clusters amongst the most transcriptionally impacted clusters as well as the enrichment of some reproductive phenotypes at the F1 and F3 suggested that ethanol exposure may have a significant functional impact on reproduction at these generations. To test this hypothesis, we measured three hallmarks of reproductive health in *C. elegans*: germline apoptosis by acridine orange staining [32], the proper segregation of chromosomes during meiosis by monitoring the segregation of the X chromosome [58], and the viability of produced embryos through plate phenotyping [33,59]. All three reproductive measures were significantly upregulated at either 0.05% or 0.5% ethanol exposures in the F1 and in the F3 compared to the water control, remarkably, with no consistent dose-response relationship (Figure 5). Analysis of germline apoptosis through acridine orange staining revealed a 2-fold increase in the number of apoptotic nuclei per gonad in F1 worms who were exposed to 0.05% ethanol at the P0 (N=5, P<0.05) and a 2.7-fold increase in those who were exposed to 0.50% ethanol (N=5, P<0.0001) when compared to water. Similarly, at the F3, a 1.9-fold increase for both ethanol exposure conditions was observed when compared to water (N=4-5, P<0.001). Next, we monitored chromosome segregation through the segregation of the X chromosome using a strain carrying the *Pxol-1::gfp* reporter [31,58]. Chromosomes that fail to properly segregate during meiosis result in embryos with aneuploidies [43]. Here, we monitored aneuploidy *via* the incidence of male (XO) embryos which are caused by mis-segregation of the X-chromosome and marked by the expression of the male-specific *xol-1* promoter driving GFP. Analysis at the F1 identified a significant increase in the incidence of GFP-positive embryos for both ethanol exposure conditions when compared to water with a 2.6-fold increase in the proportion of worms with at least one GFP-positive embryo at 0.05% ethanol exposure (N=6-7, P<0.01) and a 2.8-fold increase at 0.50% ethanol exposure (N=6-7, P<0.01). The incidence of GFP-positive embryos further increased at the F3 with a 4-fold increase at 0.05% ethanol exposure (N=6-7, P<0.01) and a 4.3-fold increase at 0.50% ethanol exposure (N=6-7, P<0.001). Together, these results indicate a profound impact of inter- and trans-generational alcohol exposure on the nematode’s reproductive function congruent with the outcome of our snRNA-seq analysis.

**Figure 5:**
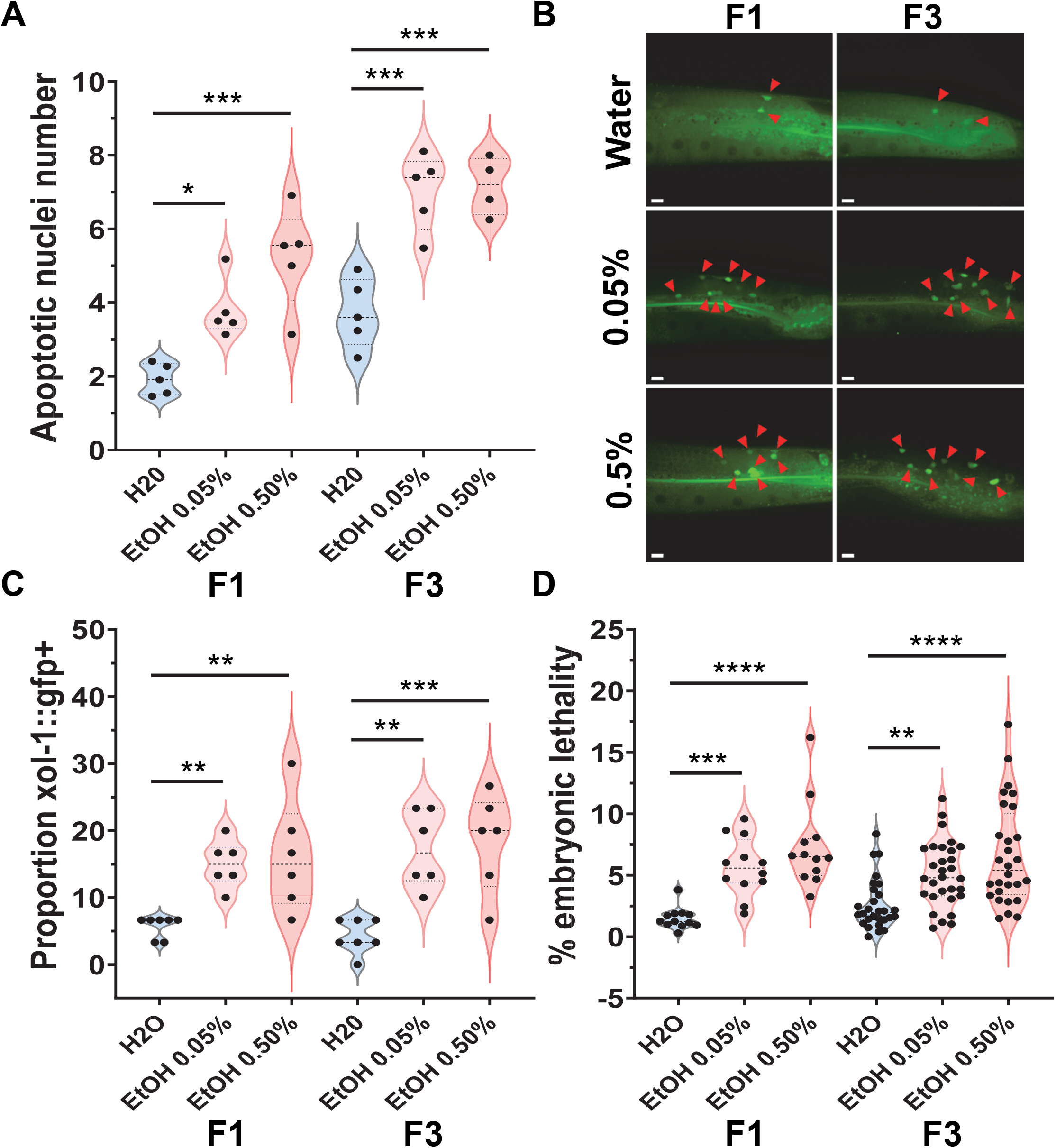
P0 ethanol exposure causes reproductive dysfunction at the F1 and F3 generations. P0 hermaphrodites were exposed to water, 0.05% ethanol, or 0.50% ethanol. **A,B.** Number of apoptotic nuclei per gonadal arm in N2 worms at the F1 and F3 exposed to the indicated ethanol levels, N=4-5, 22 worms per repeat. Scale bars = 10μm. **C.** Assessment of errors of X chromosome segregation as measured by *Pxol-1::gfp* reporter. Out of 30 total worms per repeat, the percent (%) of worms with at least 1 GFP+ embryo was recorded, N=6, 30 worms per repeat. **D.** Percent embryonic lethality per worm was measured for N2, N=4-10, 2-3 worms per repeat (D). One-way ANOVA with Dunnet correction. *P<0.05, **P<0.01, ***P<0.001, ****P<0.0001.

## DISCUSSION

We performed single-nucleus RNA-seq approach in the adult *C. elegans* nematode and identified a large number of transcriptionally distinct cell types. By applying this approach to the study of the inter- and trans-generational impacts of ethanol exposure, we also demonstrate its utility in achieving a nuanced understanding of transcriptional responses to environmental cues.

We circumvented some of the drawbacks associated with single nucleus approaches by employing multiple data clean-up pipelines to identify and remove debris-contaminated droplets and spurious signal from ambient RNA, a common artifact of snRNA-seq [35,36]. This stringent approach removed approximately half of all droplets but still generated a robust number of UMI and genes per nucleus when compared with other studies [60]. Our approach also generated a larger number of clusters corresponding to cell types previously under-represented in single cell RNA-seq studies in *C. elegans* because of their lack of cellularization, *e.g*. meiotic stages of the germline and hypodermal cells.

Our approach nonetheless has several limitations. By working in a *fog-1* mutant background, we were not able to identify sperm cells, highlighted by the absence of expression of canonical sperm markers in our dataset. This was considered a necessary trade-off to avoid the production of embryos and the crowding of snRNA-seq data with a large and diverse number of embryonic cell types. While it is possible that *fog-1*’s absence alters the transcriptional landscape of the germline, *fog-1(q253)* was chosen specifically because of the normal morphology and staging of the hermaphrodite germline in the *fog-1* mutant background [34]. Secondly, for a small number of clusters, tissue enrichment analysis did not delineate a clear cell type identity (2/31 clusters) either representing cell types with mixed identities or cell types for which other clustering methods would be beneficial. Nonetheless, the large majority of clusters bore a distinctive tissue identity not only corroborated by GO and phenotype enrichment analyses but also by the tissue-specific *in situ* expression pattern of genes showing the highest degree of cluster specificity.

The application of our snRNA-seq approach to the study of ethanol’s response across generations highlighted the complexity of organisms’ response to environmental cues. This aspect was exemplified by the diversity and specificity of GO categories by clusters (Table S4) and by the low degree of overlap of top DEGs between clusters (Figure S4 and S6). Because of their high ranking in the Euclidean analysis in both F1 and F3 generations, several tissue types stood out as being particularly affected by ethanol: the muscle system, neurons, and the germline. The direct impact of alcohol on skeletal and cardiac muscle cells is well described and a common outcome of chronic alcohol use [61–63]. An intergenerational effect of prenatal alcohol exposure on the musculature and muscle function has also been demonstrated and is referred to as fetal alcohol myopathy [64,65]. Interestingly, a proposed mechanism for this induced muscle cell dysfunction is an alteration of protein synthesis [66,67] which is represented by several GO categories (*e.g*. “peptide metabolic process”, “peptide biosynthetic process”, “translation”) in our DEG pathway analysis (Table S4). Supplementation of amino acids to facilitate translation processes might be tested as a method of improving the impact of ethanol on the F1 muscle cells. The mechanisms of transgenerational impact of alcohol on F3 musculature, however, is unclear and has not been previously described.

Similarly, cluster 11 which comprises GABAergic (*e.g. unc-25^+^)* and some cholinergic neurons (*e.g. cha-1*^+^) is also identified as being significantly altered by 0.5% ethanol at the F1 and F2. Interestingly, GABAergic neurons are also a well-known target of direct and intergenerational alcohol exposure in which alcohol leads to over stimulation of the GABA system leading to dampening of neuronal excitability [68–71]. To a lesser extent, cholinergic signaling has also been implicated in the intergenerational impact of ethanol on the nervous system [72–74]. Since in *C. elegans*,direct alcohol exposure is associated with a deregulation of cholinergic signaling and locomotory behavior [16,75], it will be important to also investigate the role of GABA signaling and its interaction with cholinergic signaling in regulating locomotion not only in a direct exposure paradigm but also across generations.

Germline clusters were ranked the highest in the Euclidean distance analysis at the F1 and F3 generations. We therefore focused our validation experiments on the germline and reproduction. Direct ethanol exposure has been known to cause aneuploidy in mammalian germ cells for many years [76–78], however whether these effects extend to the F1’s germline has remained uncertain. Our results clearly indicate that in *C. elegans*, both low and high concentrations of ethanol have a profound impact on reproductive function (germ cell apoptosis, aneuploidy, embryonic lethality) and that these impacts extend transgenerationally. We have recently demonstrated that the transgenerational reproductive effects of the environmental chemical Bisphenol A requires the alteration of the repressive histone marks H3K9me3 and H3K27me3 [23,25]. Intergenerational alcohol exposure, on the other hand, has been shown to lead to histone hyperacetylation through the metabolism of ethanol into acetate [79]. Thus, it is plausible that alcohol’s inter- and trans-generational outcomes described here may be initiated by the over-acetylation of histone in the germline. Finally, while we validated ethanol’s impacts on several DEGs by smFISH in the germline, a more comprehensive DEG validation in other tissues will be needed as well as comparison with cell type-specific intergenerational and transgenerational ethanol transcriptional outcome in mammalian models when such data becomes available.

Together, the application of snRNA-seq to the adult *C. elegans* represents a powerful approach for the comprehensive identification of cell types in the nematode and for probing the transcriptional impact of physiological and environmental changes.

## Supporting information

Supplemental File 1_Dashboard

Table S1 smFISH probes

Table S2 DEG_F1_0.05

Table S3 DEG_F1_0.50

Table S4 Pathway analysis

Table S5 DEG_F3_0.05

Table S6 DEG_F3_0.50

## DATA AVAILABILITY

All raw data is accessible on NCBI’s Gene Expression Omnibus (GEO), at: https://www.ncbi.nlm.nih.gov/geo/query/acc.cgi?acc=GSE208229

The data is also available through the Broad Single Cell Portal: https://singlecell.broadinstitute.org/single_cell/study/SCP922/single-nucleus-resolution-mapping-of-the-adult-c-elegans-and-its-application-to-elucidate-inter-and-trans-generational-response-to-alcohol

## ACKNOWLEDGEMENTS

The authors would like to thank Doug Arneson for input on single-nucleus sequencing parameters; Ingrid Cely, In Sook Ahn, and Graciel Diamante for discussions and trouble-shooting advice; Eyal Ben David for advice on single-cell/nuclei dissociation methods; Jessica Scholes, Jeffrey Calim, Felicia Codrea, and Salem Haile for guidance with single-nucleus cytometry; Michael Mashock and Marco De Simone for their library preparation and sequencing expertise. Matteo Pellegrini for advice and input on data analysis. We thank Judith Kimble, Tina Lynch, and Sarah Crittenden for advice on smFISH and providing the *sygl-1* probe. The *Caenorhabditis* Genetics Center (CGC) provided the strains used in this study. We thank Yuji Kohara for permission to use NEXTDB *in situ* data.

---

**Figure 1:**
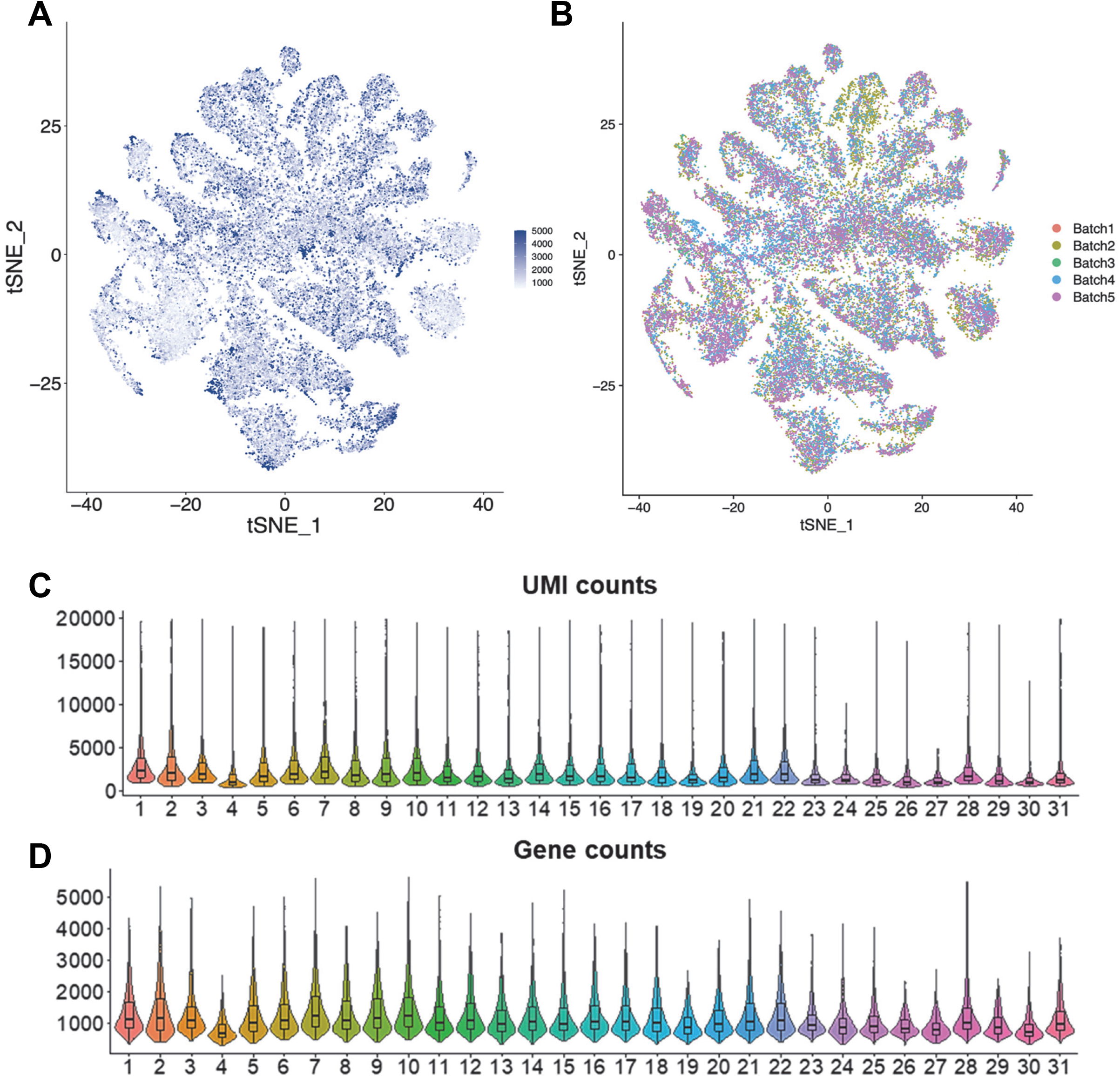
UMI, gene count, and batch distributions. **A.** UMI (unique molecular identifier) distribution as tSNE plot. X and Y-axis are representative tSNE axis. Each dot represents a cell and color scale Is plotted based on UMI count in each cell. UMI lower than 1000 are plotted as 1000 and higher than 5000 are plotted as 5000. **B,** tSNE plot colored by batch after batch effect correction by canonical correlation analysis. **C,D.** Violin and boxplot of UMI (unique molecular identifier) and gene counts per cell in each sample across 31 single-nuclei samples.

**Figure S2:**
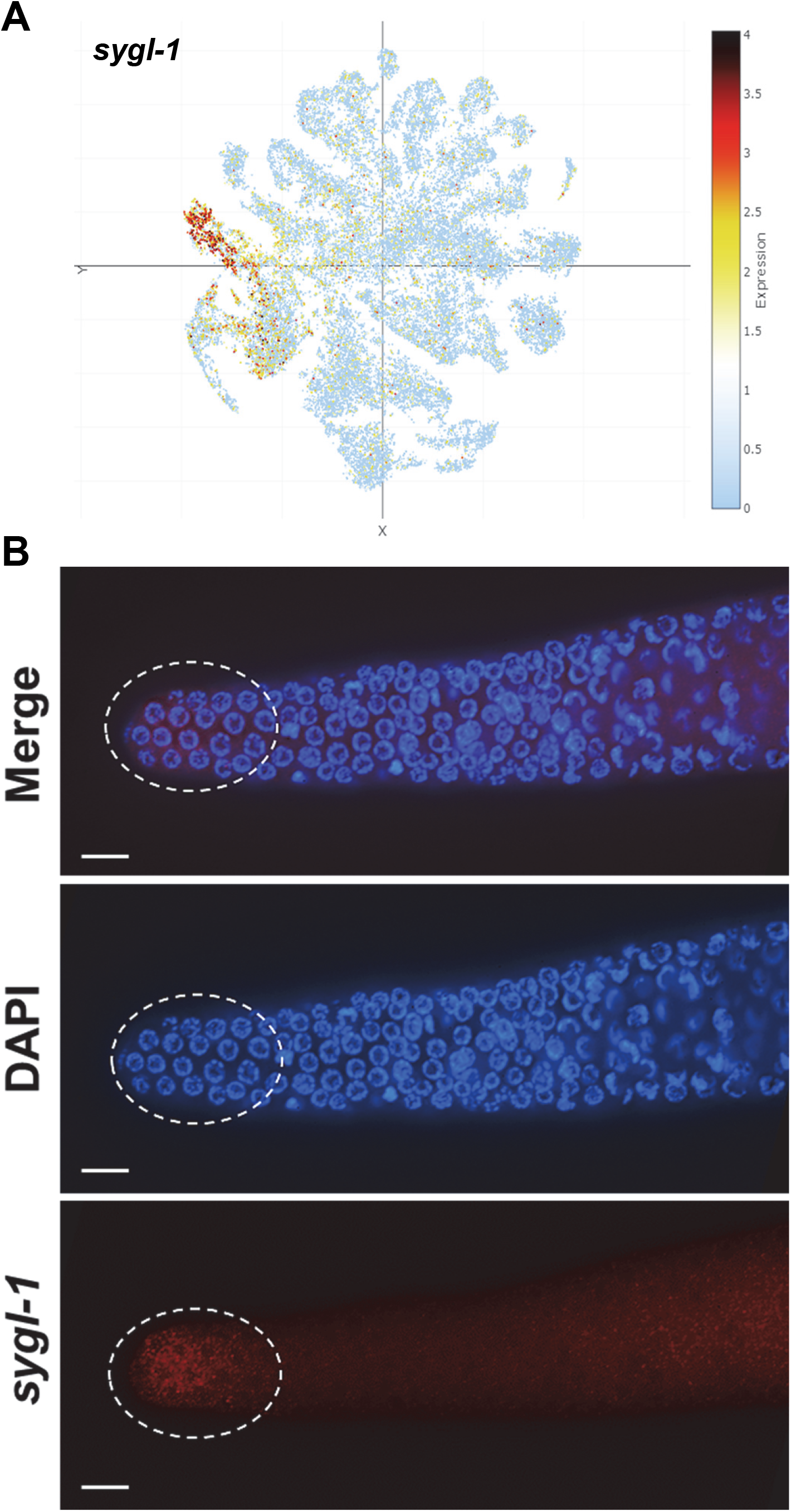
tSNE expression and smFISH for *sygl-1* transcript. **A.** Expression of *sygl-1* transcript from the adult *C. elegans* snRNA-seq data set. The bulk of its expression resolves to cluster 23. **B.** smFISH for *sygl-1* transcript showing its well-described expression within the distal gonad (circle) as well as some expression more proximally. Scale bar = 10μm.

**Figure S3:**
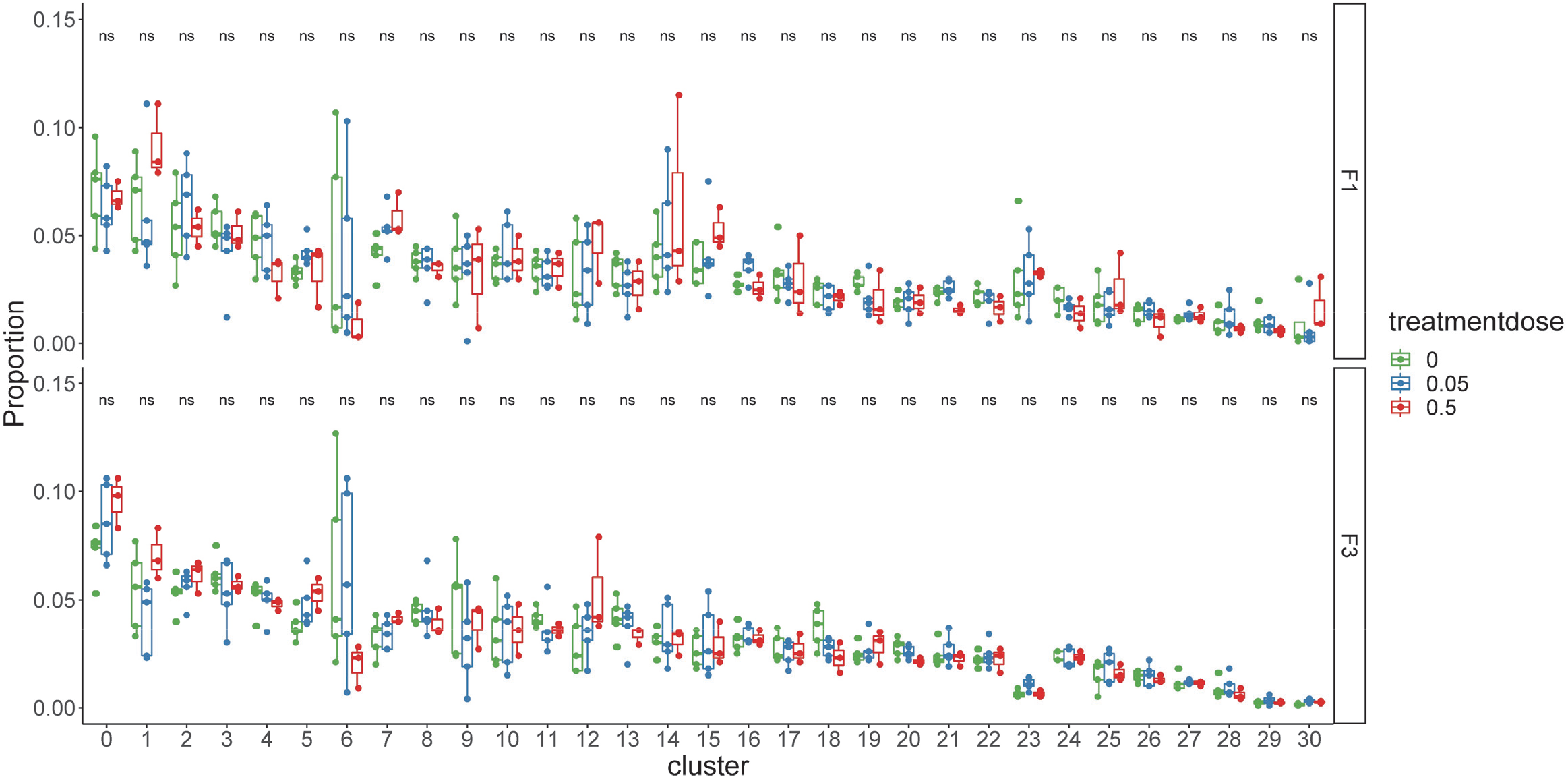
F1 and F3 cluster proportions. Proportion distribution plot of all F1 and F3 samples colored by different treatment dose (0, 0.05%, and 0.5% ethanol), each dot represents one sample. X-axis indicates cluster number assigned by Louvain clustering and Y-axis indicates cells of that cluster divided by all cells from that specific sample. All conditions were non-significant based on post-hoc Tukey statistic.

**Figure S4:**
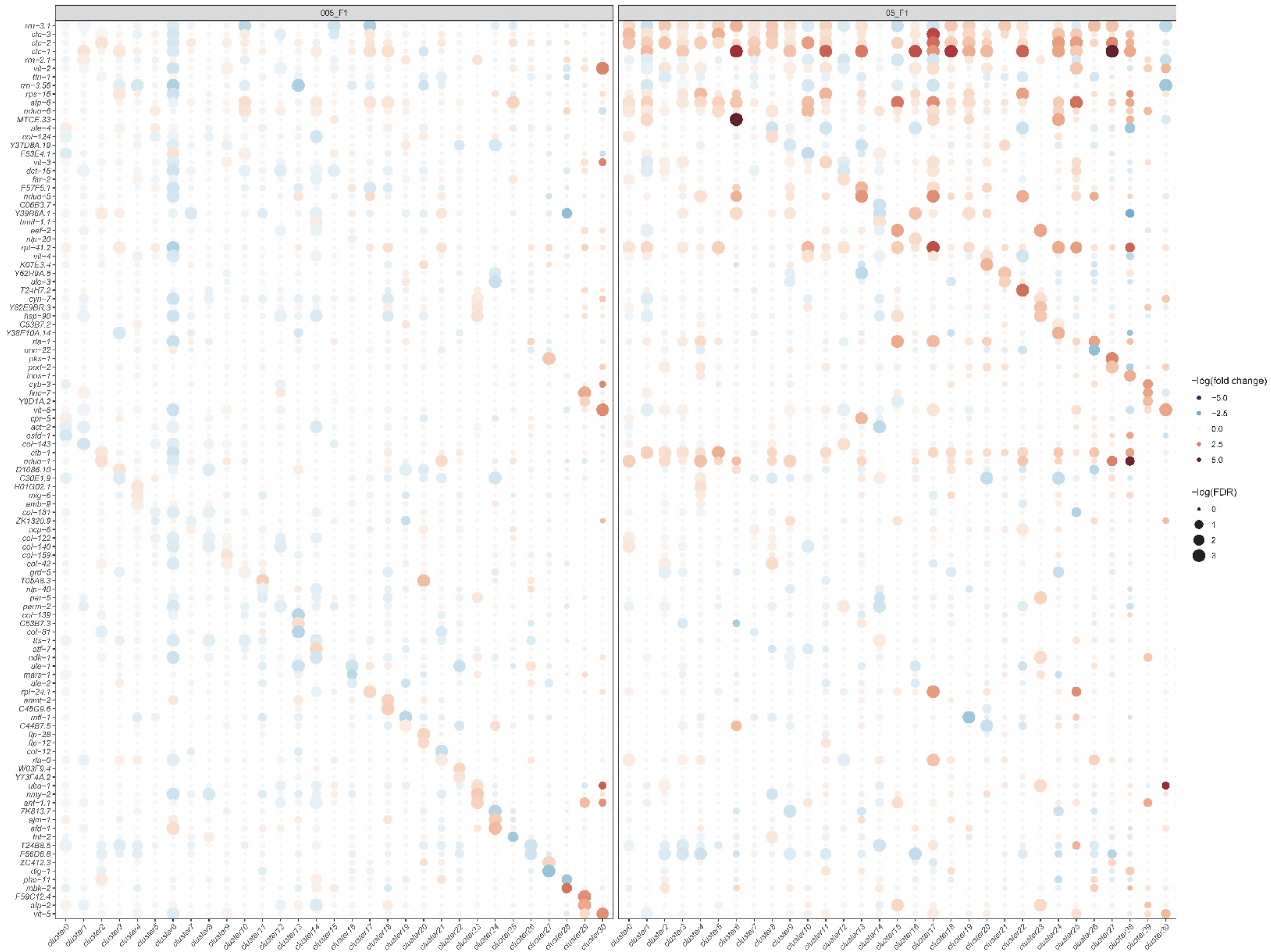
Dot heatmap of top 3 differential expressed genes across clusters at the F1. X-axis indicates different clusters representing different tissues and Y-axis indicates top 3 differentially expressed genes after ethanol treatment across clusters ranked by monocle based FDR. The plot is further separated by different ethanol doses (0.05% on the left and 0.5% on the right). The size of the dot is correlated to –log(FDR) of differential expression p-value and the color represents direction and scale of fold change with upregulation is shown in red and downregulation is shown in blue.

**Figure S5:**
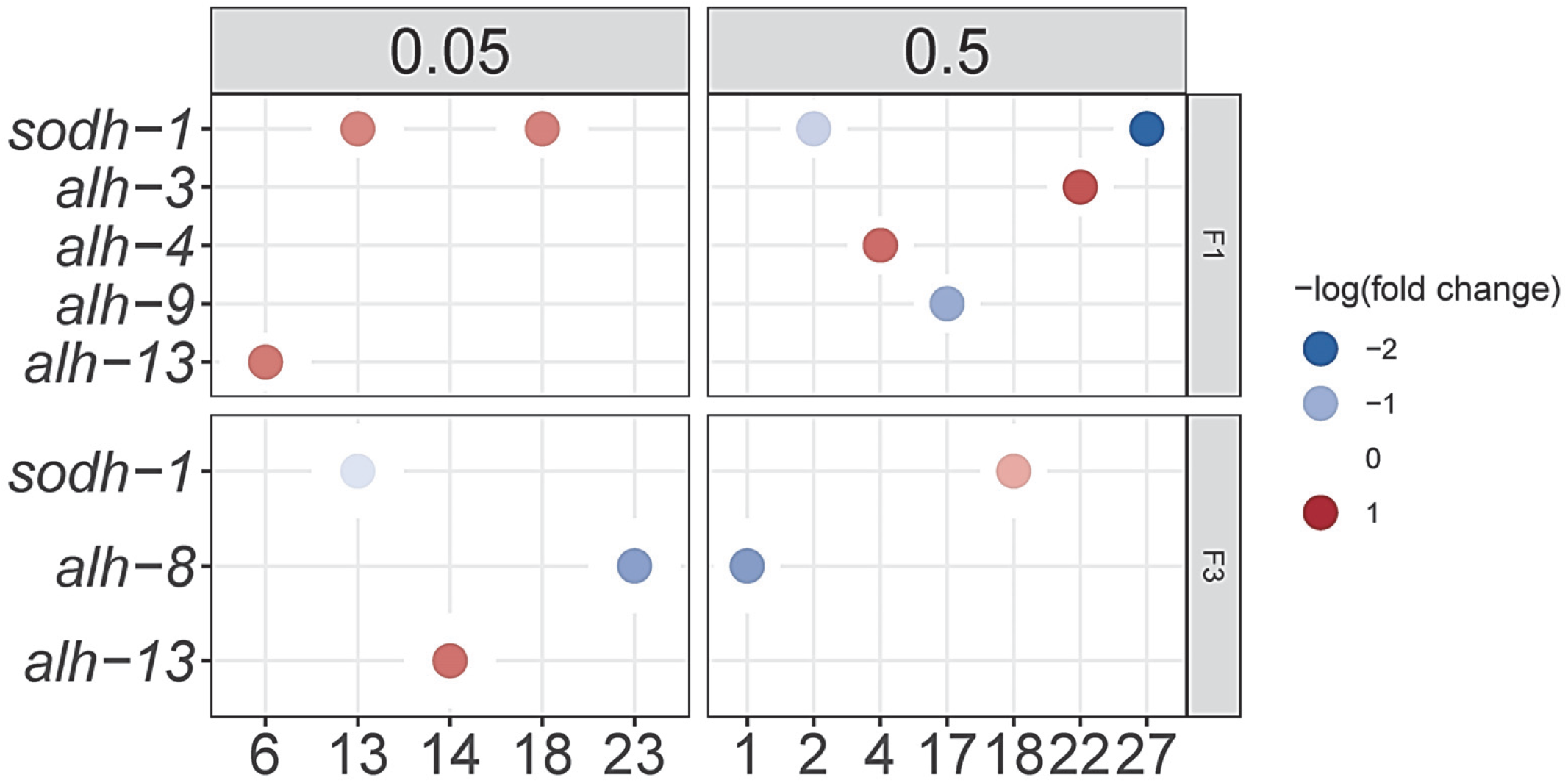
Cluster-specific DEGs related to ethanol metabolism at F1 and F3. Dot heatmap of ethanol metabolism related genes across different clusters, separated by exposure dose (0.05% or 0.5% ethanol) and generation (F1 or F3). Only DEGs with an FDR < 5% were plotted. Color indicates –log10(fold change).

**Figure S6:**
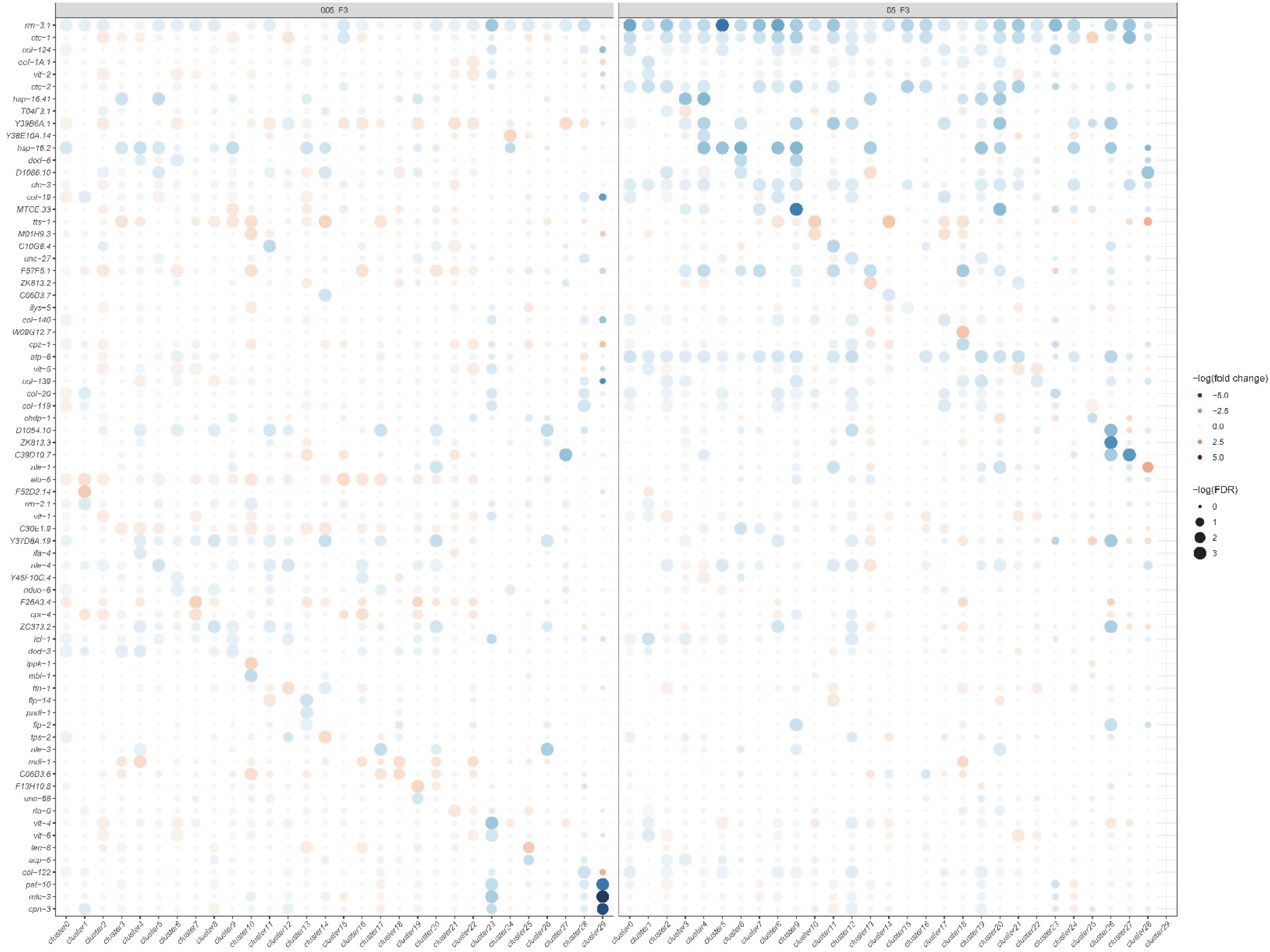
Dot heatmap of top 3 differential expressed genes across clusters at the F3. X-axis indicates different clusters representing different tissues and Y-axis indicates the top 3 differentially expressed genes after ethanol treatment across clusters ranked by monocle based FDR. The plot is further separated by different ethanol doses (0.05% on the left and 0.5% on the right). The size of the dot is correlated to –log(FDR) of differential expression p-value and the color represents direction and scale of fold change with upregulation is shown in red and downregulation is shown in blue.

