## Supplemental Information for "Single-nucleus resolution mapping of the adult *C. elegans* and its application to elucidate inter- and trans-generational response to alcohol"

**Supplemental material and methods:** Additional methodological information related to nematode culture, nuclei isolation, sequencing, and snRNA-seq data processing.

**Table S1:** Sequence of probes used for smFISH

**Table S2:** DEG list by cluster at the F1 from P0 0.05% ethanol exposure.

**Table S3:** DEG list by cluster at the F1 from P0 0.5% ethanol exposure.

**Table S4:** Significantly enriched Gene Ontology categories by generation and exposure condition.

**Table S5:** DEG list by cluster at the F3 from P0 0.05% ethanol exposure.

**Table S6:** DEG list by cluster at the F3 from P0 0.5% ethanol exposure.

**Supplemental data 1:** Dashboard including for each cluster: t-SNE, top 20 enriched and depleted genes, Tissue Enrichment Analysis, Gene Ontology, Phenotype Enrichment, and two examples of genes enriched in each cluster with corresponding *in situ* data from NEXTDB. In the top and bottom 20 genes, Pct.1 refers to the percentage of cells in the cluster expressing said genes and Pct. 2, the percentage of cells expressing this gene in all other clusters.

**Supplemental material and methods**

***C. elegans* exposures and expansions**

For ethanol exposures, a population of gravid adult worms was bleached. The embryos obtained were plated on standard OP50 seeded NGM plates and allowed to grow to the L4 stage (approximately 50 hours post bleaching). Nematodes were exposed for 48 hours in liquid culture containing M9 buffer solution, standard OP50 bacteria (10 mg/mL), and ethanol at a final concentration of 0.05% or 0.50% in 15mL conical tubes. Following the liquid exposure, the progeny of the exposed P0 generation were obtained using gravid adult bleaching. The synchronized F1 egg population was then plated on multiple standard OP50-seeded NGM plates. Half of the plates were grown for 16 hours at 20°C at which time all F1 were at the L1 stage and these plates were then transferred to 25°C for 48 hours into adulthood before proceeding with single-nucleus dissociation of the F1 generation.

The other half of the plates were grown at 20°C for 4 days or until the plates were filled with F2 L1 larvae. This population was synchronized using a 10μm nylon mesh filter (EDM Millipore NY1102500 and EDM Millipore SX00025000) which only allowed L1 staged worms to pass. The L1 worms were washed twice with M9 and centrifuged at 100g for 1 minute. These L1 worm pellets were plated on fresh OP50-seeded NGM plates and grown at 20°C for 4 days or until the plates were filled with F3 L1 larvae. This population was synchronized using a 10μm nylon mesh filter similar to the F2 population and the F3 L1 plates were transferred to 25°C. Worms were grown for 48 hours at 25°C before proceeding with single-nucleus dissociation of the F3 generation.

**Single-nucleus dissociation**

All reagents except for Triton-X100, PBS-BSA 1%, and inhibitors were prepared the night before and all low retention 1.5mL microcentrifuge tubes were clearly labeled. The bench space and equipment used in this protocol were thoroughly sanitized with 70% ethanol and cleaned with RNaseZAP (Thermo Fisher AM9780) before starting the dissociation. Reagents were prepared using RNase free water (Thermo Fisher BP2484100). The FA lysis buffer was made using the following reagents: 50mM HEPES/NaOH pH 7.5, 1mM EDTA, 0.1% Triton X-100, 150mM NaCl, Protease inhibitor 0.5X (Roche 11697498001), RNase inhibitor 0.2U/μL (Thermo Fisher 10777019), and RNase free water and stored at 4°C or on ice. BSA was prepared to a final concentration of 1% in pH 7.4 1X PBS (Thermo Fisher AM9624) using RNAse free water and RNAse free PBS. This solution was filtered using a 0.22μm pressure filter (Thermo Fisher 03-377-26, Thermo Fisher SLGP033RS).

All equipment and reagents were moved to a 4°C cold room and subsequent steps were performed at 4°C. Homogenizers were stored pre-chilled at -20°C when not in use and moved to the 4°C room before starting the extraction. Each Wheaton 1.5mL Dounce homogenizers (Sigma Z378623-1EA) was cleaned using 70% ethanol, RNaseZAP, and RNase free water. Homogenizers were rinsed twice with ethanol, twice with RNaseZAP, and 5 times with 1-2mL of RNase free water.

For both F1 and F3 generations, L1 larvae were grown at 25°C for 48 hours. Adult worms were gently washed off plates with M9 and transferred into 15 mL conical tubes, being careful not to disrupt bacterial lawn. Worms were allowed to settle to the bottom of the conical tube by gravity for 5 minutes before transferring the worm pellet to a sterilized 1.5mL low-bind microcentrifuge tube. The worm pellet was then washed 5 times with M9, centrifuging the tubes at 1,300g for 1 minute in between each wash in order to remove bacteria. After washing, worms were placed in 1mL of M9 in a 1.5mL low-bind microcentrifuge tube and incubated in a rotator at 20°C for 30 minutes to remove residual OP50 from the worms’ gut. These microcentrifuge tubes were then set upright and the worms were allowed to settle by gravity for 5 minutes. The M9 supernatant was discarded and the final compact worm pellet volume was adjusted to 30μL.

The compact 30μL pellet of adult *C. elegans* was transferred to the Dounce homogenizer and 400μL of ice-cold FA buffer was used to rinse any remaining worms from the 1.5mL low bind microcentrifuge tube and added to the homogenizer. Worms were homogenized with 10 strokes of the Dounce homogenizer using a corkscrew motion with a B (tight) pestle. Homogenized worms were transferred to a new low bind 1.5ml microcentrifuge tube and centrifuged at 100g for 1 minute to pellet debris. The supernatant containing the dissociated nuclei was removed using a 1000μL low bind tip and transferred to a fresh low bind 1.5mL microcentrifuge tube labeled pooled nuclei. 300μL of FA buffer was added to the debris remaining in the first microcentrifuge tube and homogenized using 10 strokes in a corkscrew fashion with an Eppendorf Dounce homogenizer. The newly homogenized sample was then centrifuged at 100g for 1 minute to pellet debris. The supernatant containing the newly dissociated nuclei was pooled with the previously dissociated nuclei and the previous steps with the Eppendorf Dounce homogenizer were repeated once more to further homogenize the sample. In total, worms were homogenized with 30 strokes: 10 strokes with the 1.5mL Wheaton Dounce homogenizer and 20 strokes with the Eppendorf Dounce homogenizer. Between each homogenization step, debris was pelleted at 100g for 1 minute and the supernatant containing the dissociated nuclei was removed and added to a single 1.5mL microcentrifuge tube labeled pooled nuclei. Dissociated nuclei were removed after each set of 10 homogenization strokes to prevent over digestion of nuclei.

After homogenization, the pooled supernatant containing the dissociated nuclei was centrifuged at 100g for 1 minute to pellet any remaining or accidentally transferred debris. The top 900μL of supernatant containing nuclei was transferred to a clean low bind 1.5mL microcentrifuge tube, being careful not to disturb the debris pellet. These pooled nuclei were pelleted at 500g for 4 minutes. After pelleting, approximately 800μL of FA buffer was removed, being careful not to disrupt the nuclei pellet, and the pelleted nuclei were resuspended with 1000μL of 1% PBS-BSA. The nuclei were again centrifuged at 500g for 4 minutes and 1000μL of the 1% PBS-BSA supernatant was removed. Lastly, the nuclei pellet was resuspended in 750-850μL of 1% PBS-BSA (final volume was determined by examining the size of the nuclei pellet). After resuspension, the nuclei were filtered using a 40μm Flowmi tip filter (Sigma Aldrich BAH136800040-50EA). Filtered nuclei were transferred to a 1.5mL low retention microcentrifuge tube for FACS sorting or 10X sequencing.

**FACS/FLOW**

The BD Analyzer Celesta plate reader at the UCLA BSCRC flow cytometry core was used to assess nuclei concentration. 150μL aliquots of filtered nuclei samples were stained with DAPI to determine concentration. Flow cytometry was done using the Violet 405nm 50mW laser with the slowest flow rate to obtain accurate counts. Nuclei concentration was determined to be within 700 to 1200 nuclei per microliter. If concentration was too high, filtered nuclei sample was diluted with 1% PBS-BSA. A flat bottom clear 96-well plate was used to assess nuclei concentration.

**Library Preparation and sequencing**

Library preparation was performed by UCLA Technology Center for Genomics & Bioinformatics. Nuclei were isolated into single droplets and barcoded using the 10X Chromium Next GEM single cell 3ʹ reagent kit. We sequenced using 50bp long paired end reads with the NovaSeq 6000.

**Single-nuclei transcriptional analysis**

snRNA-seq reads were demultiplexed and aligned to the ENSEMBL ce10 *C. elegans* transcriptome to generate gene expression matrices using CellRanger (10x Genomics). The reference transcriptome was converted to accommodate pre-mRNA alignment by replacing “transcript” to “exon” in annotation GTF file. We first filtered the matrices to exclude low-quality cells or potential doublets using the following criteria: 1) gene number less than 300 or more than 8000, 2) unique molecular identifier (UMI) count less than 00 or more than 40000, 3) mitochondrial RNA percentage > 15% per cell, and 4) ribosomal RNA >20% per cell. After preprocessing, 4694, 16148, 11738, 9169 cells were retained in unexposed, water treatment,0.05% and 0.5% ethanol treatment groups, respectively.

**Identification of cell clusters**

R Seurat 3.1.5 [1] package was used for normalization, cell type identification, marker identification and batch effect correction of snRNA-seq data using all 31 sample groups. snRNA-seq data was log-normalized. The top 2,000 variable genes were selected as representative features, followed by correcting gene expression with UMI counts, mitochondrial gene percentage and ribosomal RNA percentage for further clustering analysis. Canonical correlation analysis (CCA) was applied across different batches and treatment conditions to mitigate batch effects in cluster identification. Cell clusters were identified from Louvain algorithm [2]. We included all treatment groups for unsupervised clustering since increased cell numbers was shown to increase power in identifying smaller cell types [3]. Cluster specific genes were detected by Wilcoxon Rank Sum test [4]. To reduce biases from treatment in finding markers, only unexposed cells were included unless unexposed groups consist of less than 20% of the cluster of interest. Furthermore, for each cluster, the gene had to be expressed in at least 25% of the cells of the given cluster and there had to be at least a 0.25 log fold change in gene expression compared to other cells. Log-normalized expression levels in t-SNE (t-distributed stochastic neighbor embedding) plot projections were used to visualize cell clusters in two dimensions and dot heatmap were used to visualize marker expression across different cell types. While tSNE clusters were created using all 31 samples, marker genes enriched for each cluster were identified using only the unexposed samples to avoid confounding effects of ethanol.

**Differential gene expression and pathway analyses**

Monocle [5] pipeline was used in order to identify DEGs across different cell types, generations and dose levels. Four different monocle models were created to assess DEGs in F1_0.05, F1_0.5, F3_0.05 and F3_0.5 condition. For each condition (generation and dose level), only cell types with more than 10 cells in each group were included. For genes expressed in more than 20% of cells in each cell type, a negative binomial model was fitted based on raw counts to normalize data, followed by fitting a generalized linear model to retrieve dietary exposure effect with batch effects corrected as follows:

Gene expression = b1*batch+b2*ethanol+b3*gene count+b4*UMI count

Batch term is only included in F1_0.05 and F3_0.05 where two batches of water and ethanol 0.05% samples were produced, for F1_0.5 and F3_0.5 condition this term is not used since only water and ethanol 0.5% samples from the same batch were considered. The b2 coefficient obtained will be used to estimate dietary exposure effects. Statistical p-value was obtained using a likelihood ratio test against the null model where the exposure term is not included. Significant DEGs were defined as genes with Benjamini & Hochberg corrected FDR < 0.05 [6].

The DEGs were then subject to pathway annotation analysis. Only cell types with no less than 20 DEGs were included in this analysis. Gene ontology analysis was conducted using clusterprofiler package [7] with *C. elegans* gene ontology biological pathway (GOBP), molecular function (GOMF) database [8] and wormbase phenotype database [9]. Enrichment P values were corrected by Benjamini–Hochberg method and FDR < 0.05 were considered significant, only pathways with more than 2 overlapped genes were kept. For significantly enriched pathways, fold changes were calculated by averaging the fold changes of the pathway genes between treatment and control nuclei. For WormBase phenotypes, we also retrieved higher level categories of each phenotype by querying EBI OLS (ontology lookup service) API. Annotations from first level (nematode phenotype, physiology phenotype and anatomical phenotype) since these terms were too general to make interpretations. We further selected top 20 most common annotations and compared their proportion in original database with our enrichment results.

**Euclidean distance-based measurement of cell type sensitivity**

To identify cell types that are sensitive to ethanol treatment, the Euclidean distance metric was used [10]. For each cell type with more than 10 cells in both ethanol and control group per batch, expression distance between nuclei of water and ethanol treatment groups were squared and summed, followed by taking the square root. In order to avoid potential biases caused by genes that are either highly expressed or non-expressed, expression values were normalized to z-scores and only the top 1,000 expressed genes were used. To account for variabilities in expression characteristics per each cell type, null distributions for individual cell types were calculated based on permutated treatment labels for 1,000 times. P values were calculated between the observed Euclidean distance and the null distribution for each cell type and adjusted with the Benjamini & Hochberg method [6].

To visualize the differences between water and ethanol treated nuclei for individual cell types, the fold change (FC) in the Euclidean distance of ethanol treatment group compared with water treatment group in each cell type was normalized by dividing the empirical Euclidean distance by the median Euclidean distance of the null distribution per cell type. The log10(FC) vs. -log10(adjusted p value) of each cell type was then plotted to visualize and rank the vulnerable cell types in ethanol treatment. For 0.05% where two batches were generated, FDR and log10(FC) were averaged.

**Statistical Analysis**

Unless otherwise mentioned, statistical analyses were conducted by R/3.5.1.

**Supplemental References.**
