## Supplemental File 1_Dashboard for "Single-nucleus resolution mapping of the adult *C. elegans* and its application to elucidate inter- and trans-generational response to alcohol"

### t-SNE showing cell type 0

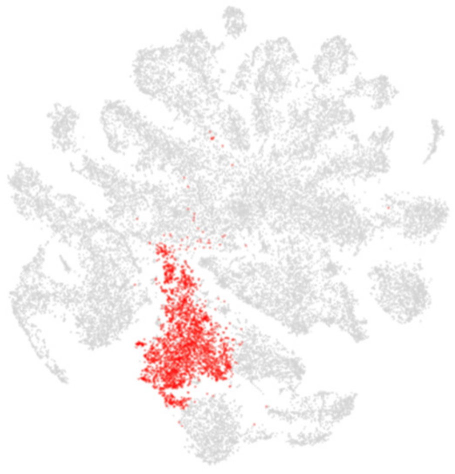

### Top 20 enriched genes

| | p_val | avg_log2FC | pct.1 | pct.2 | p_val_adj |
|---|---|---|---|---|---|
| *vit-3* | 0 | 2.40753077 | 0.97 | 0.69 | 0 |
| *F57F5.1* | 0 | 2.36625453 | 0.945 | 0.461 | 0 |
| *vit-2* | 0 | 2.35693413 | 0.987 | 0.701 | 0 |
| *vit-5* | 0 | 2.33849234 | 0.981 | 0.745 | 0 |
| *H34I24.2* | 0 | 2.32123583 | 0.717 | 0.27 | 0 |
| *vit-1* | 0 | 2.23320352 | 0.954 | 0.651 | 0 |
| *lipl-5* | 0 | 2.20121706 | 0.939 | 0.567 | 0 |
| *vit-4* | 0 | 2.1749582 | 0.886 | 0.664 | 0 |
| *spp-17* | 0 | 2.12359357 | 0.843 | 0.41 | 0 |
| *fat-2* | 0 | 2.09367012 | 0.897 | 0.451 | 0 |
| *vit-6* | 0 | 2.08227724 | 0.984 | 0.735 | 0 |
| *lys-2* | 0 | 2.05854736 | 0.875 | 0.439 | 0 |
| *cpr-6* | 0 | 2.01385847 | 0.974 | 0.654 | 0 |
| *asp-3* | 0 | 2.01285413 | 0.961 | 0.56 | 0 |
| *asah-1* | 0 | 2.00459586 | 0.678 | 0.231 | 0 |
| *K12H4.7* | 0 | 1.96435666 | 0.879 | 0.442 | 0 |
| *dct-16* | 0 | 1.94856152 | 0.992 | 0.742 | 0 |
| *Y51F10.7* | 0 | 1.8522103 | 0.745 | 0.344 | 0 |
| *F28B4.3* | 0 | 1.83235041 | 0.691 | 0.29 | 0 |
| *cpr-1* | 0 | 1.83163546 | 0.866 | 0.417 | 0 |

### Top 20 depleted genes

| | p_val | avg_log2FC | pct.1 | pct.2 | p_val_adj |
|---|---|---|---|---|---|
| *dig-1* | 5.96E-62 | -3.3888185 | 0.071 | 0.251 | 2.80E-57 |
| *D1086.10* | 1.45E-61 | -2.8996673 | 0.22 | 0.407 | 6.78E-57 |
| *ule-4* | 1.87E-50 | -2.8347517 | 0.376 | 0.518 | 8.76E-46 |
| *ule-3* | 1.11E-20 | -2.5996429 | 0.157 | 0.249 | 5.21E-16 |
| *ttn-1* | 5.59E-125 | -2.5955658 | 0.203 | 0.492 | 2.62E-120 |
| *ZC513.7* | 1.05E-109 | -2.439556 | 0.181 | 0.45 | 4.94E-105 |
| *D1054.10* | 9.83E-15 | -2.3070872 | 0.251 | 0.327 | 4.61E-10 |
| *act-4* | 2.03E-254 | -2.2836538 | 0.421 | 0.744 | 9.54E-250 |
| *ZK813.7* | 1.00E-18 | -2.2766945 | 0.143 | 0.23 | 4.70E-14 |
| *unc-54* | 2.14E-220 | -2.2620756 | 0.38 | 0.709 | 1.00E-215 |
| *far-2* | 1.19E-231 | -2.2327528 | 0.555 | 0.784 | 5.60E-227 |
| *pat-10* | 1.21E-196 | -2.1994722 | 0.481 | 0.726 | 5.66E-192 |
| *ule-2* | 2.64E-33 | -2.1939963 | 0.175 | 0.304 | 1.24E-28 |
| *fasn-1* | 1.19E-45 | -2.1630796 | 0.133 | 0.288 | 5.59E-41 |
| *ttr-16* | 3.83E-170 | -2.1557018 | 0.403 | 0.665 | 1.80E-165 |
| *mlc-3* | 1.34E-200 | -2.133881 | 0.425 | 0.704 | 6.29E-196 |
| *C10G8.4* | 1.38E-22 | -2.128373 | 0.252 | 0.351 | 6.47E-18 |
| *Y62H9A.5* | 1.13E-14 | -2.1111424 | 0.211 | 0.287 | 5.32E-10 |
| *lev-11* | 4.50E-229 | -2.1049562 | 0.454 | 0.75 | 2.11E-224 |
| *cpn-3* | 5.60E-180 | -2.0833823 | 0.449 | 0.701 | 2.63E-175 |

### Tissue enrichment

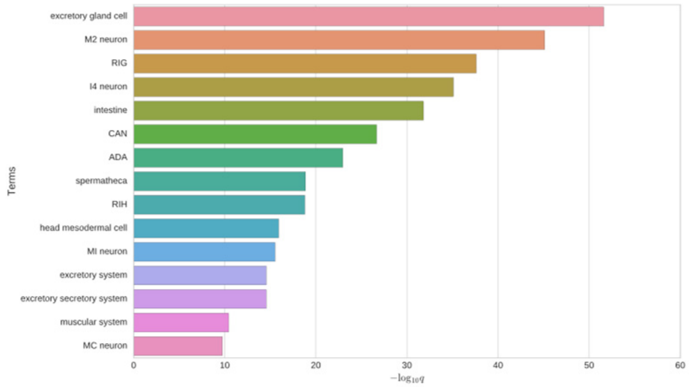


### GO enrichment

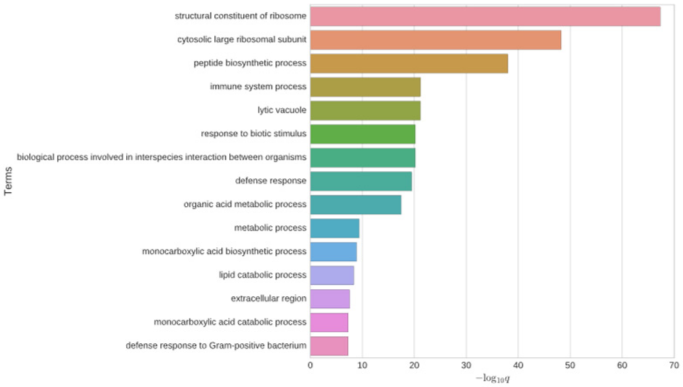


### Phenotype enrichment

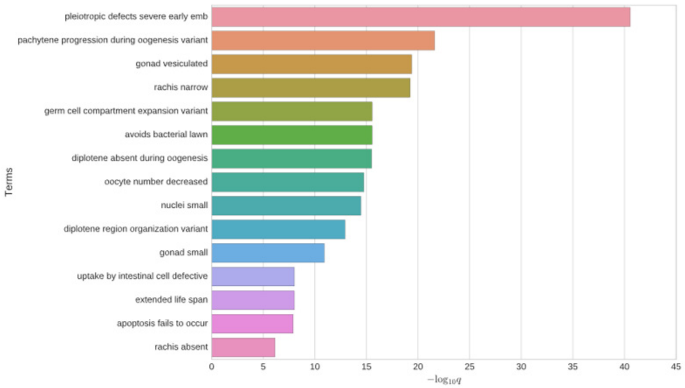


### *asah-1*

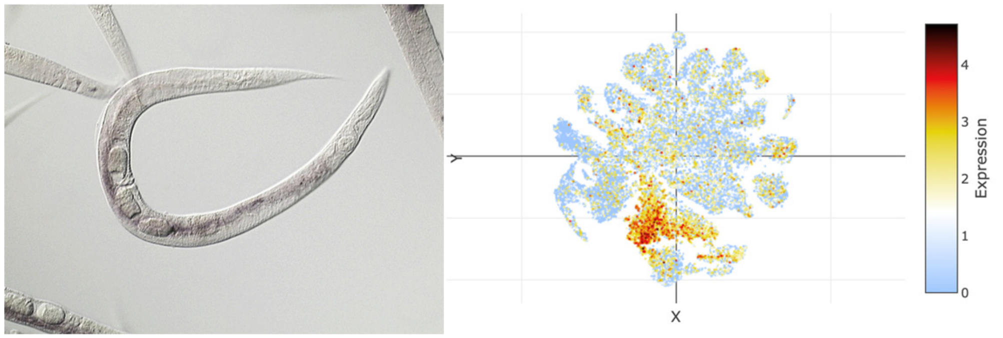


## *H34I24.2*

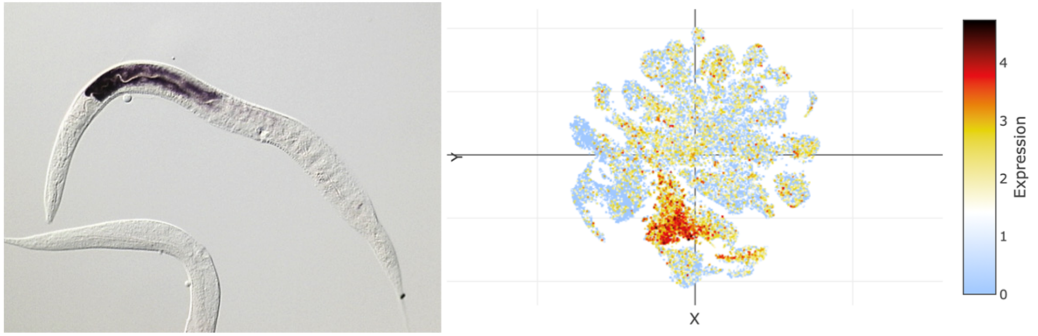

### t-SNE showing cell type 1

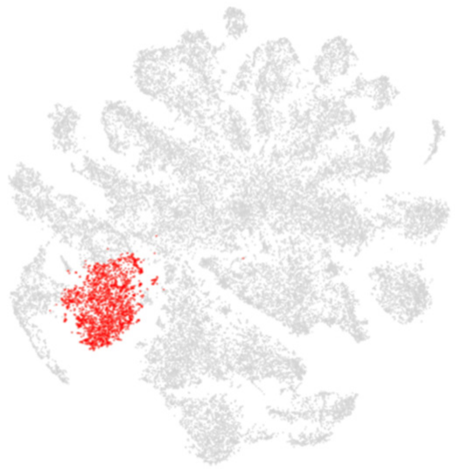

### Top 20 enriched genes

| | p_val | avg_log2FC | pct.1 | pct.2 | p_val_adj |
|---|---|---|---|---|---|
| *linc-7* | 2.79E-277 | 2.8461492 | 0.564 | 0.188 | 1.31E-272 |
| *mix-1* | 0 | 2.64656064 | 0.79 | 0.264 | 0 |
| *set-24* | 0 | 2.64356758 | 0.552 | 0.08 | 0 |
| *Y57A10A.31* | 0 | 2.59424002 | 0.593 | 0.132 | 0 |
| *such-1* | 0 | 2.4308028 | 0.533 | 0.104 | 0 |
| *Y57A10A.1* | 0 | 2.42585518 | 0.478 | 0.051 | 0 |
| *xpc-1* | 0 | 2.42568406 | 0.648 | 0.187 | 0 |
| *mdt-17* | 0 | 2.4250593 | 0.466 | 0.096 | 0 |
| *H06I04.3* | 0 | 2.41221422 | 0.683 | 0.237 | 0 |
| *F52D2.14* | 4.74E-224 | 2.40706998 | 0.253 | 0.041 | 2.22E-219 |
| *fcd-2* | 0 | 2.36277275 | 0.555 | 0.121 | 0 |
| *npp-8* | 0 | 2.36057674 | 0.642 | 0.21 | 0 |
| *T05F1.2* | 2.97E-245 | 2.3525074 | 0.619 | 0.257 | 1.39E-240 |
| *chk-2* | 0 | 2.33130698 | 0.39 | 0.028 | 0 |
| *lea-1* | 0 | 2.31164841 | 0.877 | 0.531 | 0 |
| *gcn-1* | 1.62E-286 | 2.30627886 | 0.645 | 0.264 | 7.60E-282 |
| *F53H1.4* | 0 | 2.241348 | 0.645 | 0.206 | 0 |
| *Y48G8AL.5* | 0 | 2.23551318 | 0.541 | 0.131 | 0 |
| *Y79H2A.3* | 0 | 2.20657813 | 0.631 | 0.214 | 0 |
| *attf-6* | 0 | 2.20258992 | 0.511 | 0.135 | 0 |

### Top 20 depleted genes

| | p_val | avg_log2FC | pct.1 | pct.2 | p_val_adj |
|---|---|---|---|---|---|
| *ule-4* | 1.52E-67 | -3.32952 | 0.285 | 0.52 | 7.15E-63 |
| *D1086.10* | 4.67E-54 | -3.1766948 | 0.191 | 0.404 | 2.19E-49 |
| *pat-10* | 1.34E-236 | -3.1255073 | 0.28 | 0.731 | 6.29E-232 |
| *lev-11* | 1.23E-263 | -3.0455223 | 0.27 | 0.753 | 5.75E-259 |
| *act-4* | 6.72E-265 | -3.0262073 | 0.242 | 0.746 | 3.15E-260 |
| *far-2* | 1.18E-250 | -3.016277 | 0.392 | 0.787 | 5.55E-246 |
| *dig-1* | 1.67E-33 | -2.9673044 | 0.088 | 0.246 | 7.84E-29 |
| *cpn-3* | 6.78E-221 | -2.9104893 | 0.254 | 0.706 | 3.18E-216 |
| *mlc-3* | 1.43E-233 | -2.8939846 | 0.235 | 0.708 | 6.69E-229 |
| *ttr-16* | 3.64E-214 | -2.8665463 | 0.199 | 0.669 | 1.71E-209 |
| *unc-54* | 5.17E-218 | -2.8490946 | 0.257 | 0.707 | 2.42E-213 |
| *lbp-2* | 2.38E-190 | -2.8031531 | 0.225 | 0.655 | 1.11E-185 |
| *unc-27* | 1.07E-210 | -2.7971513 | 0.24 | 0.69 | 5.03E-206 |
| *F46H5.3* | 1.07E-301 | -2.7703362 | 0.34 | 0.827 | 5.01E-297 |
| *nlp-77* | 2.54E-214 | -2.7427856 | 0.304 | 0.726 | 1.19E-209 |
| *unc-87* | 6.95E-217 | -2.7392701 | 0.171 | 0.654 | 3.26E-212 |
| *unc-15* | 1.51E-212 | -2.7286407 | 0.213 | 0.671 | 7.08E-208 |
| *col-95* | 2.76E-155 | -2.7143854 | 0.212 | 0.61 | 1.30E-150 |
| *perm-4* | 1.63E-120 | -2.6835737 | 0.257 | 0.597 | 7.65E-116 |
| *col-140* | 6.81E-223 | -2.5844856 | 0.365 | 0.758 | 3.19E-218 |

### Tissue enrichment

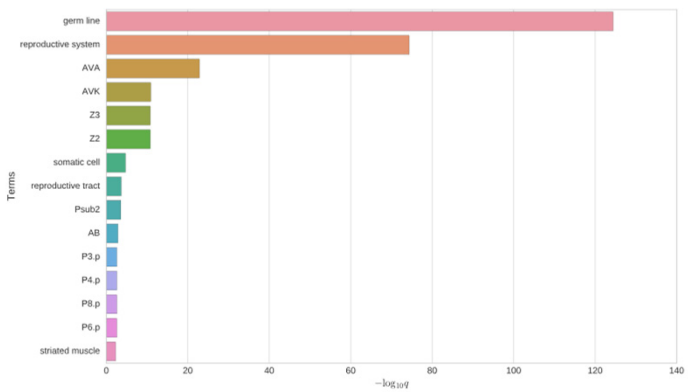


### GO enrichment

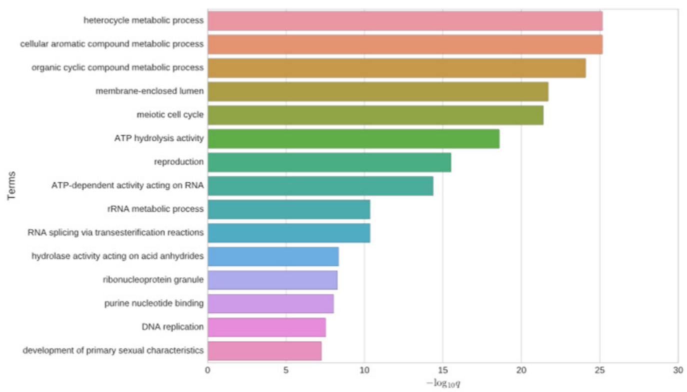


### Phenotype enrichment

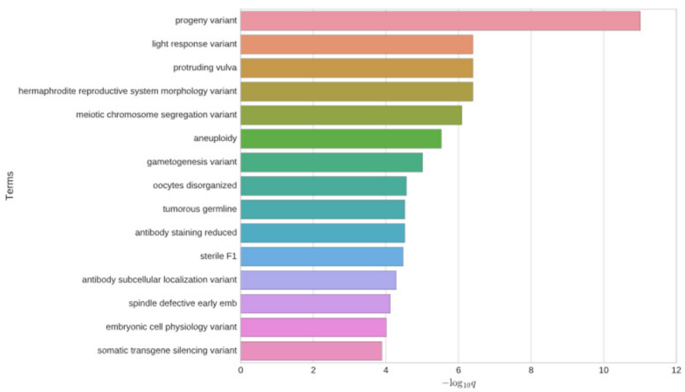


### *chk-2*

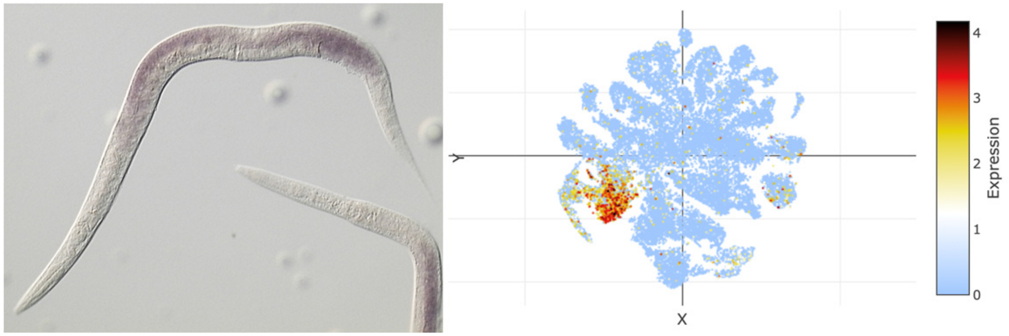


### *such-1*

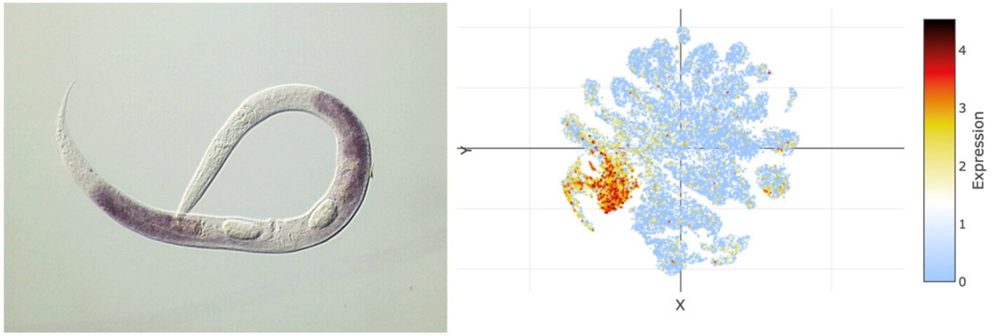

### t-SNE showing cell type 2

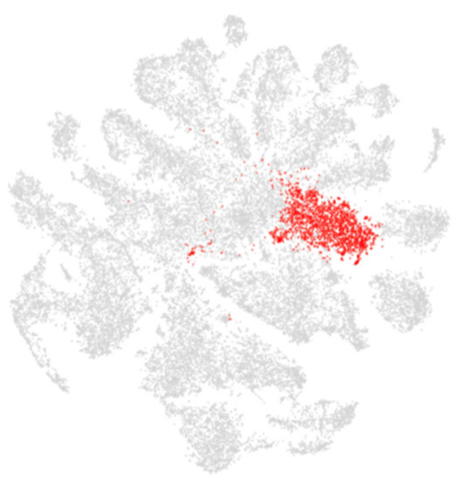

### Top 20 enriched genes

| | p_val | avg_log2FC | pct.1 | pct.2 | p_val_adj |
|---|---|---|---|---|---|
| *unc-27* | 0 | 2.51261312 | 0.992 | 0.647 | 0 |
| *unc-54* | 0 | 2.35277309 | 0.997 | 0.665 | 0 |
| *F37H8.5* | 0 | 2.35062914 | 0.911 | 0.428 | 0 |
| *tnt-2* | 0 | 2.33913036 | 0.981 | 0.58 | 0 |
| *lbp-2* | 0 | 2.30163363 | 0.982 | 0.611 | 0 |
| *cpn-3* | 0 | 2.23614211 | 0.992 | 0.663 | 0 |
| *unc-15* | 0 | 2.21255302 | 0.988 | 0.626 | 0 |
| *unc-87* | 0 | 2.2004755 | 0.989 | 0.607 | 0 |
| *clik-1* | 0 | 2.19257175 | 0.986 | 0.658 | 0 |
| *ttr-25* | 0 | 2.18420317 | 0.84 | 0.384 | 0 |
| *ttr-16* | 0 | 2.1791393 | 0.992 | 0.624 | 0 |
| *pat-10* | 0 | 2.15411398 | 0.997 | 0.69 | 0 |
| *lec-5* | 0 | 2.13330611 | 0.89 | 0.439 | 0 |
| *act-1* | 0 | 2.11696419 | 0.999 | 0.76 | 0 |
| *R13H4.2* | 0 | 2.11351016 | 0.883 | 0.407 | 0 |
| *mup-2* | 0 | 2.08212893 | 0.943 | 0.538 | 0 |
| *lev-11* | 0 | 2.06259183 | 0.997 | 0.711 | 0 |
| *myo-3* | 0 | 2.05768003 | 0.87 | 0.413 | 0 |
| *act-4* | 0 | 2.05004295 | 0.996 | 0.702 | 0 |
| *ttr-24* | 0 | 2.0464749 | 0.869 | 0.398 | 0 |

### Top 20 depleted genes

| | p_val | avg_log2FC | pct.1 | pct.2 | p_val_adj |
|---|---|---|---|---|---|
| *ule-3* | 5.58E-18 | -2.7243192 | 0.146 | 0.248 | 2.62E-13 |
| *D1054.10* | 4.77E-25 | -2.6661453 | 0.197 | 0.328 | 2.24E-20 |
| *ule-4* | 5.11E-14 | -2.5256237 | 0.449 | 0.51 | 2.40E-09 |
| *D1086.10* | 6.78E-11 | -2.5136821 | 0.334 | 0.396 | 3.18E-06 |
| *ZK813.7* | 7.39E-15 | -2.4653669 | 0.14 | 0.228 | 3.47E-10 |
| *Y37D8A.19* | 1.26E-25 | -2.4334875 | 0.232 | 0.364 | 5.92E-21 |
| *Y62H9A.5* | 1.06E-21 | -2.4331092 | 0.169 | 0.287 | 4.98E-17 |
| *Y46H3C.7* | 4.76E-12 | -2.3137245 | 0.06 | 0.127 | 2.23E-07 |
| *ule-5* | 8.38E-20 | -2.3087491 | 0.223 | 0.333 | 3.93E-15 |
| *ZK813.3* | 9.36E-17 | -2.2143448 | 0.146 | 0.246 | 4.39E-12 |
| *ZK813.1* | 9.74E-16 | -2.2066885 | 0.119 | 0.209 | 4.57E-11 |
| *ZC373.2* | 1.85E-19 | -2.1305956 | 0.228 | 0.339 | 8.66E-15 |
| *C39D10.7* | 6.15E-16 | -2.0528625 | 0.192 | 0.288 | 2.89E-11 |
| *ZK813.2* | 2.64E-20 | -2.0403526 | 0.187 | 0.295 | 1.24E-15 |
| *C10G8.4* | 3.34E-11 | -2.0156747 | 0.273 | 0.347 | 1.57E-06 |
| *F55B11.2* | 1.24E-22 | -1.8482166 | 0.097 | 0.212 | 5.80E-18 |
| *C30E1.9* | 2.24E-43 | -1.8151273 | 0.313 | 0.493 | 1.05E-38 |
| *W02D9.7* | 1.82E-16 | -1.7365146 | 0.149 | 0.246 | 8.55E-12 |
| *fipr-2* | 5.14E-15 | -1.7134813 | 0.111 | 0.2 | 2.41E-10 |
| *T21C9.13* | 1.46E-21 | -1.7054615 | 0.056 | 0.159 | 6.83E-17 |

### Tissue enrichment

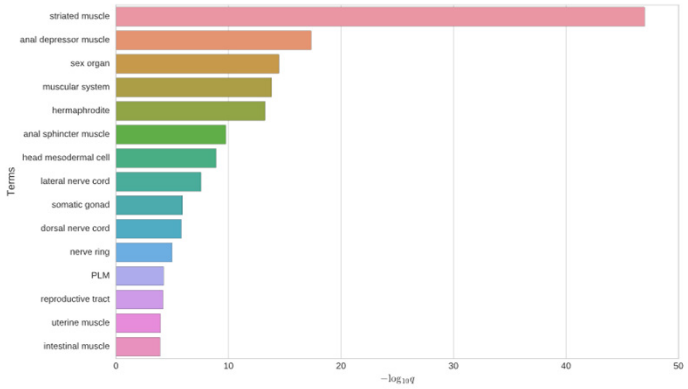


### GO enrichment

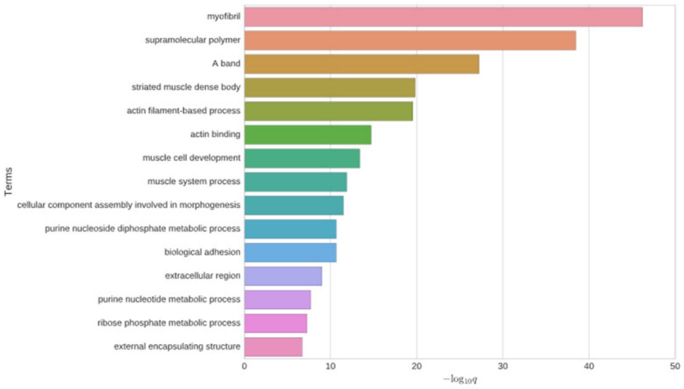


### Phenotype enrichment

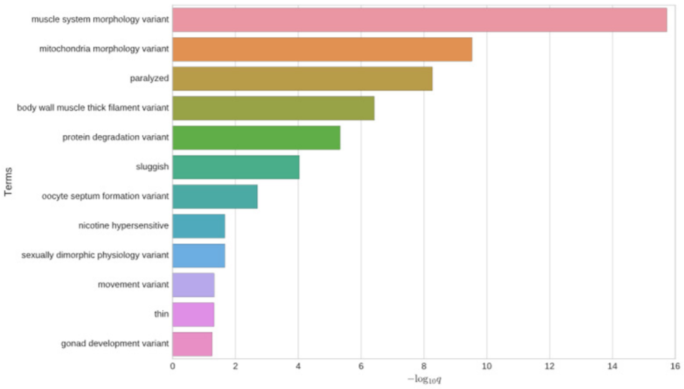


### *tni-1*

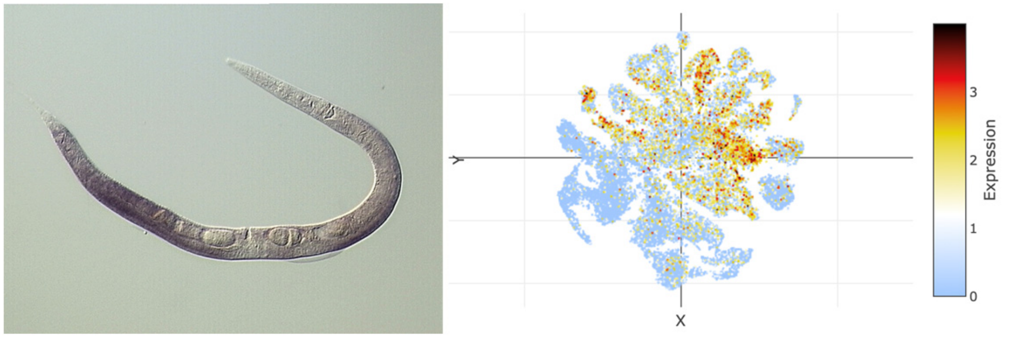


### *myo-3*

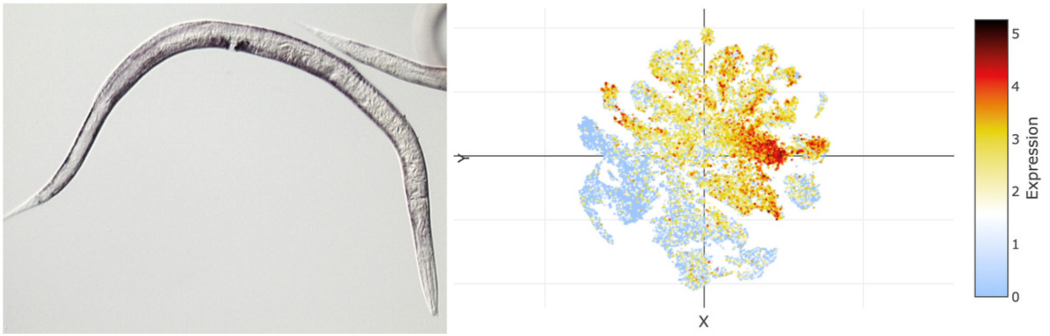

### t-SNE showing cell type 3

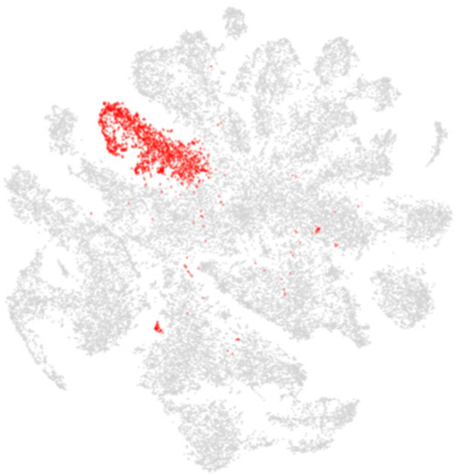

### Top 20 enriched genes

| | p_val | avg_log2FC | pct.1 | pct.2 | p_val_adj |
|---|---|---|---|---|---|
| *ule-3* | 0 | 5.53196667 | 0.943 | 0.2 | 0 |
| *ZK813.7* | 0 | 5.35917056 | 0.917 | 0.182 | 0 |
| *D1054.10* | 0 | 5.24724634 | 0.987 | 0.281 | 0 |
| *ZK813.3* | 0 | 5.11824728 | 0.944 | 0.199 | 0 |
| *ZK813.1* | 0 | 5.11695215 | 0.901 | 0.163 | 0 |
| *Y62H9A.5* | 0 | 5.00096792 | 0.955 | 0.241 | 0 |
| *Y37D8A.19* | 0 | 4.96385168 | 0.985 | 0.32 | 0 |
| *ule-5* | 0 | 4.88593002 | 0.978 | 0.289 | 0 |
| *ZC373.2* | 0 | 4.82725502 | 0.982 | 0.294 | 0 |
| *Y57G11B.5* | 0 | 4.6462841 | 0.876 | 0.139 | 0 |
| *F55B11.2* | 0 | 4.6248496 | 0.883 | 0.166 | 0 |
| *F55B11.3* | 0 | 4.53398409 | 0.846 | 0.12 | 0 |
| *W02D9.7* | 0 | 4.50938141 | 0.889 | 0.202 | 0 |
| *F57C2.4* | 0 | 4.46009522 | 0.938 | 0.201 | 0 |
| *Y62H9A.4* | 0 | 4.26011605 | 0.874 | 0.156 | 0 |
| *Y62H9A.3* | 0 | 4.23553124 | 0.81 | 0.114 | 0 |
| *F53H4.2* | 0 | 3.94550278 | 0.78 | 0.14 | 0 |
| *F17E9.4* | 0 | 3.68168104 | 0.816 | 0.147 | 0 |
| *D1086.11* | 0 | 3.3350133 | 0.567 | 0.047 | 0 |
| *W02D9.6* | 0 | 3.31209428 | 0.607 | 0.072 | 0 |

### Top 20 depleted genes

| | p_val | avg_log2FC | pct.1 | pct.2 | p_val_adj |
|---|---|---|---|---|---|
| *dig-1* | 3.51E-10 | -2.9252217 | 0.168 | 0.241 | 1.64E-05 |
| *ttn-1* | 8.26E-13 | -1.4457523 | 0.399 | 0.473 | 3.87E-08 |
| *spsb-1* | 9.84E-15 | -1.2982337 | 0.203 | 0.293 | 4.61E-10 |
| *crh-1* | 3.92E-15 | -1.1897702 | 0.179 | 0.267 | 1.84E-10 |
| *F53H2.3* | 4.96E-11 | -1.173763 | 0.135 | 0.205 | 2.33E-06 |
| *lys-7* | 2.64E-08 | -1.1636346 | 0.269 | 0.33 | 0.00123679 |
| *ddo-2* | 8.82E-10 | -1.1633019 | 0.254 | 0.322 | 4.13E-05 |
| *Y105E8A.25* | 4.02E-15 | -1.1468382 | 0.263 | 0.35 | 1.88E-10 |
| *sos-1* | 5.14E-17 | -1.0759648 | 0.143 | 0.237 | 2.41E-12 |
| *mix-1* | 7.93E-14 | -1.0676817 | 0.216 | 0.297 | 3.72E-09 |
| *ced-1* | 1.79E-23 | -1.0378924 | 0.258 | 0.381 | 8.41E-19 |
| *tos-1* | 4.37E-19 | -1.0219256 | 0.407 | 0.503 | 2.05E-14 |
| *daf-2* | 3.70E-15 | -1.0164055 | 0.119 | 0.205 | 1.73E-10 |
| *mnk-1* | 1.30E-11 | -0.9814214 | 0.137 | 0.21 | 6.11E-07 |
| *F53E4.1* | 9.30E-12 | -0.9698191 | 0.202 | 0.282 | 4.36E-07 |
| *unc-22* | 3.44E-10 | -0.9627772 | 0.276 | 0.344 | 1.61E-05 |
| *jun-1* | 7.95E-07 | -0.9303412 | 0.096 | 0.144 | 0.03727813 |
| *col-162* | 8.18E-07 | -0.9114224 | 0.073 | 0.118 | 0.03837363 |
| *elpc-2* | 2.85E-10 | -0.8936066 | 0.136 | 0.2 | 1.34E-05 |
| *C02E7.7* | 6.85E-07 | -0.8918021 | 0.063 | 0.108 | 0.03212916 |

### Tissue enrichment

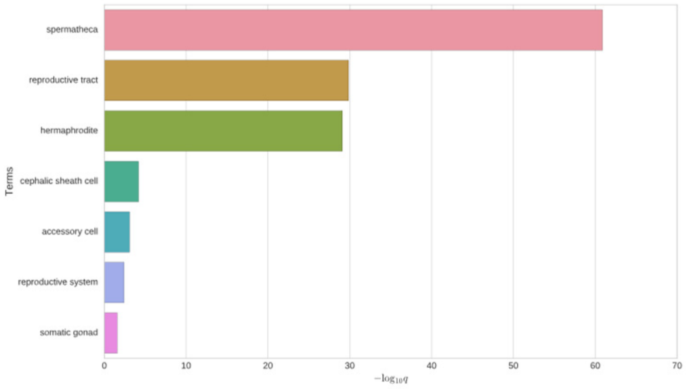


### GO enrichment

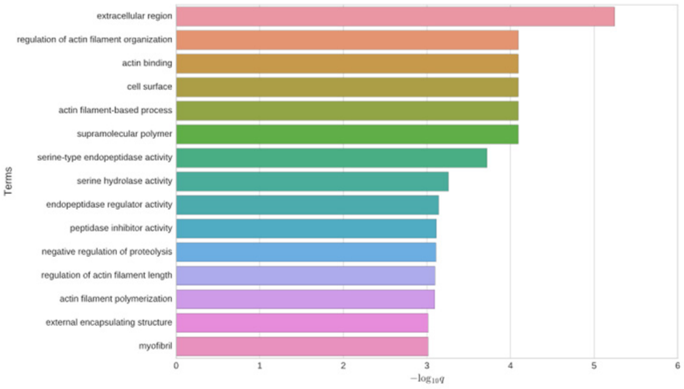


### Phenotype enrichment

**Nothing found on phenotype analysis**

### *ule-3*

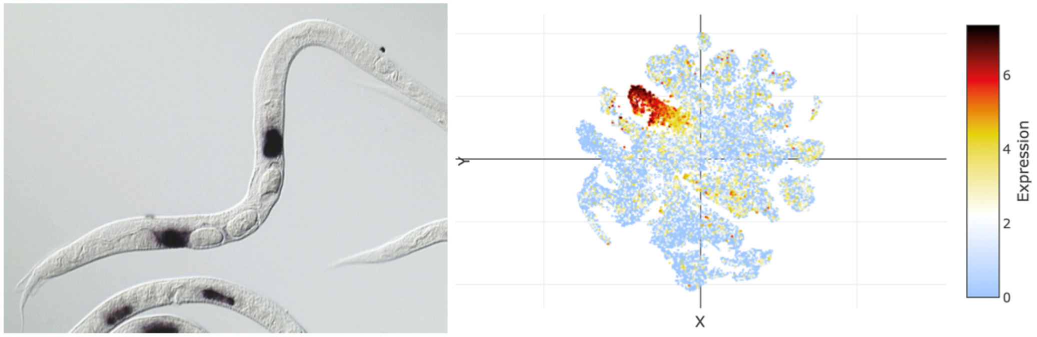


## *Y62H9A.5*

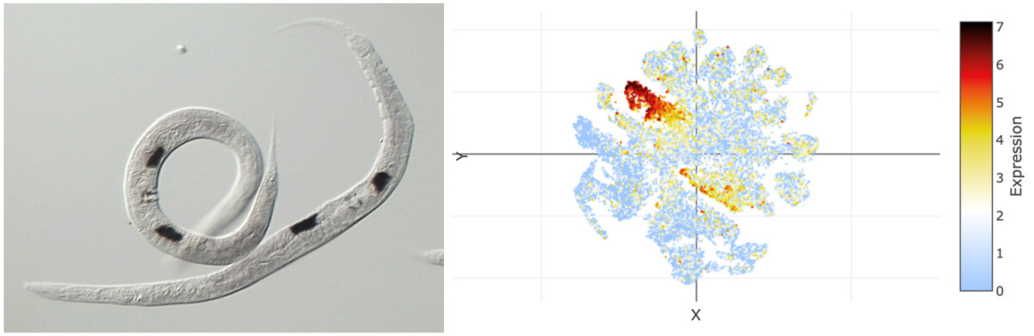

### t-SNE showing cell type 4

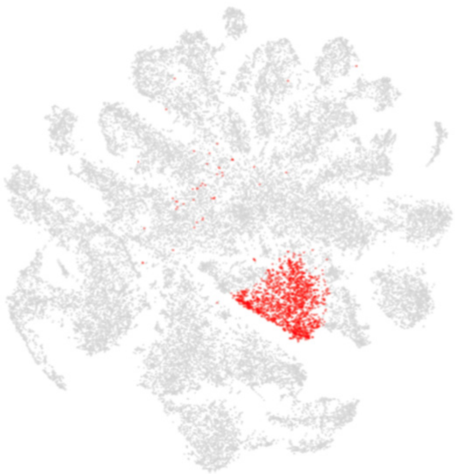

### Top 20 enriched genes

| | p_val | avg_log2FC | pct.1 | pct.2 | p_val_adj |
|---|---|---|---|---|---|
| *D1086.10* | 0 | 4.69156497 | 0.985 | 0.36 | 0 |
| *C35B1.4* | 0 | 4.23538542 | 0.836 | 0.229 | 0 |
| *Y22D7AR.10* | 0 | 4.08597529 | 0.927 | 0.259 | 0 |
| *ule-4* | 0 | 4.00805606 | 0.993 | 0.481 | 0 |
| *ZK813.2* | 0 | 3.86467499 | 0.896 | 0.256 | 0 |
| *Y45F10C.4* | 0 | 3.84203999 | 0.825 | 0.194 | 0 |
| *C10G8.4* | 0 | 3.71029704 | 0.897 | 0.313 | 0 |
| *F30A10.13* | 0 | 3.65922406 | 0.741 | 0.116 | 0 |
| *T12B5.15* | 0 | 3.61403938 | 0.7 | 0.161 | 0 |
| *Y43B11AR.1* | 0 | 3.44459617 | 0.82 | 0.138 | 0 |
| *F30A10.14* | 0 | 3.35881614 | 0.693 | 0.113 | 0 |
| *ttr-2* | 0 | 3.34558089 | 0.95 | 0.437 | 0 |
| *abf-2* | 0 | 3.28478792 | 0.466 | 0.098 | 0 |
| *nas-20* | 0 | 3.19213103 | 0.526 | 0.09 | 0 |
| *ule-2* | 0 | 3.10882655 | 0.859 | 0.263 | 0 |
| *K10C2.8* | 0 | 3.02523171 | 0.641 | 0.114 | 0 |
| *F17H10.2* | 0 | 2.57116443 | 0.678 | 0.127 | 0 |
| *ifa-1* | 0 | 2.56981153 | 0.698 | 0.167 | 0 |
| *K10C2.7* | 0 | 2.56414417 | 0.499 | 0.04 | 0 |
| *H01G02.1* | 0 | 2.4752212 | 0.457 | 0.086 | 0 |

### Top 20 depleted genes

| | p_val | avg_log2FC | pct.1 | pct.2 | p_val_adj |
|---|---|---|---|---|---|
| *crh-1* | 6.58E-27 | -1.4559612 | 0.134 | 0.269 | 3.08E-22 |
| *spsb-1* | 3.61E-17 | -1.37875 | 0.19 | 0.294 | 1.69E-12 |
| *Y105E8A.25* | 5.53E-25 | -1.3025727 | 0.221 | 0.352 | 2.59E-20 |
| *F53H2.3* | 1.34E-11 | -1.2233709 | 0.13 | 0.205 | 6.27E-07 |
| *F57F5.1* | 1.06E-23 | -1.2160887 | 0.396 | 0.505 | 4.96E-19 |
| *sos-1* | 3.47E-24 | -1.2081503 | 0.114 | 0.238 | 1.63E-19 |
| *mix-1* | 7.32E-24 | -1.1974038 | 0.169 | 0.3 | 3.43E-19 |
| *afd-1* | 1.65E-23 | -1.1973247 | 0.214 | 0.343 | 7.72E-19 |
| *daf-2* | 5.75E-22 | -1.1632197 | 0.093 | 0.206 | 2.70E-17 |
| *jun-1* | 2.66E-13 | -1.1246379 | 0.069 | 0.145 | 1.25E-08 |
| *ddo-2* | 3.22E-09 | -1.1239299 | 0.25 | 0.322 | 0.00015106 |
| *mnk-1* | 4.23E-18 | -1.0912991 | 0.107 | 0.211 | 1.98E-13 |
| *elpc-2* | 1.78E-20 | -1.0684162 | 0.093 | 0.202 | 8.37E-16 |
| *aakg-1* | 1.78E-18 | -1.0602111 | 0.11 | 0.213 | 8.37E-14 |
| *plg-1* | 1.14E-13 | -1.050763 | 0.143 | 0.23 | 5.36E-09 |
| *tos-1* | 5.21E-15 | -1.019243 | 0.423 | 0.502 | 2.44E-10 |
| *vit-2* | 4.48E-18 | -1.0115699 | 0.706 | 0.725 | 2.10E-13 |
| *lys-2* | 5.48E-17 | -1.0064367 | 0.39 | 0.478 | 2.57E-12 |
| *C23H5.8* | 6.11E-15 | -1.0052236 | 0.372 | 0.457 | 2.87E-10 |
| *asp-3* | 4.05E-22 | -1.0000237 | 0.511 | 0.596 | 1.90E-17 |

### Tissue enrichment

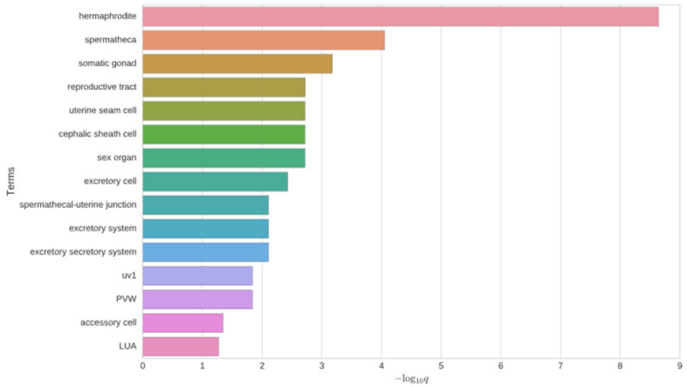


### GO enrichment

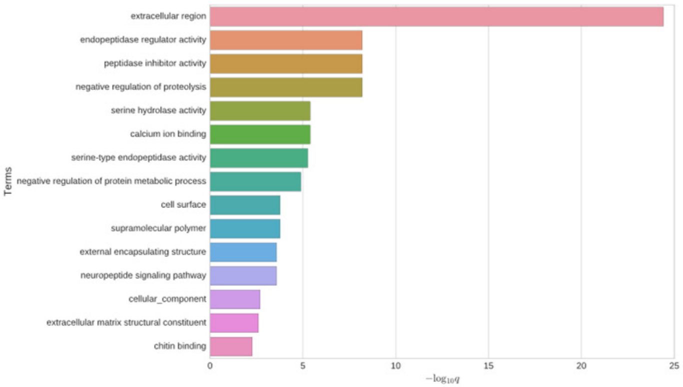


### Phenotype enrichment

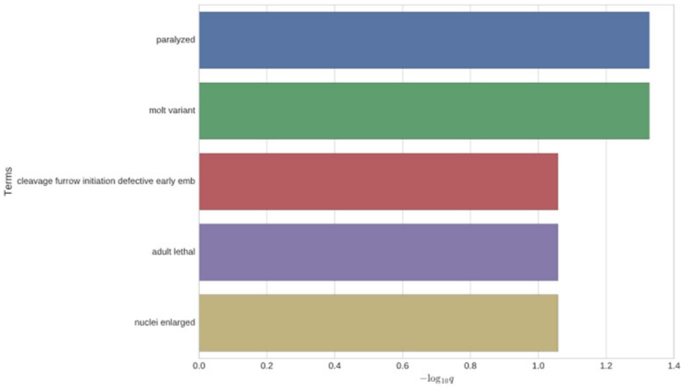


## *C35B1.4*

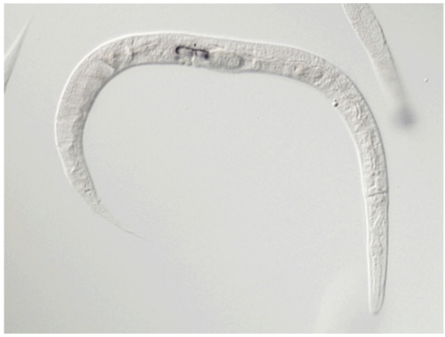

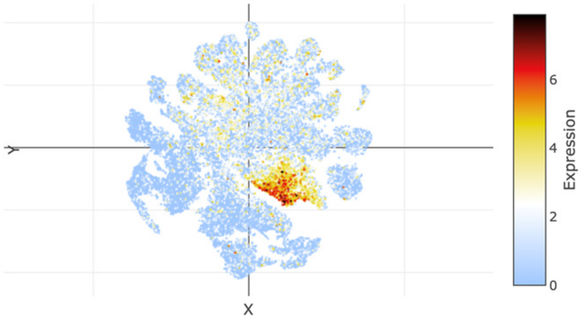


### *ifa-1*

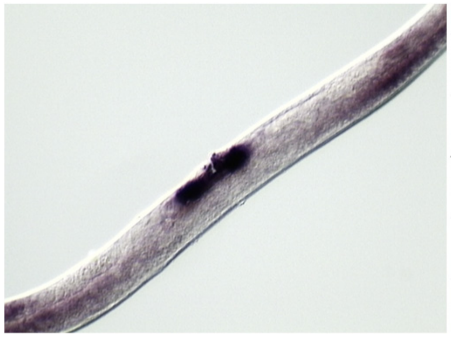

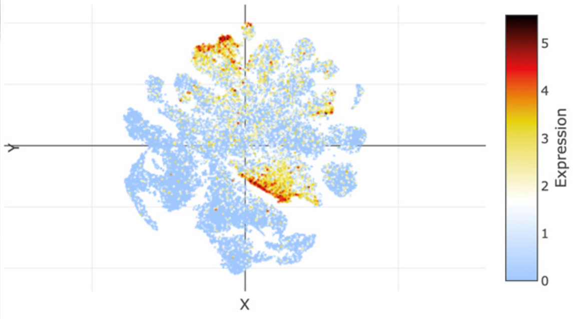

### t-SNE showing cell type 5

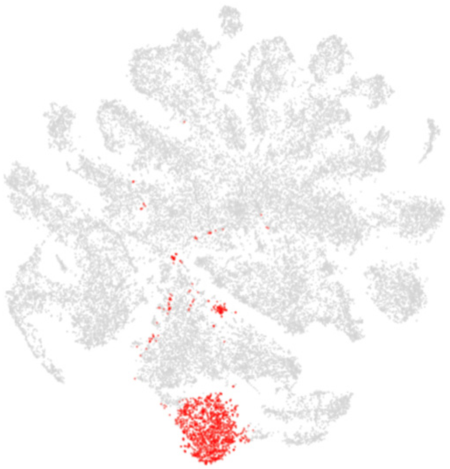

### Top 20 enriched genes

| | p_val | avg_log2FC | pct.1 | pct.2 | p_val_adj |
|---|---|---|---|---|---|
| *pho-11* | 0 | 3.29863936 | 0.925 | 0.345 | 0 |
| *Y39B6A.1* | 7.09E-271 | 3.12430316 | 0.857 | 0.412 | 3.33E-266 |
| *irg-7* | 0 | 3.09445961 | 0.928 | 0.332 | 0 |
| *B0035.13* | 0 | 2.73722258 | 0.789 | 0.186 | 0 |
| *T15B7.1* | 0 | 2.64635729 | 0.785 | 0.19 | 0 |
| *clec-48* | 0 | 2.58047161 | 0.821 | 0.249 | 0 |
| *C17F4.7* | 1.26E-259 | 2.31138011 | 0.96 | 0.702 | 5.93E-255 |
| *T25C12.3* | 2.37E-277 | 2.14591607 | 0.809 | 0.352 | 1.11E-272 |
| *asp-14* | 6.23E-266 | 2.07244978 | 0.687 | 0.22 | 2.92E-261 |
| *ZK6.11* | 7.79E-196 | 1.99431006 | 0.771 | 0.392 | 3.66E-191 |
| *elo-6* | 1.19E-240 | 1.89115393 | 0.911 | 0.536 | 5.59E-236 |
| *asp-5* | 4.11E-239 | 1.88841255 | 0.897 | 0.489 | 1.93E-234 |
| *F53A9.8* | 3.68E-250 | 1.88357533 | 0.606 | 0.172 | 1.72E-245 |
| *F19C7.1* | 9.48E-245 | 1.86857115 | 0.829 | 0.417 | 4.45E-240 |
| *mtl-1* | 7.60E-74 | 1.82874675 | 0.545 | 0.296 | 3.56E-69 |
| *clec-50* | 6.06E-252 | 1.82190629 | 0.936 | 0.553 | 2.84E-247 |
| *asp-2* | 1.24E-256 | 1.81738524 | 0.932 | 0.554 | 5.84E-252 |
| *Y34B4A.6* | 8.63E-257 | 1.80575958 | 0.855 | 0.416 | 4.05E-252 |
| *clec-47* | 6.57E-210 | 1.79353628 | 0.535 | 0.151 | 3.08E-205 |
| *drd-5* | 0 | 1.78454448 | 0.496 | 0.082 | 0 |

### Top 20 depleted genes

| | p_val | avg_log2FC | pct.1 | pct.2 | p_val_adj |
|---|---|---|---|---|---|
| *dig-1* | 2.51E-30 | -3.4157436 | 0.072 | 0.244 | 1.18E-25 |
| *ule-4* | 2.36E-24 | -2.5508011 | 0.37 | 0.512 | 1.10E-19 |
| *ZK813.7* | 2.89E-16 | -2.4964465 | 0.112 | 0.227 | 1.35E-11 |
| *ule-3* | 5.23E-15 | -2.4638828 | 0.131 | 0.246 | 2.45E-10 |
| *D1086.10* | 7.40E-23 | -2.4446669 | 0.247 | 0.398 | 3.47E-18 |
| *ZC513.7* | 1.03E-38 | -2.1423755 | 0.218 | 0.438 | 4.81E-34 |
| *C10G8.4* | 2.04E-14 | -2.1110012 | 0.231 | 0.347 | 9.59E-10 |
| *D1054.10* | 3.61E-10 | -1.9864446 | 0.23 | 0.324 | 1.70E-05 |
| *ZK813.1* | 4.72E-12 | -1.9754372 | 0.112 | 0.208 | 2.21E-07 |
| *ZK813.3* | 4.37E-10 | -1.9747983 | 0.156 | 0.244 | 2.05E-05 |
| *ule-2* | 7.62E-18 | -1.9733314 | 0.164 | 0.299 | 3.57E-13 |
| *ule-5* | 2.19E-11 | -1.9732913 | 0.233 | 0.331 | 1.03E-06 |
| *fasn-1* | 3.24E-14 | -1.9579536 | 0.165 | 0.28 | 1.52E-09 |
| *fipr-2* | 7.02E-17 | -1.952572 | 0.086 | 0.199 | 3.29E-12 |
| *Y62H9A.5* | 6.72E-13 | -1.9454639 | 0.179 | 0.285 | 3.15E-08 |
| *ZK813.2* | 1.02E-17 | -1.9269024 | 0.166 | 0.294 | 4.77E-13 |
| *ttn-1* | 5.63E-31 | -1.8747178 | 0.293 | 0.476 | 2.64E-26 |
| *Y37D8A.19* | 1.29E-09 | -1.8663619 | 0.273 | 0.36 | 6.05E-05 |
| *ule-1* | 6.23E-09 | -1.827216 | 0.184 | 0.267 | 0.00029223 |
| *ZC373.2* | 6.75E-09 | -1.7912988 | 0.253 | 0.336 | 0.00031654 |

### Tissue enrichment

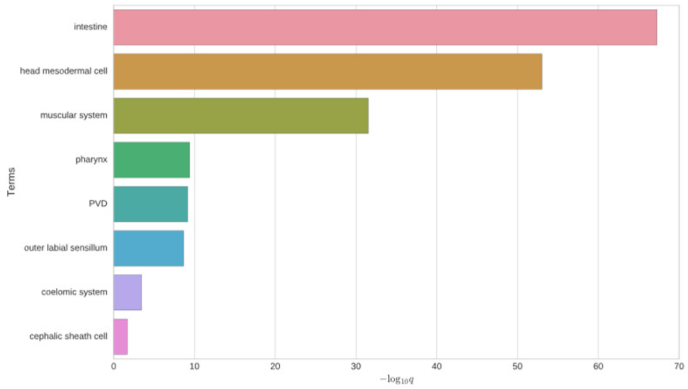


### GO enrichment

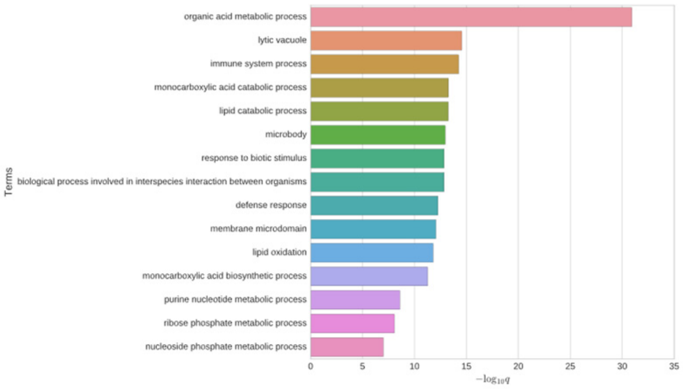


### Phenotype enrichment

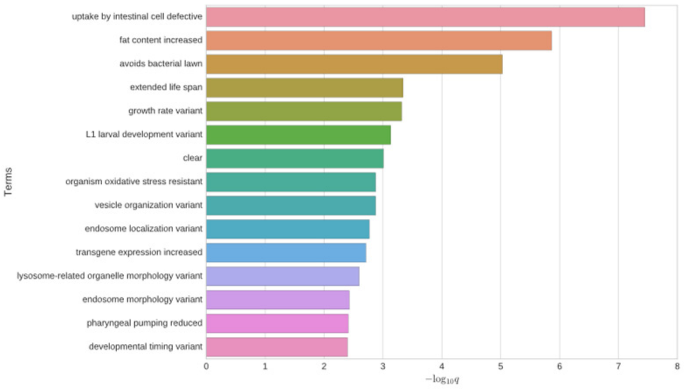


## *F53A9.8*

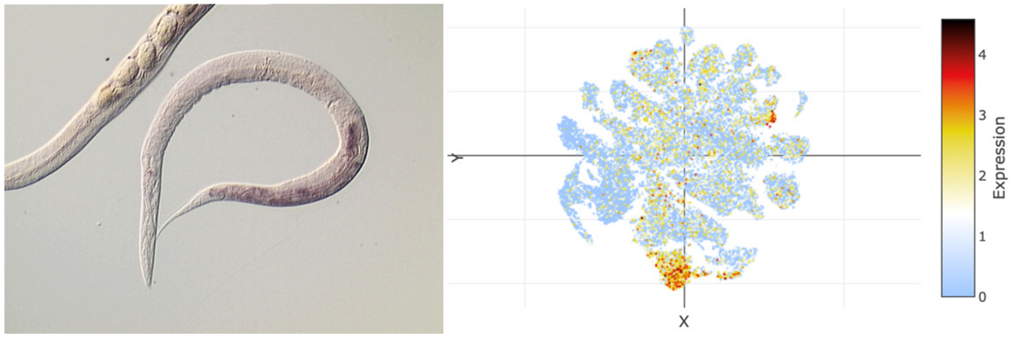


### *pho-11*

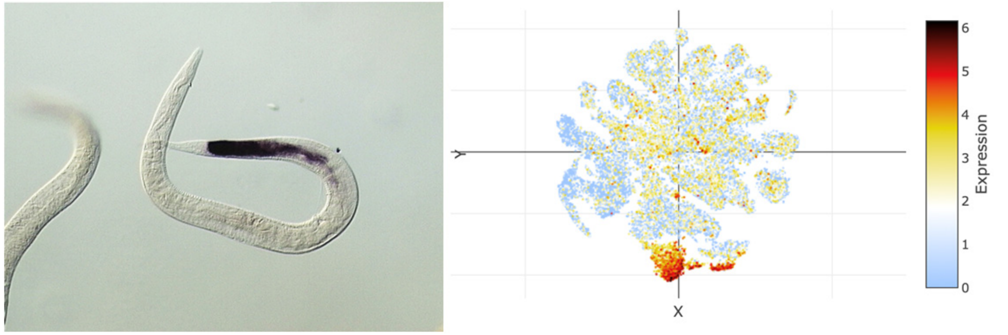

### t-SNE showing cell type 6

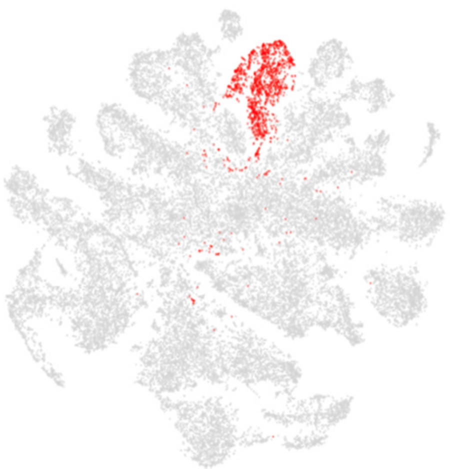

### Top 20 enriched genes

| | p_val | avg_log2FC | pct.1 | pct.2 | p_val_adj |
|---|---|---|---|---|---|
| *F41F3.3* | 0 | 4.44951499 | 0.749 | 0.067 | 0 |
| *col-77* | 0 | 4.33925381 | 0.769 | 0.087 | 0 |
| *dpy-13* | 0 | 4.14527702 | 0.713 | 0.076 | 0 |
| *dpy-4* | 0 | 4.11006311 | 0.739 | 0.079 | 0 |
| *col-71* | 0 | 4.07774858 | 0.702 | 0.079 | 0 |
| *col-125* | 0 | 3.99889991 | 0.815 | 0.12 | 0 |
| *C02E7.6* | 0 | 3.99220953 | 0.739 | 0.094 | 0 |
| *col-38* | 0 | 3.97704091 | 0.683 | 0.074 | 0 |
| *F53F1.4* | 0 | 3.91022414 | 0.803 | 0.18 | 0 |
| *Y48G8AL.12* | 0 | 3.90138684 | 0.676 | 0.062 | 0 |
| *col-73* | 0 | 3.84007543 | 0.696 | 0.07 | 0 |
| *bli-6* | 0 | 3.8038908 | 0.807 | 0.129 | 0 |
| *col-161* | 0 | 3.79978985 | 0.741 | 0.098 | 0 |
| *col-175* | 0 | 3.79038887 | 0.633 | 0.052 | 0 |
| *col-107* | 0 | 3.75951143 | 0.732 | 0.079 | 0 |
| *col-138* | 0 | 3.73929899 | 0.627 | 0.059 | 0 |
| *col-162* | 0 | 3.68473222 | 0.672 | 0.086 | 0 |
| *C02E7.7* | 0 | 3.68014325 | 0.643 | 0.076 | 0 |
| *ram-2* | 0 | 3.60736456 | 0.865 | 0.226 | 0 |
| *col-14* | 0 | 3.56049401 | 0.708 | 0.051 | 0 |

### Top 20 depleted genes

| | p_val | avg_log2FC | pct.1 | pct.2 | p_val_adj |
|---|---|---|---|---|---|
| *dig-1* | 2.17E-08 | -2.1578555 | 0.318 | 0.233 | 0.00101789 |
| *ule-1* | 1.52E-26 | -2.0047118 | 0.126 | 0.271 | 7.13E-22 |
| *C10G8.4* | 1.93E-11 | -1.9180115 | 0.264 | 0.347 | 9.06E-07 |
| *C39D10.7* | 7.11E-15 | -1.907543 | 0.185 | 0.288 | 3.33E-10 |
| *vit-5* | 5.23E-98 | -1.85104 | 0.577 | 0.774 | 2.45E-93 |
| *Y62H9A.5* | 9.84E-20 | -1.6864584 | 0.163 | 0.287 | 4.61E-15 |
| *ule-3* | 3.46E-09 | -1.6826192 | 0.17 | 0.246 | 0.00016222 |
| *vit-2* | 7.90E-78 | -1.6602462 | 0.55 | 0.733 | 3.71E-73 |
| *T21C9.13* | 5.32E-18 | -1.6488546 | 0.062 | 0.158 | 2.49E-13 |
| *col-95* | 8.51E-53 | -1.6478596 | 0.405 | 0.598 | 3.99E-48 |
| *clec-87* | 3.81E-39 | -1.6052114 | 0.183 | 0.377 | 1.79E-34 |
| *Y22D7AR.10* | 1.60E-11 | -1.5674783 | 0.214 | 0.298 | 7.49E-07 |
| *vit-6* | 3.75E-76 | -1.5588859 | 0.597 | 0.763 | 1.76E-71 |
| *F56D6.8* | 2.93E-29 | -1.5506839 | 0.079 | 0.219 | 1.37E-24 |
| *ule-2* | 1.61E-08 | -1.5339991 | 0.225 | 0.297 | 0.000756 |
| *ule-5* | 9.54E-11 | -1.4958015 | 0.247 | 0.331 | 4.48E-06 |
| *T24B8.5* | 2.37E-15 | -1.4932008 | 0.328 | 0.412 | 1.11E-10 |
| *ZK813.1* | 1.28E-16 | -1.4378633 | 0.105 | 0.209 | 5.98E-12 |
| *F56D6.9* | 1.67E-27 | -1.4338885 | 0.123 | 0.265 | 7.81E-23 |
| *vit-3* | 3.93E-58 | -1.4144058 | 0.567 | 0.72 | 1.84E-53 |

### Tissue enrichment

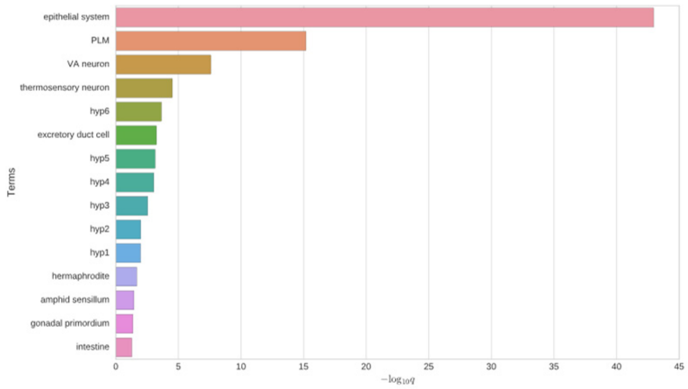


### GO enrichment

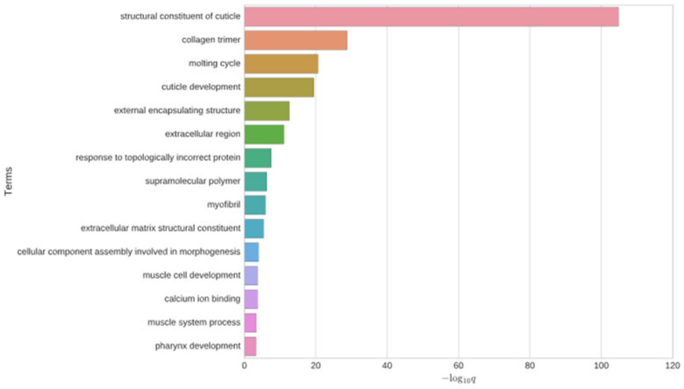


### Phenotype enrichment

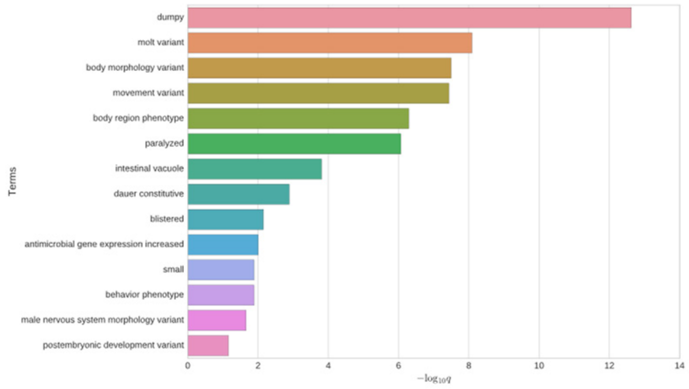


### *dpy-5*

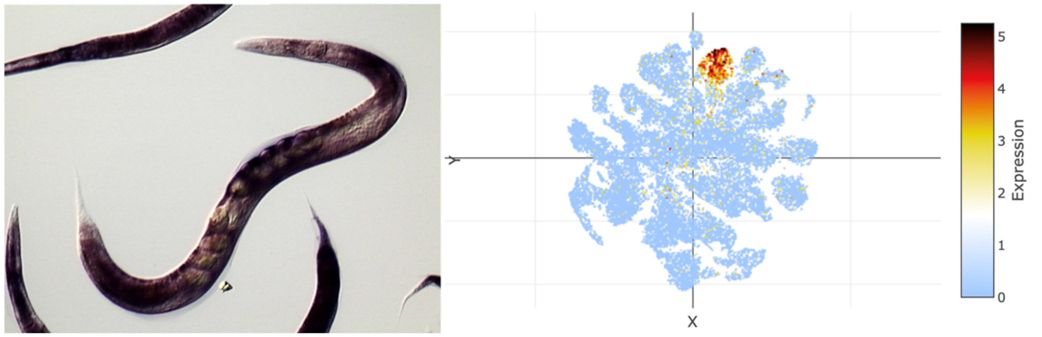


### *sqt-3*

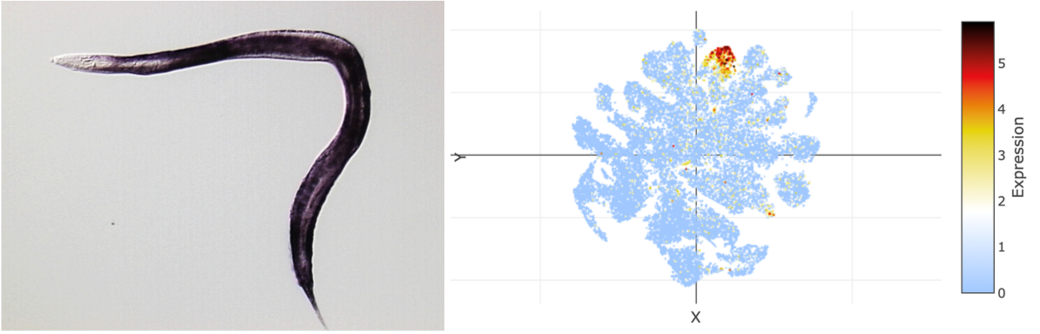

### t-SNE showing cell type 7

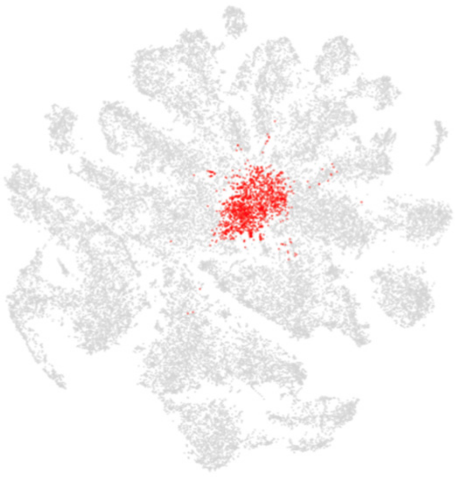

### Top 20 enriched genes

| | p_val | avg_log2FC | pct.1 | pct.2 | p_val_adj |
|---|---|---|---|---|---|
| *grd-3* | 1.72E-158 | 2.37200642 | 0.776 | 0.419 | 8.09E-154 |
| *grd-5* | 8.85E-155 | 2.2833273 | 0.796 | 0.45 | 4.15E-150 |
| *col-95* | 1.37E-231 | 2.22281109 | 0.942 | 0.574 | 6.43E-227 |
| *col-103* | 4.46E-284 | 2.05943165 | 0.98 | 0.624 | 2.09E-279 |
| *col-42* | 1.52E-284 | 2.05906452 | 0.965 | 0.594 | 7.11E-280 |
| *col-98* | 7.94E-296 | 2.03754652 | 0.984 | 0.622 | 3.72E-291 |
| *D1086.3* | 4.47E-154 | 2.01044005 | 0.693 | 0.34 | 2.10E-149 |
| *col-119* | 0 | 1.9486887 | 0.998 | 0.719 | 0 |
| *col-101* | 1.87E-299 | 1.9321734 | 0.978 | 0.705 | 8.76E-295 |
| *F15E6.3* | 2.58E-103 | 1.92641576 | 0.54 | 0.246 | 1.21E-98 |
| *brp-1* | 5.97E-229 | 1.91297689 | 0.93 | 0.537 | 2.80E-224 |
| *H39E23.3* | 1.50E-58 | 1.89606716 | 0.2 | 0.061 | 7.03E-54 |
| *osm-11* | 5.70E-169 | 1.88990853 | 0.722 | 0.339 | 2.67E-164 |
| *col-143* | 6.96E-293 | 1.85958865 | 0.991 | 0.737 | 3.26E-288 |
| *col-8* | 5.27E-268 | 1.83289202 | 0.968 | 0.632 | 2.47E-263 |
| *F56D6.9* | 1.90E-83 | 1.83251155 | 0.515 | 0.247 | 8.93E-79 |
| *dct-8* | 4.47E-108 | 1.82923756 | 0.44 | 0.16 | 2.10E-103 |
| *F56D6.8* | 1.08E-79 | 1.79915976 | 0.452 | 0.202 | 5.07E-75 |
| *col-106* | 8.37E-293 | 1.79407149 | 0.989 | 0.672 | 3.93E-288 |
| *col-140* | 0 | 1.78407051 | 0.998 | 0.727 | 0 |

### Top 20 depleted genes

| | p_val | avg_log2FC | pct.1 | pct.2 | p_val_adj |
|---|---|---|---|---|---|
| *D1086.10* | 3.71E-11 | -2.5933411 | 0.313 | 0.396 | 1.74E-06 |
| *ule-4* | 2.11E-07 | -2.4563715 | 0.488 | 0.508 | 0.00990206 |
| *D1054.10* | 1.82E-10 | -2.4090505 | 0.237 | 0.324 | 8.52E-06 |
| *ule-5* | 2.58E-08 | -1.9182479 | 0.252 | 0.33 | 0.00120943 |
| *ZK813.2* | 8.94E-17 | -1.9072577 | 0.173 | 0.294 | 4.19E-12 |
| *Y37D8A.19* | 8.29E-07 | -1.8344605 | 0.29 | 0.359 | 0.03886863 |
| *fipr-2* | 8.39E-11 | -1.673803 | 0.112 | 0.198 | 3.93E-06 |
| *T21C9.13* | 6.91E-11 | -1.6038852 | 0.075 | 0.156 | 3.24E-06 |
| *F55B11.2* | 2.32E-08 | -1.5857711 | 0.135 | 0.209 | 0.00108911 |
| *W02D9.7* | 9.26E-10 | -1.5803547 | 0.158 | 0.244 | 4.34E-05 |
| *C53B7.3* | 2.33E-11 | -1.503996 | 0.162 | 0.256 | 1.09E-06 |
| *F57C2.4* | 5.19E-07 | -1.4784079 | 0.178 | 0.245 | 0.02435468 |
| *C49G7.3* | 1.50E-09 | -1.4349223 | 0.108 | 0.188 | 7.04E-05 |
| *F55B11.3* | 3.79E-07 | -1.3736901 | 0.1 | 0.164 | 0.01776466 |
| *Y45F10C.4* | 2.01E-13 | -1.3289441 | 0.126 | 0.231 | 9.41E-09 |
| *Y105E8A.25* | 8.38E-24 | -1.3263241 | 0.193 | 0.351 | 3.93E-19 |
| *D1086.7* | 1.42E-10 | -1.3262246 | 0.146 | 0.237 | 6.66E-06 |
| *T20G5.8* | 8.25E-08 | -1.319649 | 0.106 | 0.174 | 0.0038719 |
| *Y62H9A.4* | 9.84E-09 | -1.3189104 | 0.124 | 0.2 | 0.00046158 |
| *Y39B6A.1* | 1.73E-14 | -1.2559505 | 0.32 | 0.434 | 8.10E-10 |

### Tissue enrichment

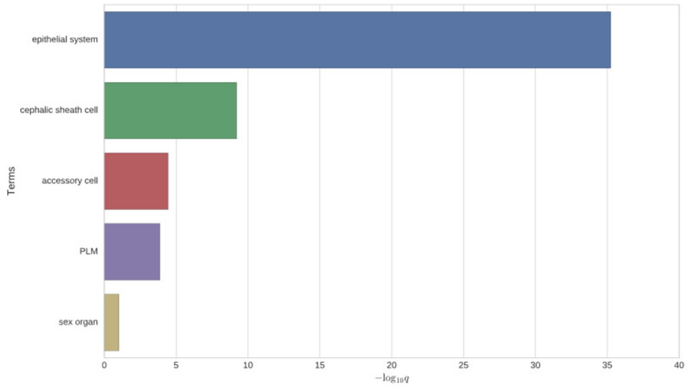


### GO enrichment

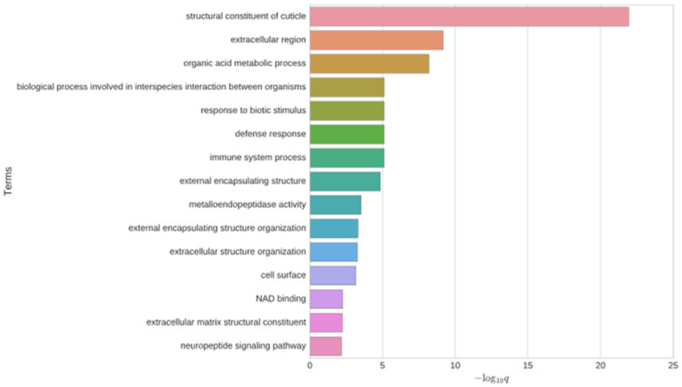


### Phenotype enrichment

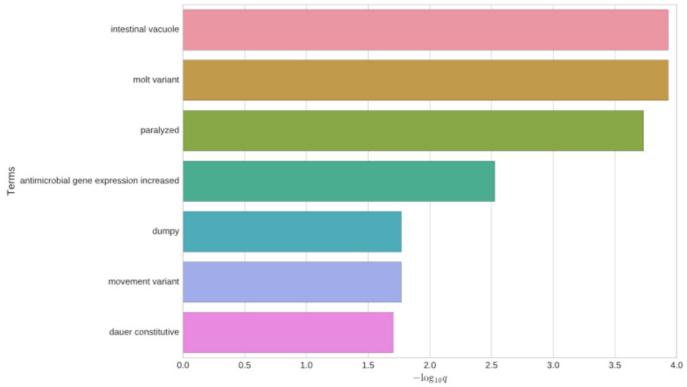


### *dct-8*

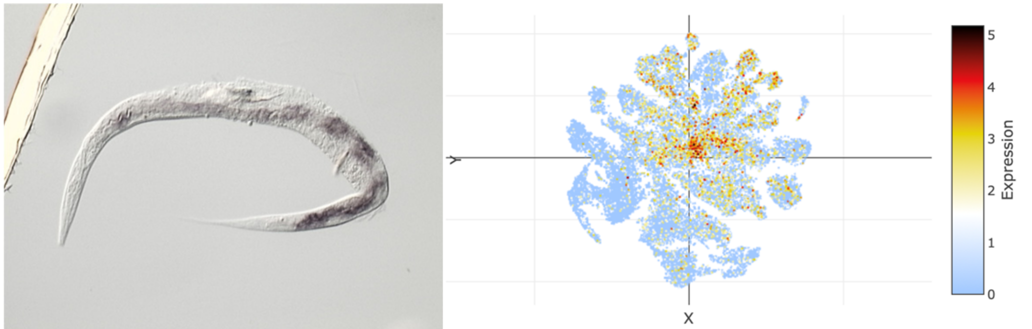


### *osm-11*

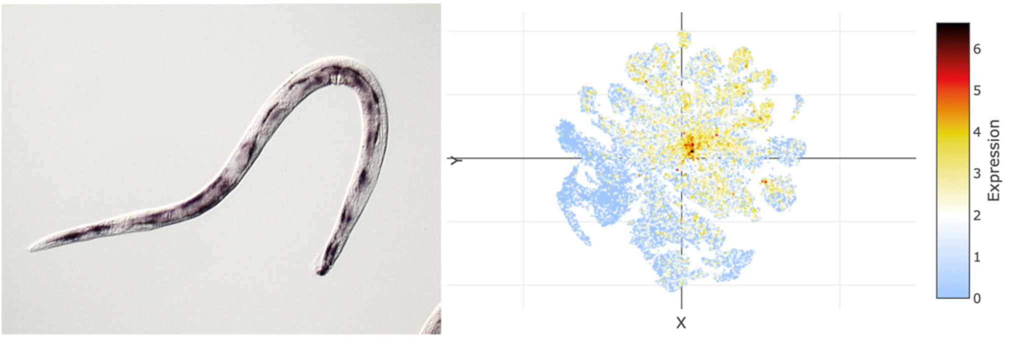

### t-SNE showing cell type 8

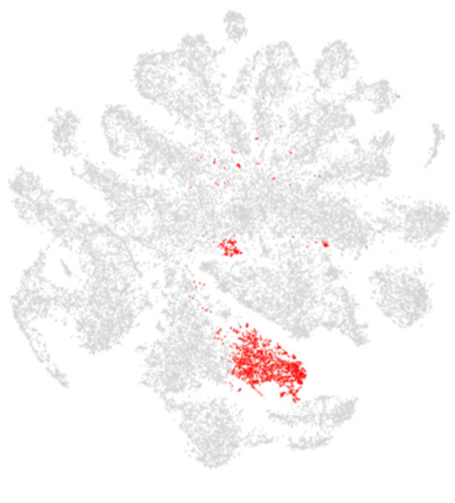

### Top 20 enriched genes

| | p_val | avg_log2FC | pct.1 | pct.2 | p_val_adj |
|---|---|---|---|---|---|
| *lys-7* | 0 | 4.0594949 | 0.963 | 0.298 | 0 |
| *F55G11.4* | 0 | 3.70860887 | 0.759 | 0.191 | 0 |
| *C14C6.5* | 0 | 3.40609692 | 0.871 | 0.219 | 0 |
| *F01D5.5* | 0 | 3.21148937 | 0.704 | 0.104 | 0 |
| *dod-19* | 0 | 3.10116108 | 0.905 | 0.344 | 0 |
| *F13D12.6* | 0 | 3.08995782 | 0.953 | 0.365 | 0 |
| *spp-2* | 0 | 2.86771753 | 0.858 | 0.283 | 0 |
| *tag-10* | 0 | 2.82908937 | 0.718 | 0.12 | 0 |
| *T24B8.5* | 0 | 2.71803449 | 0.895 | 0.387 | 0 |
| *C14C6.2* | 0 | 2.67714794 | 0.668 | 0.082 | 0 |
| *clec-41* | 0 | 2.63464402 | 0.754 | 0.178 | 0 |
| *T01D3.6* | 0 | 2.56268599 | 0.778 | 0.197 | 0 |
| *C50F7.5* | 0 | 2.54477688 | 0.504 | 0.08 | 0 |
| *cpr-1* | 0 | 2.52352544 | 0.938 | 0.431 | 0 |
| *F52E1.14* | 0 | 2.46028867 | 0.847 | 0.337 | 0 |
| *Y43C5A.2* | 0 | 2.42181633 | 0.729 | 0.208 | 0 |
| *lys-1* | 0 | 2.40233128 | 0.961 | 0.553 | 0 |
| *lys-8* | 0 | 2.32468153 | 0.928 | 0.455 | 0 |
| *spp-14* | 0 | 2.31550767 | 0.956 | 0.587 | 0 |
| *F01D5.1* | 0 | 2.31320803 | 0.652 | 0.134 | 0 |

### Top 20 depleted genes

| | p_val | avg_log2FC | pct.1 | pct.2 | p_val_adj |
|---|---|---|---|---|---|
| *dig-1* | 2.02E-15 | -2.8017102 | 0.128 | 0.242 | 9.48E-11 |
| *ule-3* | 9.65E-11 | -1.9787025 | 0.158 | 0.246 | 4.52E-06 |
| *ule-4* | 8.31E-15 | -1.9074952 | 0.415 | 0.511 | 3.90E-10 |
| *ZC513.7* | 6.49E-26 | -1.8628847 | 0.271 | 0.436 | 3.04E-21 |
| *fasn-1* | 1.88E-13 | -1.8489222 | 0.176 | 0.28 | 8.81E-09 |
| *C10G8.4* | 1.16E-11 | -1.805678 | 0.252 | 0.347 | 5.44E-07 |
| *D1054.10* | 4.28E-07 | -1.7656258 | 0.253 | 0.324 | 0.02006366 |
| *ule-1* | 8.13E-11 | -1.7613421 | 0.178 | 0.268 | 3.81E-06 |
| *C39D10.7* | 3.83E-08 | -1.7500612 | 0.211 | 0.286 | 0.00179501 |
| *ule-2* | 1.37E-14 | -1.7229286 | 0.184 | 0.298 | 6.43E-10 |
| *Y62H9A.5* | 3.28E-09 | -1.677695 | 0.204 | 0.284 | 0.00015397 |
| *ZK813.1* | 5.11E-08 | -1.6555386 | 0.136 | 0.207 | 0.00239575 |
| *D1086.10* | 7.27E-14 | -1.5848056 | 0.284 | 0.397 | 3.41E-09 |
| *inos-1* | 2.85E-14 | -1.5581579 | 0.045 | 0.132 | 1.34E-09 |
| *Y22D7AR.10* | 2.06E-10 | -1.5114018 | 0.21 | 0.297 | 9.65E-06 |
| *T21C9.13* | 2.02E-09 | -1.5102531 | 0.085 | 0.156 | 9.46E-05 |
| *Y37D8A.19* | 1.09E-07 | -1.5046894 | 0.288 | 0.36 | 0.00512857 |
| *ule-5* | 6.96E-08 | -1.4536947 | 0.258 | 0.33 | 0.00326462 |
| *ttn-1* | 1.11E-14 | -1.4025093 | 0.367 | 0.473 | 5.20E-10 |
| *crh-1* | 1.68E-20 | -1.3867873 | 0.14 | 0.267 | 7.89E-16 |

### Tissue enrichment

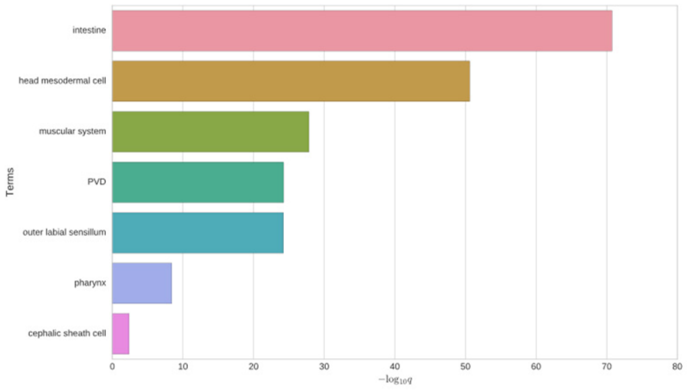


### GO enrichment

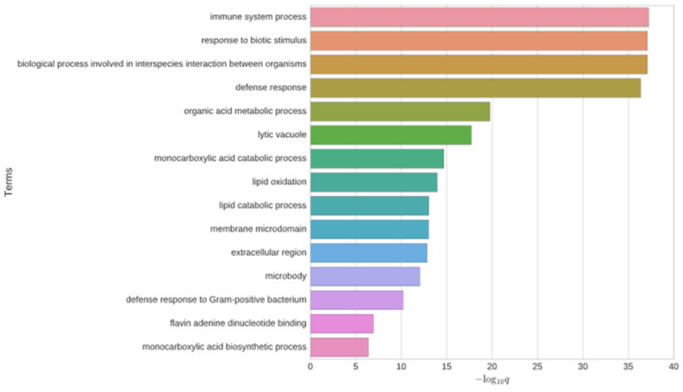


### Phenotype enrichment

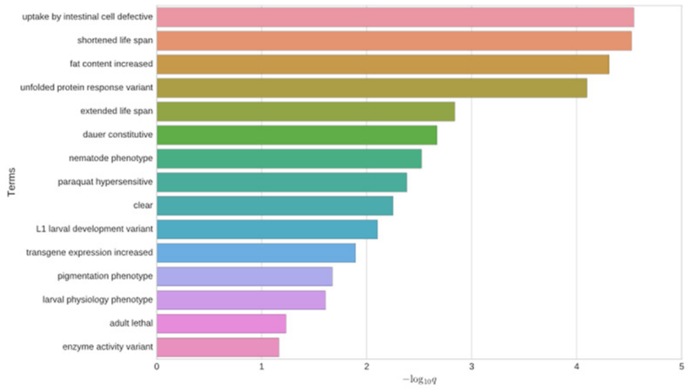


### *lys-7*

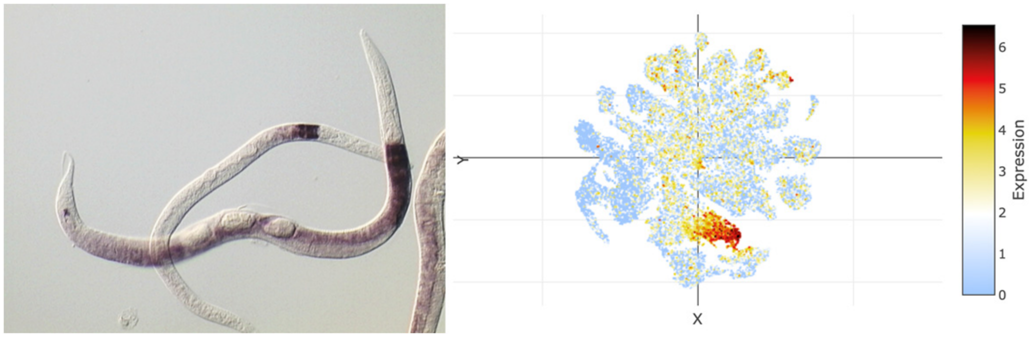


### *dod-19*

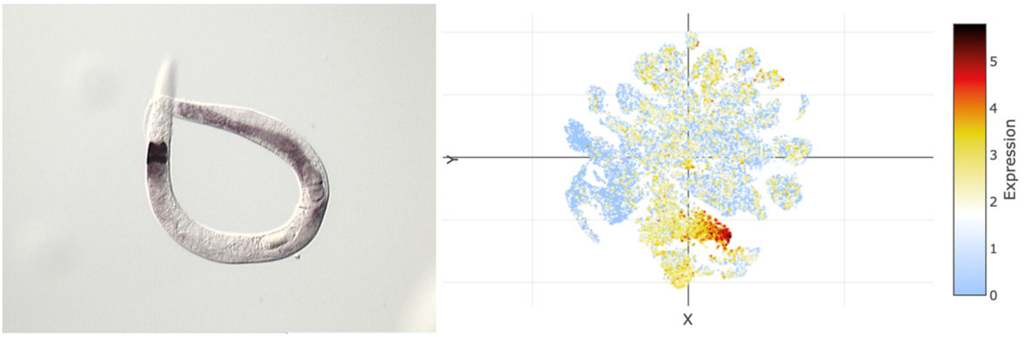

### t-SNE showing cell type 9

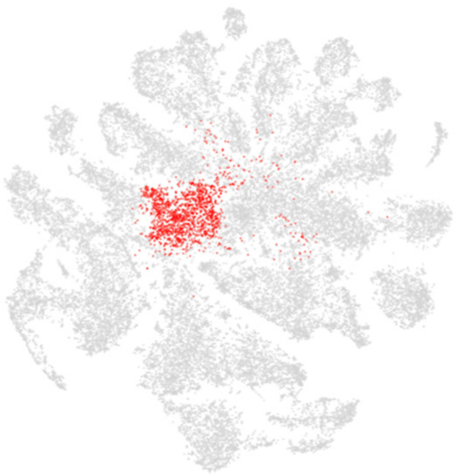

### Top 20 enriched genes

| | p_val | avg_log2FC | pct.1 | pct.2 | p_val_adj |
|---|---|---|---|---|---|
| *cey-2* | 8.14E-131 | 1.14445315 | 0.669 | 0.311 | 3.82E-126 |
| *cpg-1* | 4.24E-141 | 1.11774305 | 0.681 | 0.309 | 1.99E-136 |
| *cpg-2* | 1.23E-135 | 1.07027514 | 0.729 | 0.363 | 5.76E-131 |
| *cbd-1* | 2.32E-143 | 1.06625001 | 0.699 | 0.316 | 1.09E-138 |
| *sip-1* | 3.27E-131 | 1.02858936 | 0.784 | 0.428 | 1.53E-126 |
| *rmd-1* | 9.58E-96 | 1.00273309 | 0.579 | 0.288 | 4.49E-91 |
| *clec-87* | 6.67E-148 | 0.99947049 | 0.731 | 0.35 | 3.13E-143 |
| *mesp-1* | 1.08E-88 | 0.94704534 | 0.428 | 0.176 | 5.08E-84 |
| *gln-6* | 5.85E-102 | 0.94194278 | 0.515 | 0.219 | 2.75E-97 |
| *W05F2.3* | 3.47E-103 | 0.9211276 | 0.581 | 0.271 | 1.63E-98 |
| *cpg-3* | 1.70E-92 | 0.86340233 | 0.661 | 0.352 | 8.00E-88 |
| *ran-1* | 1.47E-75 | 0.85027519 | 0.592 | 0.324 | 6.88E-71 |
| *nasp-2* | 2.18E-87 | 0.84774327 | 0.567 | 0.274 | 1.02E-82 |
| *rnr-2* | 3.86E-98 | 0.845464 | 0.47 | 0.191 | 1.81E-93 |
| *cgh-1* | 4.50E-84 | 0.84188692 | 0.693 | 0.411 | 2.11E-79 |
| *clec-91* | 2.62E-73 | 0.81629824 | 0.466 | 0.22 | 1.23E-68 |
| *rpl-11.1* | 3.46E-78 | 0.80981696 | 0.587 | 0.303 | 1.62E-73 |
| *M28.5* | 4.17E-68 | 0.79616773 | 0.415 | 0.186 | 1.96E-63 |
| *clec-88* | 6.91E-83 | 0.77826019 | 0.427 | 0.178 | 3.24E-78 |
| *hil-4* | 3.42E-83 | 0.77802479 | 0.598 | 0.298 | 1.60E-78 |

### Top 20 depleted genes

| | p_val | avg_log2FC | pct.1 | pct.2 | p_val_adj |
|---|---|---|---|---|---|
| *C30E1.9* | 8.11E-14 | -1.4485576 | 0.401 | 0.487 | 3.80E-09 |
| *F53H2.3* | 7.03E-18 | -1.4376546 | 0.098 | 0.206 | 3.30E-13 |
| *Y105E8A.25* | 4.84E-25 | -1.4312299 | 0.213 | 0.351 | 2.27E-20 |
| *crh-1* | 1.75E-20 | -1.4079217 | 0.145 | 0.267 | 8.22E-16 |
| *D1054.10* | 2.71E-14 | -1.3622332 | 0.464 | 0.314 | 1.27E-09 |
| *D1086.10* | 1.27E-13 | -1.3463393 | 0.544 | 0.385 | 5.94E-09 |
| *spsb-1* | 1.18E-12 | -1.2972569 | 0.2 | 0.292 | 5.54E-08 |
| *ZK813.7* | 1.25E-20 | -1.2869616 | 0.366 | 0.216 | 5.86E-16 |
| *sos-1* | 4.23E-24 | -1.2383515 | 0.102 | 0.238 | 1.98E-19 |
| *tos-1* | 3.12E-16 | -1.2051101 | 0.417 | 0.502 | 1.46E-11 |
| *anc-1* | 2.12E-30 | -1.2044106 | 0.6 | 0.694 | 9.95E-26 |
| *ZK813.1* | 8.79E-15 | -1.2003792 | 0.321 | 0.199 | 4.12E-10 |
| *aakg-1* | 2.84E-20 | -1.1985001 | 0.096 | 0.213 | 1.33E-15 |
| *mnk-1* | 1.87E-18 | -1.192513 | 0.099 | 0.21 | 8.79E-14 |
| *Y62H9A.5* | 3.14E-13 | -1.1890592 | 0.407 | 0.275 | 1.47E-08 |
| *ule-4* | 7.80E-30 | -1.1852717 | 0.728 | 0.497 | 3.66E-25 |
| *daf-2* | 1.75E-16 | -1.1571312 | 0.102 | 0.205 | 8.19E-12 |
| *ule-5* | 8.36E-17 | -1.1499111 | 0.482 | 0.32 | 3.92E-12 |
| *afd-1* | 4.09E-12 | -1.1379935 | 0.255 | 0.34 | 1.92E-07 |
| *ell-1* | 8.14E-19 | -1.1126796 | 0.288 | 0.403 | 3.82E-14 |

### Tissue enrichment

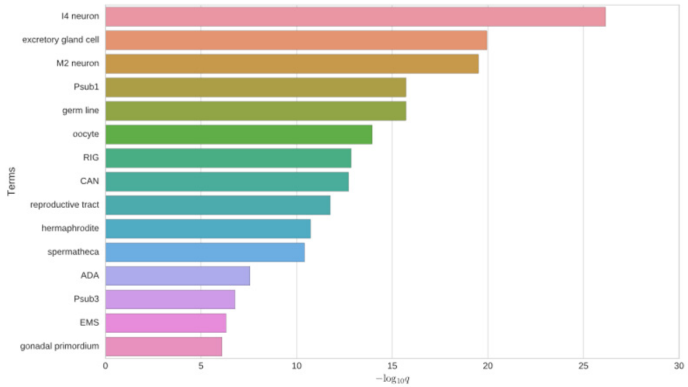


### GO enrichment

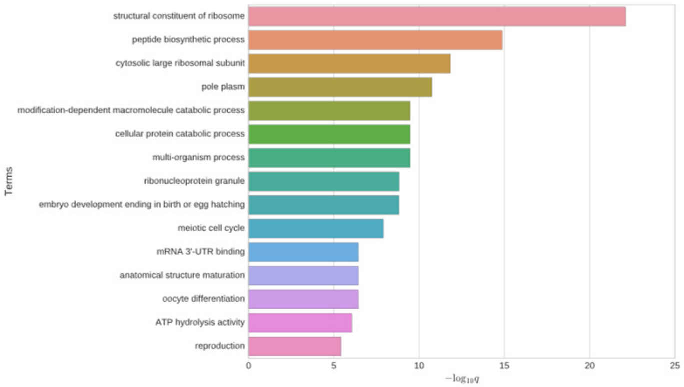


### Phenotype enrichment

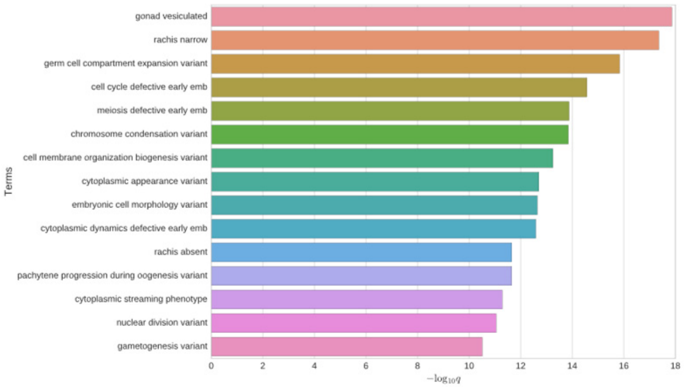


### *rnr-2*

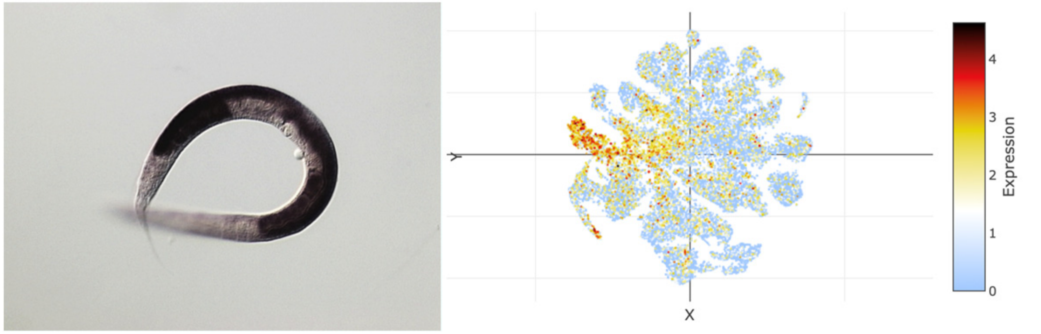


### *cey-2*

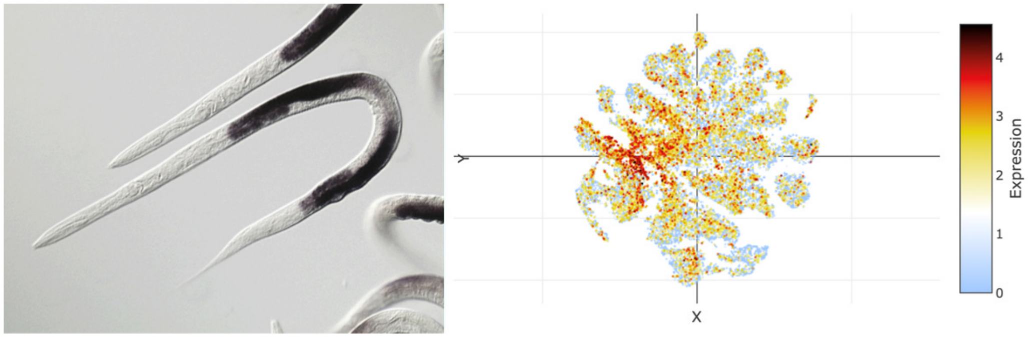

### t-SNE showing cell type 10

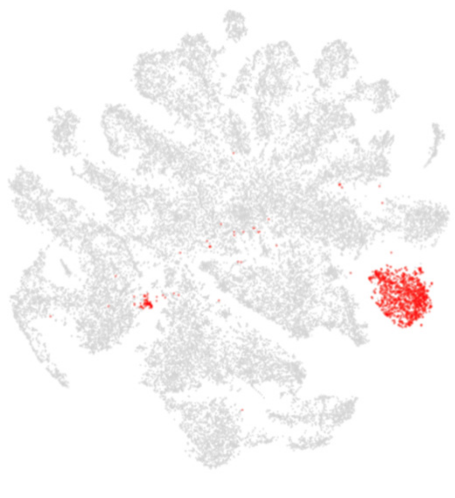

### Top 20 enriched genes

| | p_val | avg_log2FC | pct.1 | pct.2 | p_val_adj |
|---|---|---|---|---|---|
| *spsb-1* | 0 | 3.48086164 | 0.894 | 0.266 | 0 |
| *D2023.1* | 1.96E-277 | 3.1846028 | 0.428 | 0.075 | 9.21E-273 |
| *crh-1* | 0 | 3.12326781 | 0.848 | 0.24 | 0 |
| *Y37A1A.2* | 0 | 3.05464434 | 0.45 | 0.069 | 0 |
| *C15A11.7* | 1.84E-214 | 2.94361625 | 0.39 | 0.078 | 8.65E-210 |
| *F23F12.12* | 4.62E-183 | 2.93421689 | 0.755 | 0.417 | 2.17E-178 |
| *F16C3.2* | 7.50E-273 | 2.75324784 | 0.473 | 0.095 | 3.52E-268 |
| *afd-1* | 0 | 2.73694822 | 0.855 | 0.317 | 0 |
| *T12A2.1* | 0 | 2.73264997 | 0.531 | 0.091 | 0 |
| *fkh-7* | 1.45E-296 | 2.68763878 | 0.642 | 0.167 | 6.79E-292 |
| *ist-1* | 1.84E-272 | 2.65253224 | 0.453 | 0.084 | 8.63E-268 |
| *Y17G7B.10* | 1.41E-198 | 2.64500738 | 0.615 | 0.223 | 6.60E-194 |
| *mnk-1* | 0 | 2.61023332 | 0.724 | 0.186 | 0 |
| *ell-1* | 5.30E-253 | 2.60624036 | 0.82 | 0.383 | 2.48E-248 |
| *linc-72* | 0 | 2.5929721 | 0.364 | 0.031 | 0 |
| *sms-3* | 2.72E-269 | 2.54123223 | 0.564 | 0.138 | 1.28E-264 |
| *daf-2* | 0 | 2.5182282 | 0.703 | 0.182 | 0 |
| *C30E1.9* | 7.63E-287 | 2.49428335 | 0.927 | 0.467 | 3.58E-282 |
| *lmd-3* | 6.82E-239 | 2.46861555 | 0.808 | 0.392 | 3.20E-234 |
| *lim-9* | 5.50E-198 | 2.4685942 | 0.612 | 0.219 | 2.58E-193 |

### Top 20 depleted genes

| | p_val | avg_log2FC | pct.1 | pct.2 | p_val_adj |
|---|---|---|---|---|---|
| *ule-4* | 5.69E-10 | -2.3953821 | 0.439 | 0.509 | 2.67E-05 |
| *pat-10* | 6.54E-90 | -2.2979457 | 0.473 | 0.715 | 3.07E-85 |
| *C10G8.4* | 1.67E-14 | -2.2961586 | 0.226 | 0.347 | 7.84E-10 |
| *perm-4* | 2.63E-43 | -2.276476 | 0.375 | 0.586 | 1.24E-38 |
| *lev-11* | 3.32E-110 | -2.275305 | 0.438 | 0.737 | 1.56E-105 |
| *D1086.10* | 1.52E-08 | -2.2750585 | 0.325 | 0.395 | 0.00071474 |
| *far-2* | 8.08E-98 | -2.2374198 | 0.579 | 0.773 | 3.79E-93 |
| *act-4* | 2.33E-99 | -2.2255171 | 0.462 | 0.728 | 1.10E-94 |
| *lbp-2* | 2.64E-78 | -2.2120837 | 0.369 | 0.641 | 1.24E-73 |
| *ttr-16* | 1.39E-78 | -2.1985425 | 0.393 | 0.653 | 6.54E-74 |
| *ZK813.2* | 1.66E-24 | -2.1542344 | 0.13 | 0.295 | 7.76E-20 |
| *cpn-3* | 9.73E-77 | -2.069591 | 0.455 | 0.69 | 4.57E-72 |
| *mlc-2* | 1.10E-88 | -2.0602468 | 0.346 | 0.646 | 5.18E-84 |
| *mlc-3* | 1.73E-82 | -2.0109373 | 0.424 | 0.691 | 8.11E-78 |
| *ule-2* | 7.07E-10 | -2.0104303 | 0.206 | 0.297 | 3.32E-05 |
| *act-1* | 6.49E-93 | -2.0009527 | 0.553 | 0.781 | 3.04E-88 |
| *tnt-2* | 8.44E-72 | -1.9978179 | 0.339 | 0.611 | 3.96E-67 |
| *unc-87* | 2.64E-84 | -1.9953146 | 0.32 | 0.639 | 1.24E-79 |
| *unc-27* | 3.47E-68 | -1.96237 | 0.447 | 0.674 | 1.63E-63 |
| *act-3* | 3.50E-104 | -1.9411241 | 0.497 | 0.783 | 1.64E-99 |

### Tissue enrichment

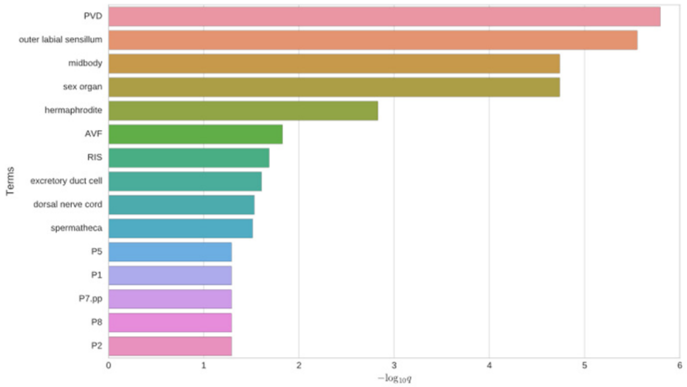


### GO enrichment

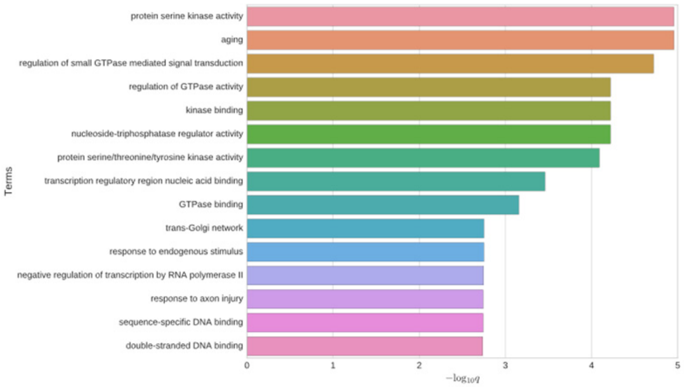


### Phenotype enrichment

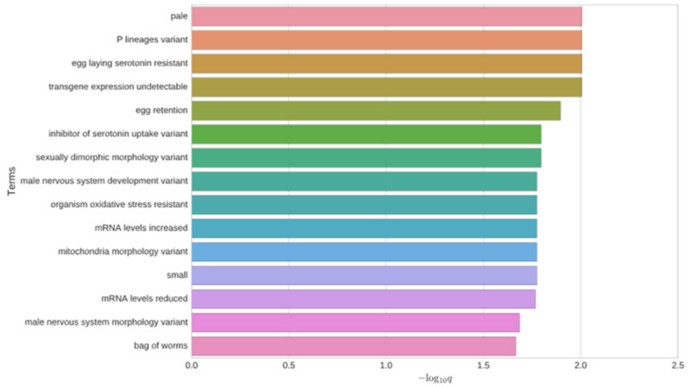


## *C15A11.7*

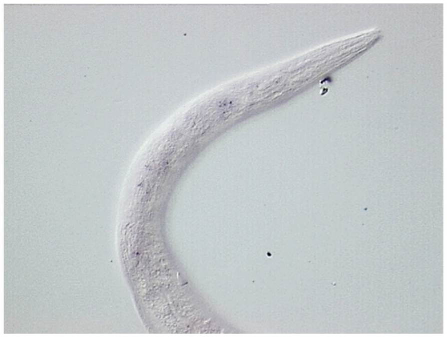

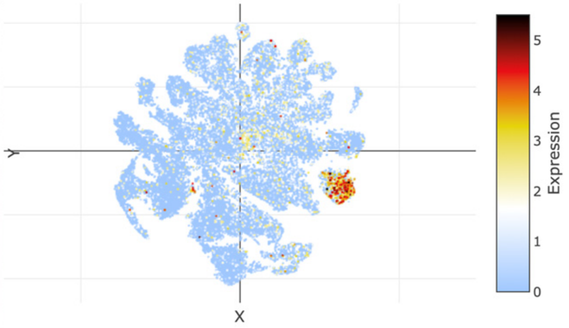


### *ist-1*

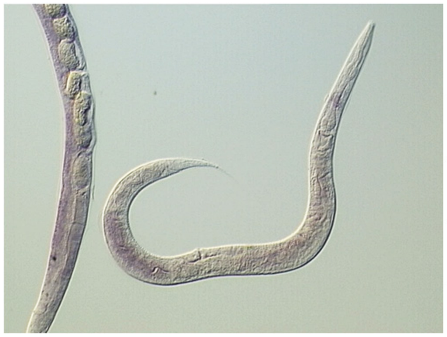

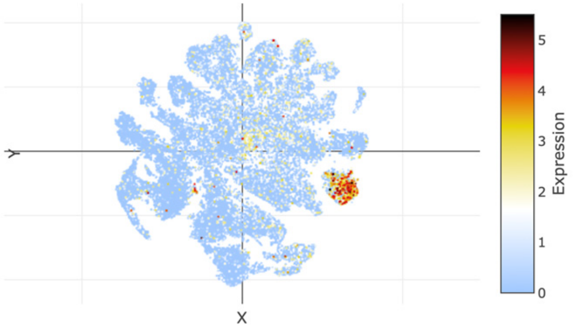

### t-SNE showing cell type 11

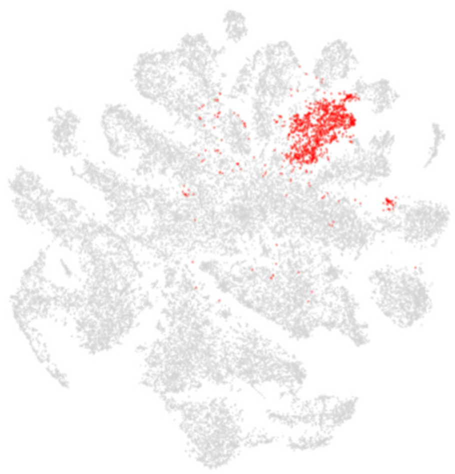

### Top 20 enriched genes

| | p_val | avg_log2FC | pct.1 | pct.2 | p_val_adj |
|---|---|---|---|---|---|
| *pdf-1* | 1.14E-263 | 3.15792657 | 0.493 | 0.109 | 5.37E-259 |
| *flp-11* | 9.30E-97 | 2.88892322 | 0.22 | 0.05 | 4.36E-92 |
| *pgal-1* | 0 | 2.65560862 | 0.637 | 0.136 | 0 |
| *sbt-1* | 0 | 2.59544926 | 0.743 | 0.204 | 0 |
| *flp-1* | 2.89E-33 | 2.58795689 | 0.136 | 0.045 | 1.36E-28 |
| *egl-3* | 0 | 2.58001703 | 0.596 | 0.118 | 0 |
| *egl-21* | 0 | 2.57536604 | 0.707 | 0.195 | 0 |
| *flp-14* | 1.71E-166 | 2.55395872 | 0.425 | 0.116 | 8.04E-162 |
| *flp-9* | 1.12E-284 | 2.46279816 | 0.516 | 0.113 | 5.25E-280 |
| *nlp-6* | 1.69E-112 | 2.38301882 | 0.213 | 0.043 | 7.92E-108 |
| *pghm-1* | 0 | 2.35901892 | 0.575 | 0.104 | 0 |
| *nlp-21* | 1.13E-295 | 2.35357859 | 0.449 | 0.081 | 5.28E-291 |
| *ida-1* | 0 | 2.3037682 | 0.54 | 0.091 | 0 |
| *T27C4.1* | 0 | 2.2822674 | 0.602 | 0.121 | 0 |
| *nlp-49* | 6.03E-82 | 2.19388208 | 0.199 | 0.048 | 2.83E-77 |
| *flp-19* | 8.36E-104 | 2.18585826 | 0.201 | 0.041 | 3.92E-99 |
| *flp-28* | 1.09E-59 | 2.11679099 | 0.294 | 0.113 | 5.11E-55 |
| *cla-1* | 3.17E-258 | 2.1069887 | 0.304 | 0.041 | 1.49E-253 |
| *flp-12* | 5.62E-77 | 2.08639834 | 0.245 | 0.072 | 2.64E-72 |
| *unc-25* | 1.54E-110 | 2.06484427 | 0.222 | 0.047 | 7.24E-106 |

### Top 20 depleted genes

| | p_val | avg_log2FC | pct.1 | pct.2 | p_val_adj |
|---|---|---|---|---|---|
| *dig-1* | 8.14E-21 | -1.7349397 | 0.375 | 0.231 | 3.82E-16 |
| *pos-1* | 3.06E-12 | -1.1794904 | 0.119 | 0.214 | 1.43E-07 |
| *clec-87* | 4.19E-08 | -1.1481222 | 0.297 | 0.37 | 0.0019661 |
| *Y105E8A.25* | 7.81E-16 | -1.0939968 | 0.23 | 0.35 | 3.66E-11 |
| *spn-4* | 5.96E-10 | -1.0274175 | 0.139 | 0.224 | 2.80E-05 |
| *C05C10.5* | 1.48E-07 | -0.9652952 | 0.113 | 0.18 | 0.00691876 |
| *ipla-1* | 1.86E-09 | -0.9564141 | 0.272 | 0.352 | 8.70E-05 |
| *plg-1* | 7.81E-09 | -0.949854 | 0.153 | 0.228 | 0.00036647 |
| *afd-1* | 5.70E-12 | -0.9290684 | 0.242 | 0.34 | 2.68E-07 |
| *hil-5* | 1.75E-10 | -0.898671 | 0.149 | 0.238 | 8.19E-06 |
| *ced-1* | 1.74E-12 | -0.8920637 | 0.28 | 0.378 | 8.18E-08 |
| *npp-8* | 7.59E-13 | -0.8548046 | 0.138 | 0.237 | 3.56E-08 |
| *patr-1* | 6.23E-07 | -0.8203308 | 0.139 | 0.204 | 0.02923966 |
| *F53H1.4* | 8.71E-12 | -0.81501 | 0.139 | 0.234 | 4.09E-07 |
| *C23H5.8* | 3.11E-07 | -0.8111844 | 0.396 | 0.455 | 0.01457872 |
| *vit-6* | 2.87E-09 | -0.8095996 | 0.752 | 0.755 | 0.00013463 |
| *vit-2* | 1.27E-09 | -0.8023571 | 0.699 | 0.725 | 5.97E-05 |
| *vit-3* | 1.68E-11 | -0.8020149 | 0.669 | 0.714 | 7.87E-07 |
| *F57F5.1* | 1.61E-08 | -0.788964 | 0.439 | 0.502 | 0.00075311 |
| *vit-5* | 1.95E-09 | -0.786694 | 0.757 | 0.764 | 9.13E-05 |

### Tissue enrichment

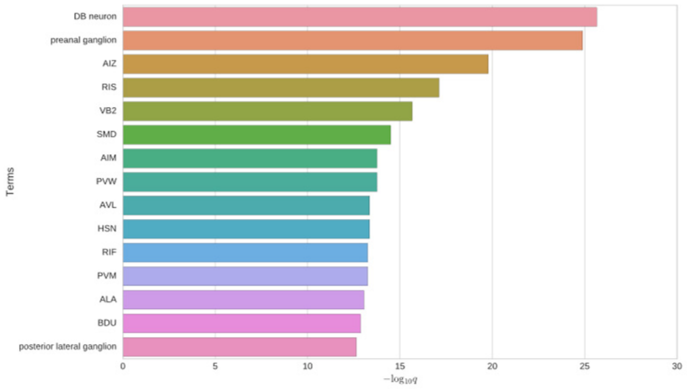


### GO enrichment

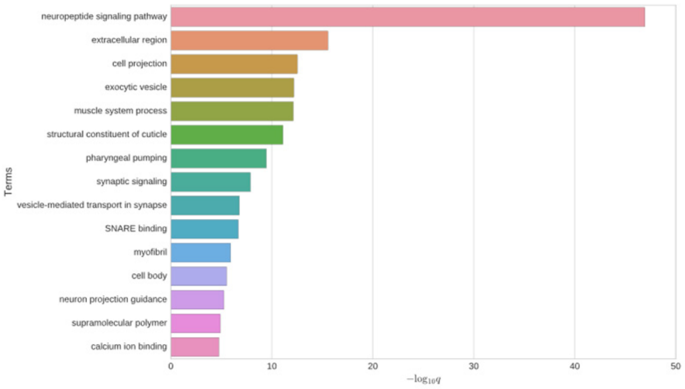


### Phenotype enrichment

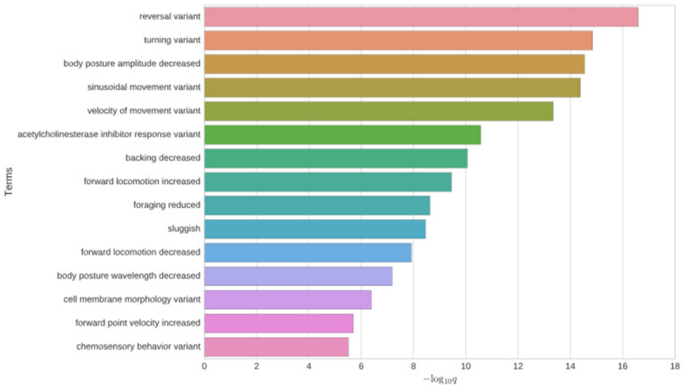


### *egl-3*

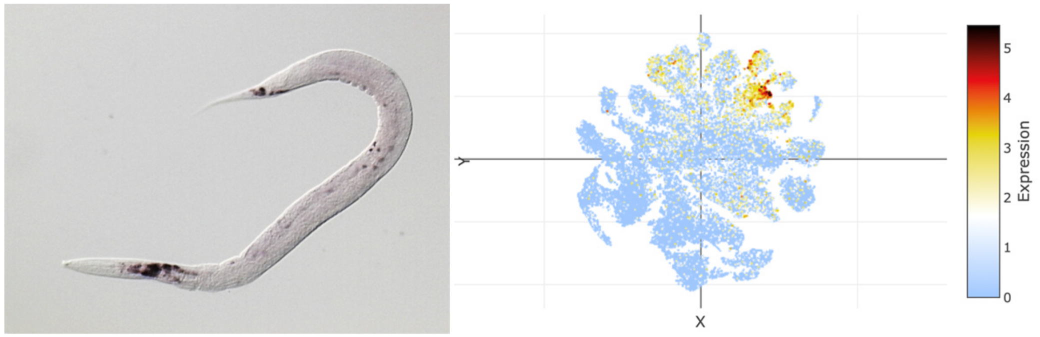


### *pgal-1*

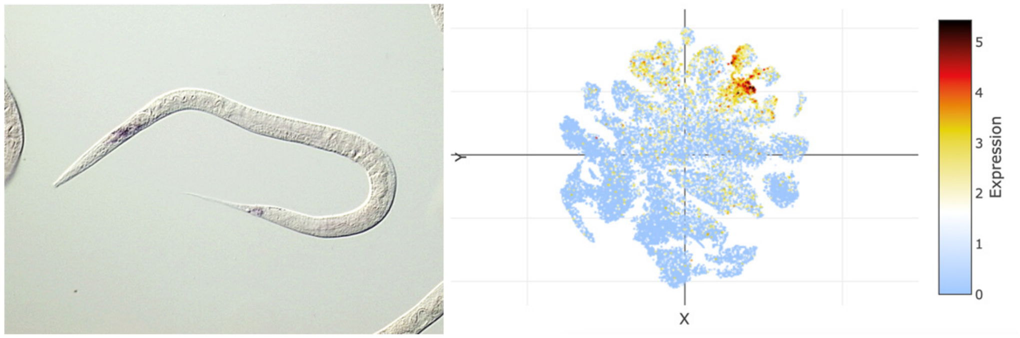

### t-SNE showing cell type 12

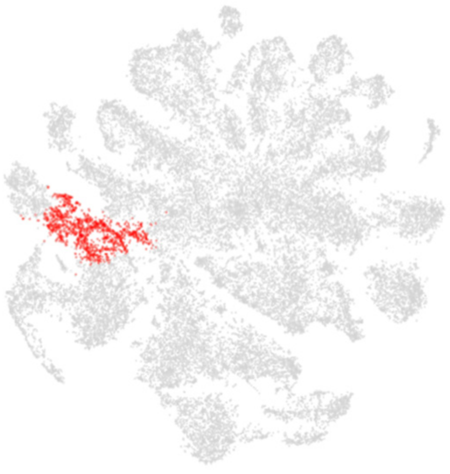

### Top 20 enriched genes

| | p_val | avg_log2FC | pct.1 | pct.2 | p_val_adj |
|---|---|---|---|---|---|
| *T21C9.13* | 4.82E-208 | 2.4510172 | 0.575 | 0.141 | 2.26E-203 |
| *clec-87* | 6.98E-187 | 2.38901326 | 0.805 | 0.354 | 3.27E-182 |
| *pos-1* | 3.32E-210 | 2.33671855 | 0.678 | 0.197 | 1.56E-205 |
| *C05C10.5* | 2.87E-183 | 2.30531624 | 0.586 | 0.166 | 1.35E-178 |
| *spn-4* | 4.68E-199 | 2.1976582 | 0.678 | 0.208 | 2.20E-194 |
| *F14H3.6* | 4.56E-194 | 2.09675757 | 0.483 | 0.104 | 2.14E-189 |
| *cpg-2* | 1.87E-134 | 2.00619924 | 0.751 | 0.369 | 8.76E-130 |
| *cpg-1* | 1.64E-134 | 2.00164692 | 0.707 | 0.315 | 7.71E-130 |
| *gyg-2* | 3.77E-167 | 1.99890985 | 0.625 | 0.206 | 1.77E-162 |
| *cey-3* | 3.90E-160 | 1.99649716 | 0.688 | 0.261 | 1.83E-155 |
| *patr-1* | 1.26E-119 | 1.98802883 | 0.537 | 0.192 | 5.91E-115 |
| *chs-1* | 4.11E-147 | 1.97655156 | 0.597 | 0.203 | 1.93E-142 |
| *hil-5* | 3.66E-167 | 1.97308027 | 0.653 | 0.222 | 1.72E-162 |
| *mex-5* | 5.85E-151 | 1.94497216 | 0.664 | 0.254 | 2.74E-146 |
| *oma-2* | 2.80E-161 | 1.93810121 | 0.639 | 0.22 | 1.31E-156 |
| *puf-5* | 5.57E-158 | 1.93576749 | 0.649 | 0.23 | 2.61E-153 |
| *cyb-3* | 1.18E-162 | 1.89649281 | 0.632 | 0.208 | 5.55E-158 |
| *cbd-1* | 1.99E-137 | 1.8950696 | 0.722 | 0.321 | 9.33E-133 |
| *hil-4* | 1.58E-158 | 1.87890268 | 0.739 | 0.299 | 7.43E-154 |
| *era-1* | 4.78E-133 | 1.87052723 | 0.48 | 0.14 | 2.24E-128 |

### Top 20 depleted genes

| | p_val | avg_log2FC | pct.1 | pct.2 | p_val_adj |
|---|---|---|---|---|---|
| *dig-1* | 3.36E-26 | -3.5631672 | 0.056 | 0.242 | 1.57E-21 |
| *ttn-1* | 3.59E-64 | -3.0264135 | 0.119 | 0.479 | 1.68E-59 |
| *D1086.10* | 6.65E-33 | -2.8301514 | 0.163 | 0.399 | 3.12E-28 |
| *ule-4* | 1.26E-45 | -2.7514809 | 0.22 | 0.515 | 5.90E-41 |
| *act-4* | 6.95E-115 | -2.4636351 | 0.273 | 0.731 | 3.26E-110 |
| *nlp-77* | 2.04E-100 | -2.4192281 | 0.312 | 0.715 | 9.55E-96 |
| *C10G8.4* | 8.07E-25 | -2.4138488 | 0.149 | 0.349 | 3.78E-20 |
| *C30E1.9* | 1.81E-65 | -2.3884692 | 0.127 | 0.493 | 8.48E-61 |
| *ule-2* | 6.93E-28 | -2.3755741 | 0.093 | 0.299 | 3.25E-23 |
| *ule-3* | 7.81E-10 | -2.3588929 | 0.139 | 0.245 | 3.66E-05 |
| *col-95* | 3.55E-66 | -2.3486214 | 0.244 | 0.598 | 1.66E-61 |
| *ttr-16* | 1.59E-89 | -2.3221053 | 0.239 | 0.656 | 7.44E-85 |
| *D1054.10* | 1.39E-13 | -2.2894495 | 0.186 | 0.325 | 6.50E-09 |
| *unc-54* | 1.71E-90 | -2.2856174 | 0.3 | 0.694 | 8.00E-86 |
| *ZC513.7* | 9.11E-36 | -2.2806017 | 0.185 | 0.436 | 4.27E-31 |
| *pat-10* | 7.01E-97 | -2.2675325 | 0.303 | 0.718 | 3.29E-92 |
| *anc-1* | 2.31E-100 | -2.2157306 | 0.251 | 0.702 | 1.08E-95 |
| *cpn-3* | 4.92E-88 | -2.1972421 | 0.308 | 0.692 | 2.31E-83 |
| *ttr-2* | 1.42E-46 | -2.1843719 | 0.176 | 0.471 | 6.67E-42 |
| *acp-6* | 2.93E-70 | -2.1766639 | 0.18 | 0.556 | 1.37E-65 |

### Tissue enrichment

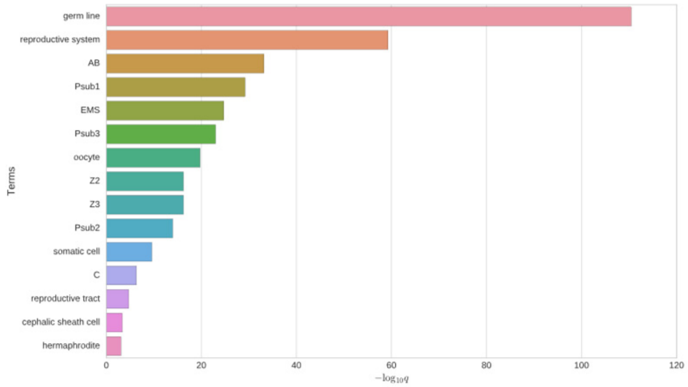


### GO enrichment

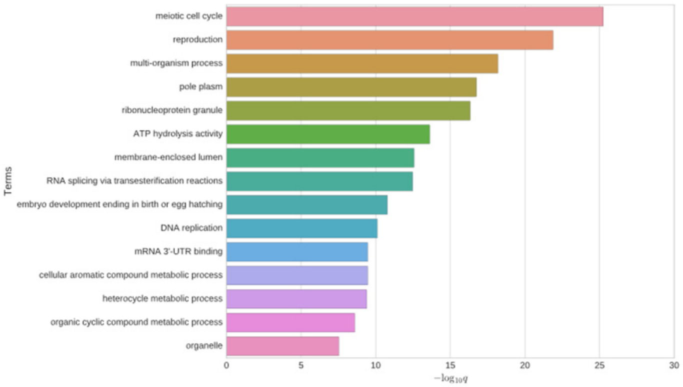


### Phenotype enrichment

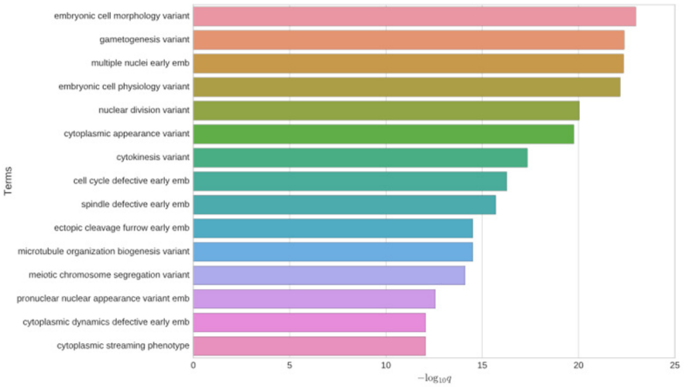


### *pos-1*

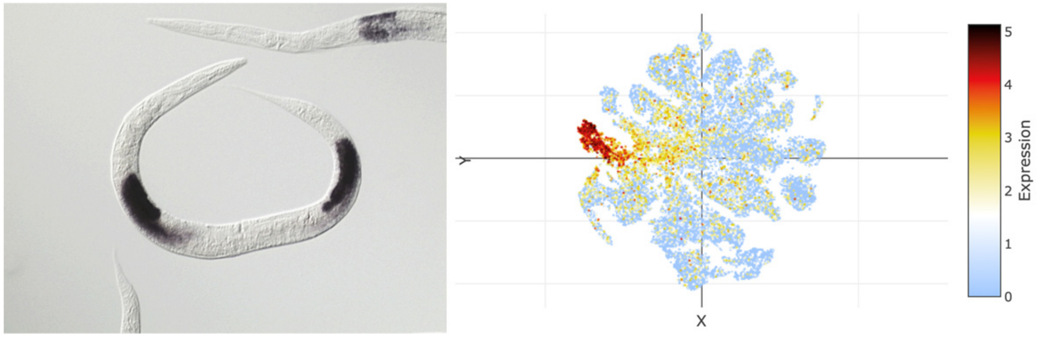


### *era-1*

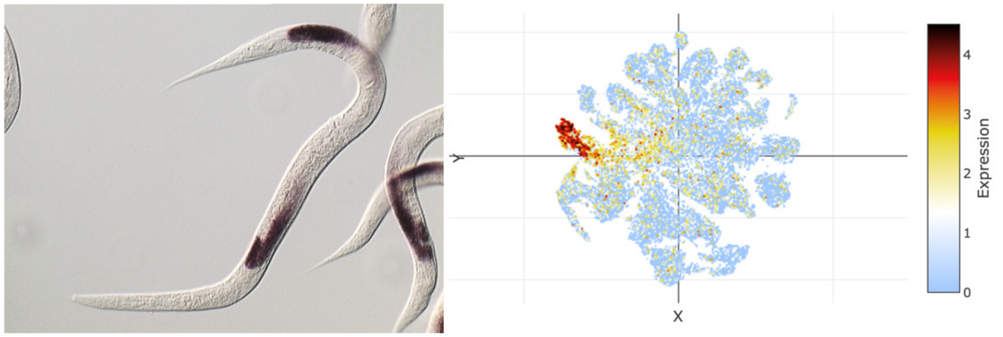

### t-SNE showing cell type 13

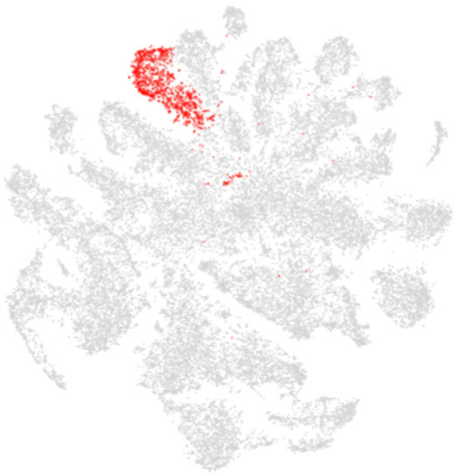

### Top 20 enriched genes

| | p_val | avg_log2FC | pct.1 | pct.2 | p_val_adj |
|---|---|---|---|---|---|
| *C49G7.3* | 0 | 4.76498013 | 0.911 | 0.157 | 0 |
| *T20G5.8* | 0 | 4.48679784 | 0.896 | 0.143 | 0 |
| *dod-6* | 0 | 4.32944692 | 0.94 | 0.26 | 0 |
| *C45G9.6* | 0 | 4.07980917 | 0.772 | 0.107 | 0 |
| *F15A4.6* | 0 | 4.06762594 | 0.826 | 0.107 | 0 |
| *scl-5* | 0 | 3.58833138 | 0.647 | 0.05 | 0 |
| *ZK596.1* | 0 | 3.51535406 | 0.703 | 0.06 | 0 |
| *fip-2* | 0 | 3.46578445 | 0.638 | 0.13 | 0 |
| *fipr-1* | 0 | 3.40325944 | 0.726 | 0.113 | 0 |
| *F41G3.10* | 0 | 3.29428648 | 0.665 | 0.055 | 0 |
| *fipr-5* | 0 | 3.20836232 | 0.502 | 0.075 | 0 |
| *msrp-7* | 0 | 3.15755235 | 0.587 | 0.04 | 0 |
| *ttr-21* | 0 | 2.97396129 | 0.674 | 0.099 | 0 |
| *R06F6.14* | 0 | 2.89970209 | 0.573 | 0.046 | 0 |
| *pqn-75* | 0 | 2.86196876 | 0.528 | 0.037 | 0 |
| *abf-5* | 0 | 2.76422695 | 0.556 | 0.06 | 0 |
| *myo-1* | 0 | 2.72050712 | 0.664 | 0.13 | 0 |
| *fipr-9* | 0 | 2.69876417 | 0.411 | 0.052 | 0 |
| *myo-2* | 0 | 2.68324963 | 0.645 | 0.133 | 0 |
| *ttr-26* | 2.25E-307 | 2.67652747 | 0.644 | 0.163 | 1.06E-302 |

### Top 20 depleted genes

| | p_val | avg_log2FC | pct.1 | pct.2 | p_val_adj |
|---|---|---|---|---|---|
| *mix-1* | 2.90E-17 | -1.260283 | 0.175 | 0.297 | 1.36E-12 |
| *afd-1* | 5.34E-15 | -1.2392402 | 0.23 | 0.341 | 2.51E-10 |
| *elpc-2* | 1.70E-16 | -1.131765 | 0.089 | 0.201 | 8.00E-12 |
| *F57F5.1* | 3.78E-15 | -1.1203366 | 0.403 | 0.503 | 1.77E-10 |
| *daf-2* | 1.37E-10 | -1.0981456 | 0.12 | 0.204 | 6.45E-06 |
| *sos-1* | 5.00E-13 | -1.0970272 | 0.136 | 0.235 | 2.34E-08 |
| *plg-1* | 1.12E-13 | -1.0809637 | 0.124 | 0.229 | 5.25E-09 |
| *lpd-3* | 2.40E-13 | -1.0697997 | 0.238 | 0.336 | 1.13E-08 |
| *npp-8* | 2.83E-21 | -1.0691297 | 0.099 | 0.238 | 1.33E-16 |
| *crh-1* | 4.52E-08 | -1.0595862 | 0.193 | 0.264 | 0.00211953 |
| *Y105E8A.25* | 4.47E-09 | -1.0572706 | 0.274 | 0.348 | 0.00020988 |
| *H06I04.3* | 2.59E-16 | -1.0357275 | 0.149 | 0.266 | 1.21E-11 |
| *mnk-1* | 5.56E-08 | -0.9994211 | 0.138 | 0.208 | 0.002606 |
| *F53H1.4* | 1.74E-17 | -0.9896699 | 0.111 | 0.235 | 8.18E-13 |
| *ceh-100* | 9.07E-15 | -0.9589538 | 0.134 | 0.244 | 4.26E-10 |
| *ipla-1* | 1.58E-09 | -0.9479123 | 0.272 | 0.352 | 7.40E-05 |
| *golg-4* | 5.87E-13 | -0.9314871 | 0.167 | 0.268 | 2.75E-08 |
| *aakg-1* | 4.62E-08 | -0.9294196 | 0.14 | 0.211 | 0.00216874 |
| *pos-1* | 6.68E-07 | -0.913956 | 0.143 | 0.213 | 0.03134372 |
| *klp-12* | 2.49E-11 | -0.9089067 | 0.292 | 0.377 | 1.17E-06 |

### Tissue enrichment

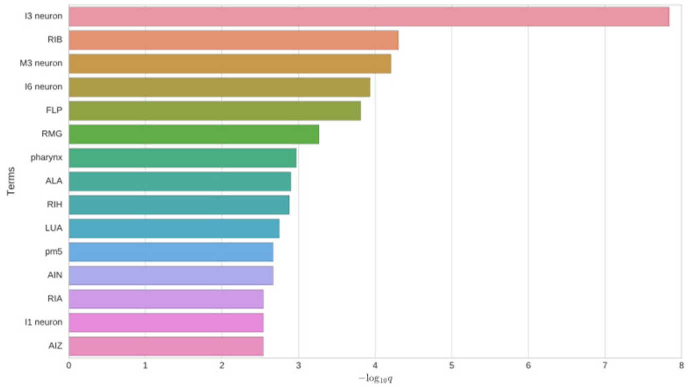


### GO enrichment

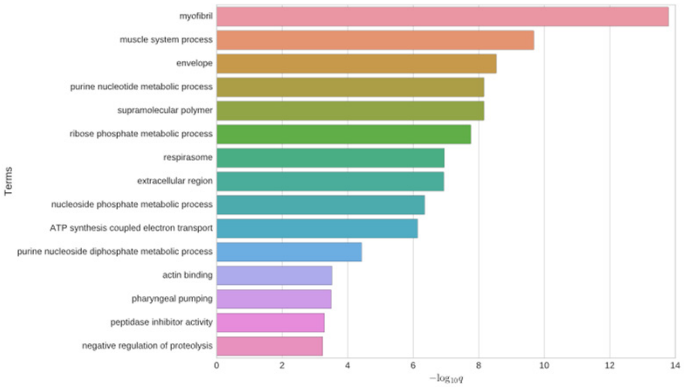


### Phenotype enrichment

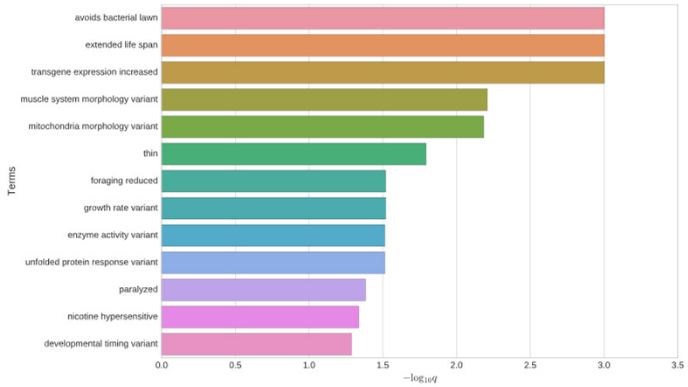


### *phat-3*

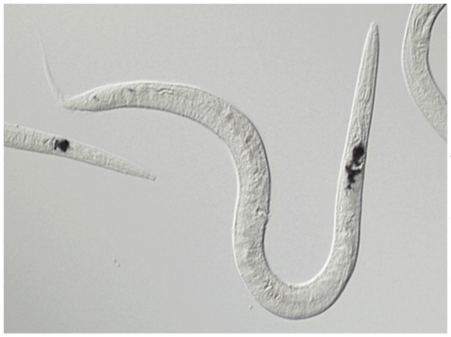

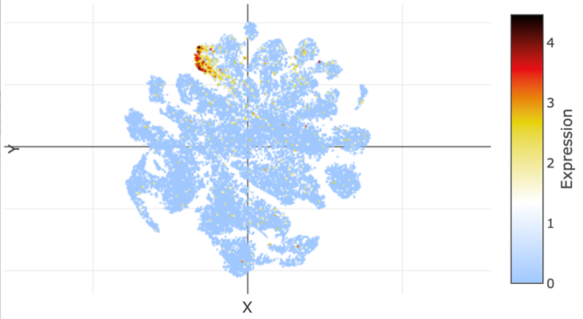


## *F15A4.6*

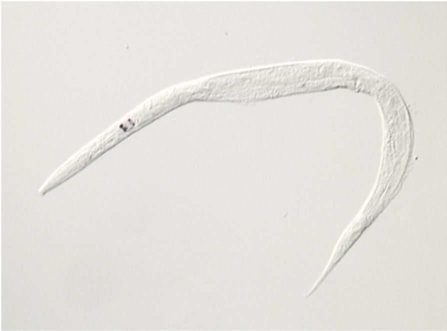

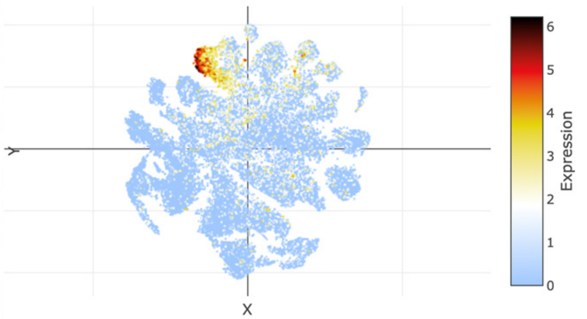

### t-SNE showing cell type 14

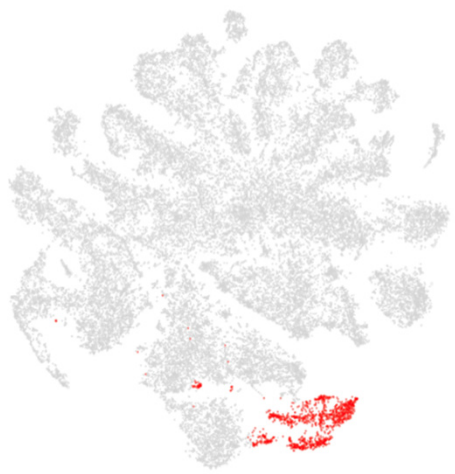

### Top 20 enriched genes

| | p_val | avg_log2FC | pct.1 | pct.2 | p_val_adj |
|---|---|---|---|---|---|
| *ddo-2* | 0 | 3.97541971 | 0.9 | 0.299 | 0 |
| *C52D10.3* | 0 | 3.92693037 | 0.646 | 0.073 | 0 |
| *jun-1* | 0 | 3.82675611 | 0.844 | 0.119 | 0 |
| *C23H5.8* | 0 | 3.34288879 | 0.965 | 0.436 | 0 |
| *ipla-1* | 7.61E-241 | 3.27978183 | 0.823 | 0.334 | 3.57E-236 |
| *fkh-7* | 0 | 3.11782258 | 0.749 | 0.166 | 0 |
| *F53H2.3* | 0 | 3.1177772 | 0.798 | 0.182 | 0 |
| *pals-24* | 0 | 3.08561474 | 0.476 | 0.051 | 0 |
| *fat-6* | 1.96E-229 | 2.92262566 | 0.912 | 0.539 | 9.19E-225 |
| *Y105C5A.15* | 0 | 2.91868665 | 0.737 | 0.116 | 0 |
| *F19B2.5* | 0 | 2.86278305 | 0.681 | 0.125 | 0 |
| *dhs-3* | 0 | 2.82789468 | 0.886 | 0.295 | 0 |
| *mnk-1* | 0 | 2.74638539 | 0.794 | 0.187 | 0 |
| *spsb-1* | 9.17E-271 | 2.70029077 | 0.806 | 0.272 | 4.30E-266 |
| *ZK228.4* | 0 | 2.68464437 | 0.555 | 0.071 | 0 |
| *Y105E8A.25* | 1.83E-273 | 2.66687103 | 0.854 | 0.329 | 8.60E-269 |
| *R193.2* | 6.47E-226 | 2.66437263 | 0.505 | 0.11 | 3.03E-221 |
| *math-27* | 0 | 2.65024879 | 0.695 | 0.038 | 0 |
| *ugt-29* | 0 | 2.63283628 | 0.598 | 0.086 | 0 |
| *ugt-26* | 5.54E-156 | 2.63214985 | 0.561 | 0.178 | 2.60E-151 |

### Top 20 depleted genes

| | p_val | avg_log2FC | pct.1 | pct.2 | p_val_adj |
|---|---|---|---|---|---|
| *dig-1* | 7.81E-27 | -3.4443488 | 0.063 | 0.243 | 3.66E-22 |
| *ule-4* | 1.72E-24 | -2.6602656 | 0.345 | 0.512 | 8.05E-20 |
| *ule-3* | 2.78E-09 | -2.6090091 | 0.154 | 0.245 | 0.00013018 |
| *ZK813.7* | 3.68E-13 | -2.5801459 | 0.116 | 0.226 | 1.72E-08 |
| *D1086.10* | 4.32E-18 | -2.5625027 | 0.257 | 0.397 | 2.02E-13 |
| *perm-4* | 1.77E-62 | -2.5587253 | 0.288 | 0.588 | 8.32E-58 |
| *ZC513.7* | 6.52E-48 | -2.4508518 | 0.166 | 0.438 | 3.06E-43 |
| *far-2* | 1.78E-105 | -2.4113742 | 0.504 | 0.774 | 8.35E-101 |
| *C10G8.4* | 1.49E-21 | -2.3593901 | 0.183 | 0.348 | 7.00E-17 |
| *C39D10.7* | 2.49E-14 | -2.3292463 | 0.165 | 0.287 | 1.17E-09 |
| *act-4* | 2.79E-101 | -2.2656725 | 0.405 | 0.728 | 1.31E-96 |
| *ZK813.3* | 3.74E-12 | -2.2340126 | 0.133 | 0.244 | 1.75E-07 |
| *Y62H9A.5* | 4.64E-08 | -2.2233081 | 0.2 | 0.284 | 0.00217622 |
| *far-1* | 1.92E-67 | -2.2072894 | 0.368 | 0.64 | 9.02E-63 |
| *ule-2* | 2.92E-19 | -2.1965398 | 0.146 | 0.298 | 1.37E-14 |
| *lbp-2* | 5.60E-62 | -2.1440925 | 0.401 | 0.639 | 2.62E-57 |
| *pat-10* | 1.32E-77 | -2.1339196 | 0.467 | 0.714 | 6.18E-73 |
| *ZK813.2* | 3.81E-20 | -2.1324268 | 0.142 | 0.294 | 1.79E-15 |
| *fasn-1* | 2.82E-22 | -2.1258283 | 0.117 | 0.281 | 1.32E-17 |
| *unc-27* | 1.33E-75 | -2.0966367 | 0.391 | 0.675 | 6.22E-71 |

### Tissue enrichment

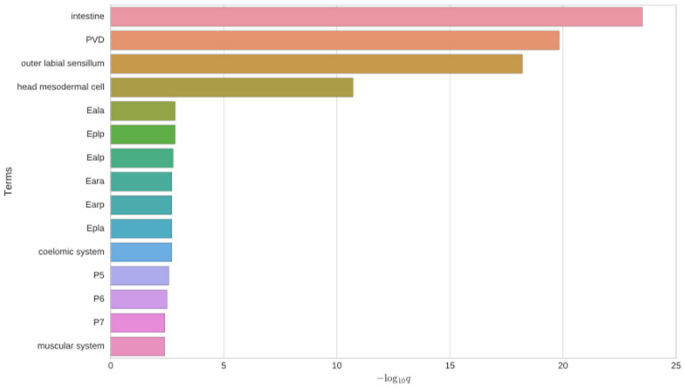


### GO enrichment

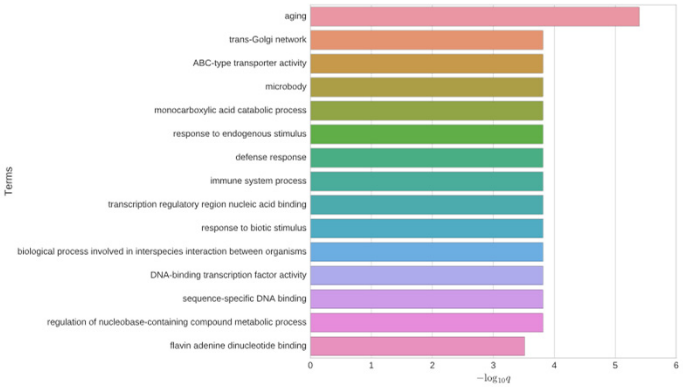


### Phenotype enrichment

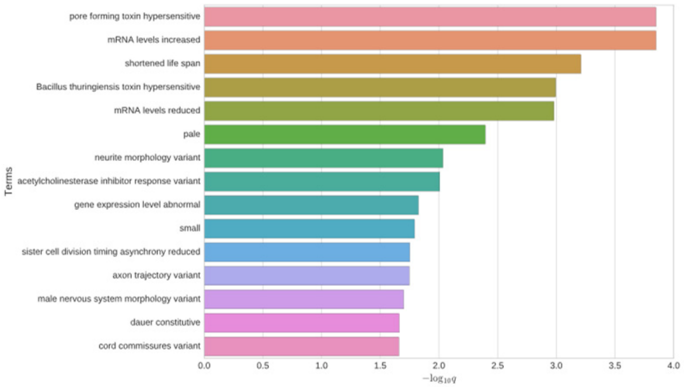


### *hmit-1.1*

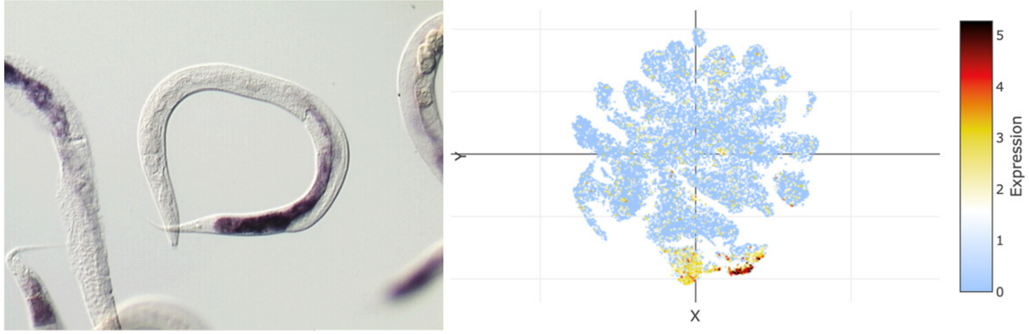


## *ZK228.4*

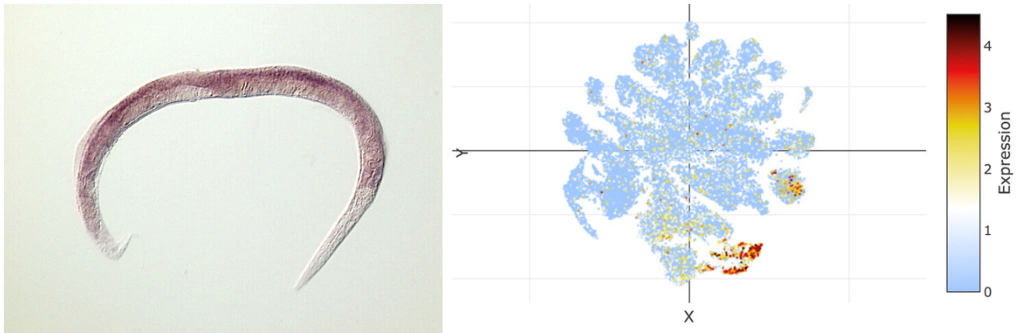

### t-SNE showing cell type 15

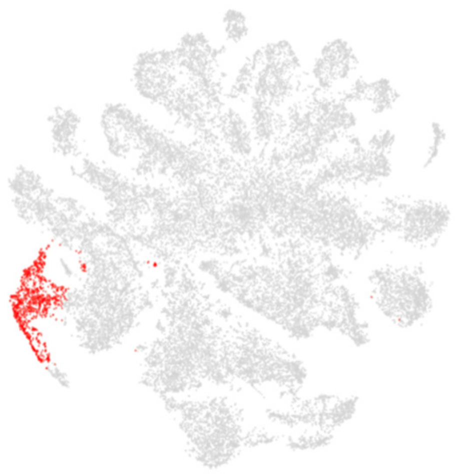

### Top 20 enriched genes

| | p_val | avg_log2FC | pct.1 | pct.2 | p_val_adj |
|---|---|---|---|---|---|
| *Y46H3C.7* | 0 | 6.15006879 | 0.93 | 0.099 | 0 |
| *Y9D1A.2* | 0 | 5.52199185 | 0.877 | 0.069 | 0 |
| *Y51H7C.15* | 0 | 4.42494134 | 0.757 | 0.033 | 0 |
| *F59C12.4* | 0 | 3.74340812 | 0.515 | 0.046 | 0 |
| *Y46H3C.5* | 0 | 2.92288937 | 0.508 | 0.019 | 0 |
| *elf-1* | 0 | 2.78345416 | 0.526 | 0.042 | 0 |
| *egl-1* | 0 | 2.38028235 | 0.371 | 0.037 | 0 |
| *mix-1* | 2.05E-268 | 2.30741333 | 0.81 | 0.277 | 9.63E-264 |
| *F29G9.1* | 0 | 2.25867831 | 0.392 | 0.01 | 0 |
| *spat-3* | 6.90E-237 | 2.1926356 | 0.569 | 0.132 | 3.24E-232 |
| *Y32H12A.8* | 1.42E-188 | 2.18849146 | 0.624 | 0.197 | 6.64E-184 |
| *Y9D1A.1* | 0 | 2.17054062 | 0.312 | 0.009 | 0 |
| *knl-2* | 1.22E-211 | 2.06610802 | 0.564 | 0.142 | 5.71E-207 |
| *afd-1* | 5.78E-195 | 2.00796476 | 0.786 | 0.323 | 2.71E-190 |
| *Y17D7C.3* | 2.57E-297 | 1.95416135 | 0.462 | 0.068 | 1.21E-292 |
| *ssl-1* | 1.86E-186 | 1.91506025 | 0.603 | 0.177 | 8.72E-182 |
| *F53H1.4* | 7.39E-187 | 1.9029219 | 0.658 | 0.217 | 3.47E-182 |
| *daf-19* | 1.68E-213 | 1.89315279 | 0.417 | 0.075 | 7.88E-209 |
| *xnp-1* | 6.57E-159 | 1.88359903 | 0.709 | 0.301 | 3.08E-154 |
| *uvs-1* | 1.28E-208 | 1.87663107 | 0.459 | 0.092 | 5.99E-204 |

### Top 20 depleted genes

| | p_val | avg_log2FC | pct.1 | pct.2 | p_val_adj |
|---|---|---|---|---|---|
| *pat-10* | 2.21E-137 | -3.091701 | 0.225 | 0.721 | 1.04E-132 |
| *ule-4* | 4.90E-47 | -3.0434865 | 0.24 | 0.515 | 2.30E-42 |
| *dig-1* | 1.18E-17 | -3.033376 | 0.093 | 0.242 | 5.53E-13 |
| *lev-11* | 1.94E-148 | -3.0173882 | 0.232 | 0.742 | 9.08E-144 |
| *D1086.10* | 9.02E-33 | -3.0136596 | 0.172 | 0.399 | 4.23E-28 |
| *act-4* | 9.93E-146 | -3.0041424 | 0.212 | 0.734 | 4.66E-141 |
| *far-2* | 3.40E-147 | -2.9022608 | 0.31 | 0.78 | 1.59E-142 |
| *unc-54* | 6.06E-119 | -2.8991705 | 0.237 | 0.696 | 2.84E-114 |
| *cpn-3* | 5.16E-122 | -2.8383992 | 0.239 | 0.695 | 2.42E-117 |
| *unc-27* | 7.07E-121 | -2.7958162 | 0.208 | 0.68 | 3.32E-116 |
| *nlp-77* | 1.23E-123 | -2.7903966 | 0.266 | 0.717 | 5.76E-119 |
| *mlc-3* | 4.08E-127 | -2.7852102 | 0.216 | 0.696 | 1.92E-122 |
| *ttr-16* | 9.06E-112 | -2.7733748 | 0.208 | 0.657 | 4.25E-107 |
| *F46H5.3* | 3.15E-161 | -2.7082589 | 0.33 | 0.815 | 1.48E-156 |
| *unc-15* | 4.65E-112 | -2.6995118 | 0.216 | 0.659 | 2.18E-107 |
| *col-119* | 1.85E-127 | -2.6820211 | 0.299 | 0.742 | 8.69E-123 |
| *col-181* | 4.10E-128 | -2.6567225 | 0.291 | 0.738 | 1.93E-123 |
| *Y37D8A.19* | 1.26E-25 | -2.6532443 | 0.168 | 0.362 | 5.90E-21 |
| *unc-87* | 5.37E-112 | -2.6204657 | 0.186 | 0.641 | 2.52E-107 |
| *ZK813.7* | 1.00E-16 | -2.6156693 | 0.09 | 0.227 | 4.70E-12 |

### Tissue enrichment

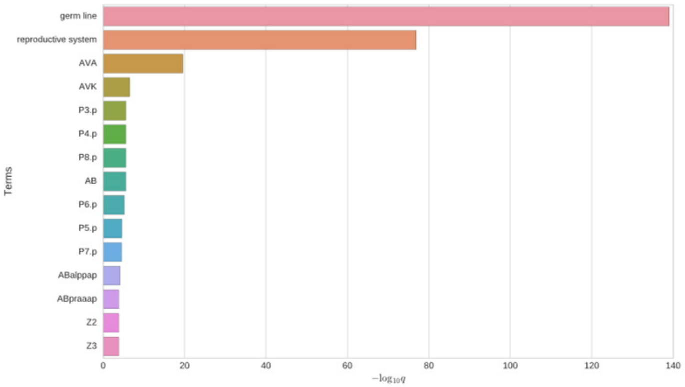


### GO enrichment

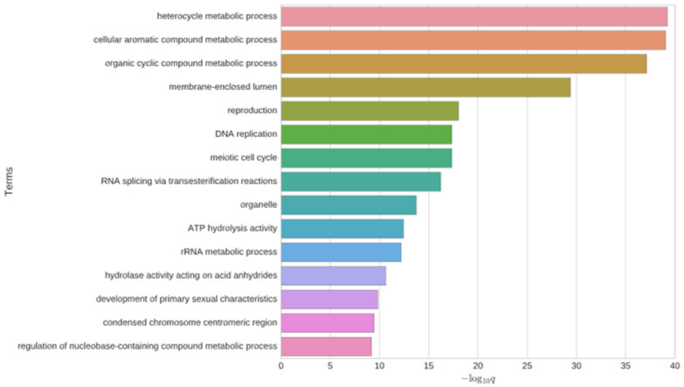


### Phenotype enrichment

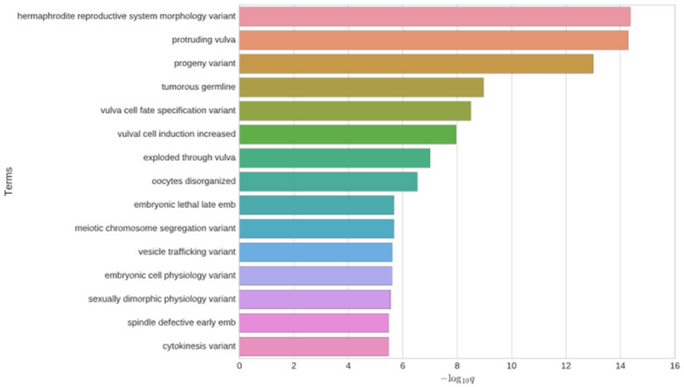


## *Y9D1A.1*

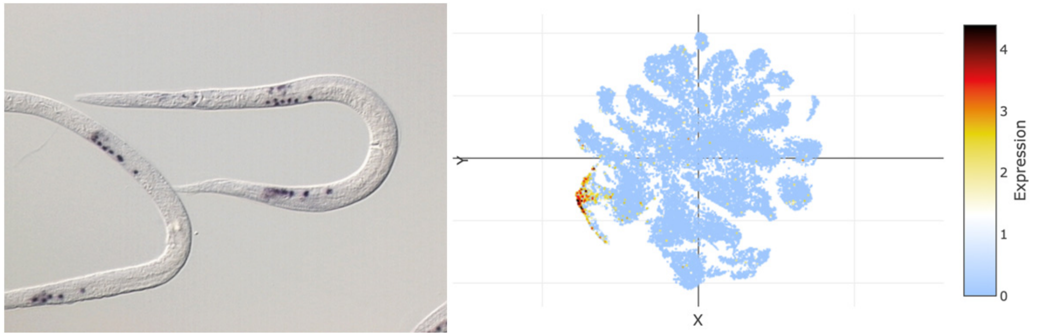


### *uvs-1*

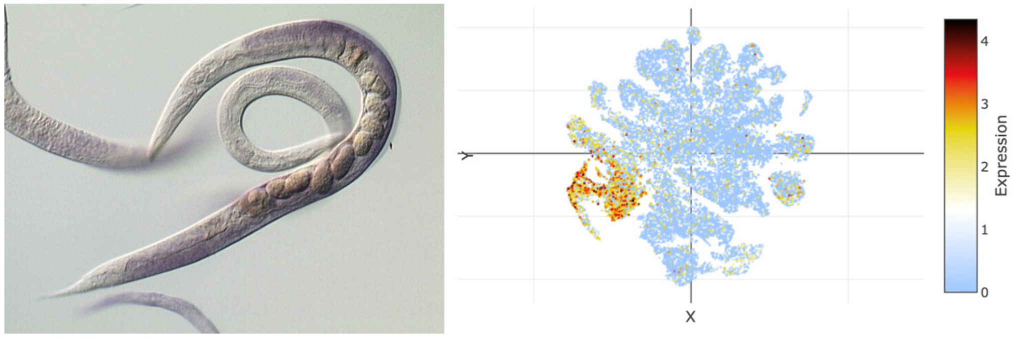

### t-SNE showing cell type 16

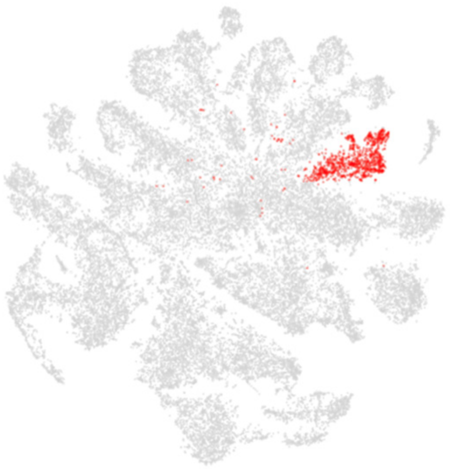

### Top 20 enriched genes

| | p_val | avg_log2FC | pct.1 | pct.2 | p_val_adj |
|---|---|---|---|---|---|
| *nlp-20* | 0 | 3.38587624 | 0.455 | 0.05 | 0 |
| *C02F5.14* | 0 | 3.05523726 | 0.554 | 0.077 | 0 |
| *pah-1* | 0 | 2.81978947 | 0.846 | 0.224 | 0 |
| *haly-1* | 0 | 2.67644923 | 0.752 | 0.17 | 0 |
| *Y51H4A.7* | 0 | 2.65518573 | 0.653 | 0.095 | 0 |
| *col-118* | 2.49E-173 | 2.58170994 | 0.53 | 0.145 | 1.17E-168 |
| *col-34* | 4.57E-269 | 2.51585999 | 0.455 | 0.072 | 2.14E-264 |
| *nlp-12* | 8.33E-165 | 2.30297016 | 0.287 | 0.044 | 3.91E-160 |
| *col-153* | 1.29E-248 | 2.3028854 | 0.371 | 0.051 | 6.07E-244 |
| *F55H12.4* | 8.07E-241 | 2.17031807 | 0.891 | 0.383 | 3.78E-236 |
| *mec-17* | 1.51E-111 | 2.15966739 | 0.282 | 0.06 | 7.08E-107 |
| *acp-6* | 4.69E-195 | 2.09288874 | 0.943 | 0.533 | 2.20E-190 |
| *F20A1.1* | 3.10E-142 | 2.0512075 | 0.548 | 0.174 | 1.45E-137 |
| *nlp-77* | 4.14E-227 | 2.01281701 | 0.997 | 0.694 | 1.94E-222 |
| *mec-7* | 3.48E-103 | 1.96252335 | 0.222 | 0.04 | 1.63E-98 |
| *Y47D7A.6* | 8.39E-155 | 1.95789023 | 0.264 | 0.04 | 3.94E-150 |
| *T23B12.8* | 7.87E-291 | 1.93687575 | 0.373 | 0.044 | 3.69E-286 |
| *Y46H3A.4* | 0 | 1.88512427 | 0.277 | 0.021 | 0 |
| *cnc-10* | 1.60E-195 | 1.8530118 | 0.389 | 0.069 | 7.50E-191 |
| *ttr-10* | 1.39E-303 | 1.83922593 | 0.361 | 0.039 | 6.50E-299 |

### Top 20 depleted genes

| | p_val | avg_log2FC | pct.1 | pct.2 | p_val_adj |
|---|---|---|---|---|---|
| *dig-1* | 9.70E-13 | -2.0217388 | 0.368 | 0.233 | 4.55E-08 |
| *F57F5.1* | 1.26E-29 | -1.4958071 | 0.301 | 0.505 | 5.89E-25 |
| *vit-5* | 5.26E-29 | -1.4383023 | 0.687 | 0.766 | 2.47E-24 |
| *vit-2* | 3.14E-30 | -1.4135271 | 0.598 | 0.728 | 1.47E-25 |
| *vit-3* | 2.31E-28 | -1.3640309 | 0.603 | 0.716 | 1.09E-23 |
| *vit-1* | 1.69E-26 | -1.3615896 | 0.559 | 0.679 | 7.92E-22 |
| *cpr-1* | 3.58E-19 | -1.3498833 | 0.305 | 0.457 | 1.68E-14 |
| *F53H2.3* | 2.88E-08 | -1.306804 | 0.123 | 0.203 | 0.00135151 |
| *T24B8.5* | 3.44E-09 | -1.2997323 | 0.318 | 0.411 | 0.00016118 |
| *lys-2* | 9.90E-21 | -1.2887652 | 0.326 | 0.478 | 4.64E-16 |
| *C23H5.8* | 1.65E-15 | -1.2802881 | 0.337 | 0.456 | 7.73E-11 |
| *lys-7* | 8.30E-07 | -1.2728533 | 0.251 | 0.328 | 0.03894275 |
| *mix-1* | 1.34E-13 | -1.2703497 | 0.177 | 0.296 | 6.30E-09 |
| *vit-4* | 4.83E-17 | -1.2605453 | 0.6 | 0.684 | 2.27E-12 |
| *Y105E8A.25* | 3.23E-09 | -1.2412627 | 0.266 | 0.347 | 0.00015146 |
| *vit-6* | 2.51E-22 | -1.23215 | 0.699 | 0.757 | 1.18E-17 |
| *daf-2* | 2.91E-13 | -1.2258175 | 0.092 | 0.204 | 1.36E-08 |
| *clec-87* | 4.45E-07 | -1.2085648 | 0.295 | 0.369 | 0.02087809 |
| *pos-1* | 5.60E-08 | -1.2041826 | 0.13 | 0.213 | 0.00262603 |
| *lys-1* | 2.76E-17 | -1.1659088 | 0.485 | 0.573 | 1.30E-12 |

### Tissue enrichment

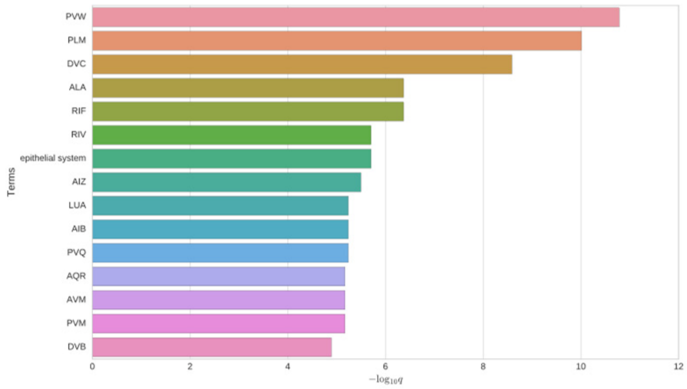


### GO enrichment

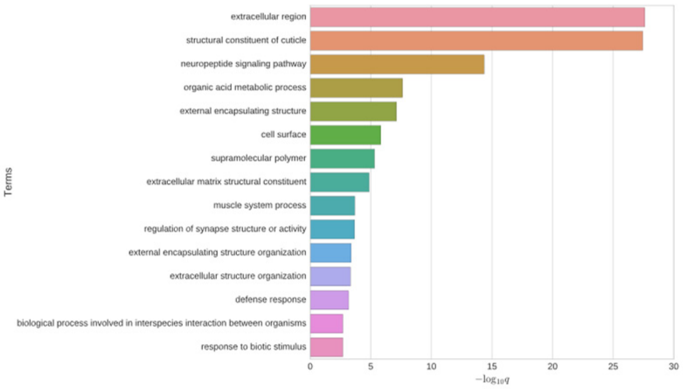


### Phenotype enrichment

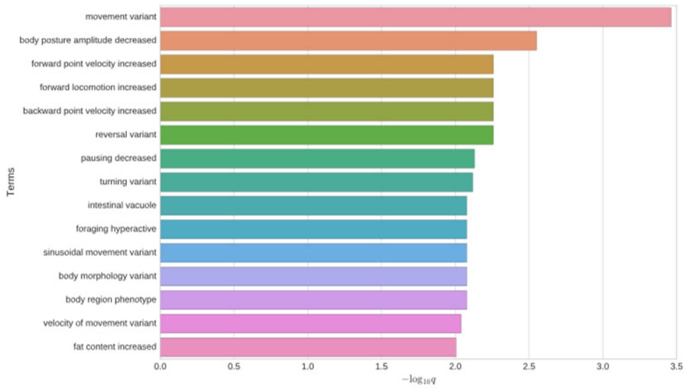


### *nlp-20*

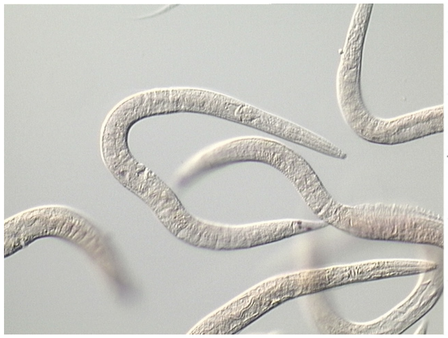

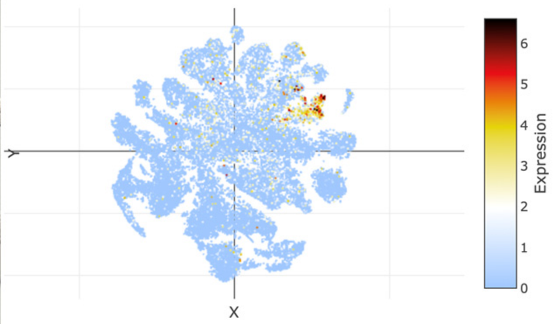


### *haly-1*

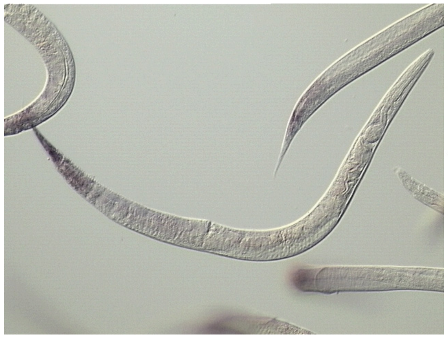

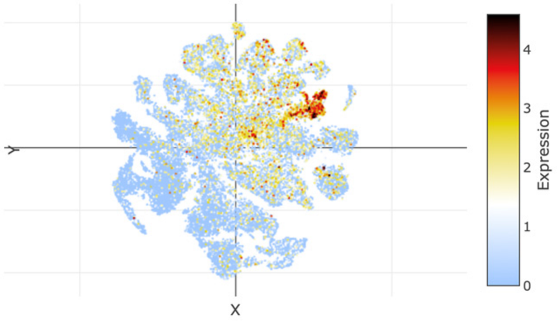

### t-SNE showing cell type 17

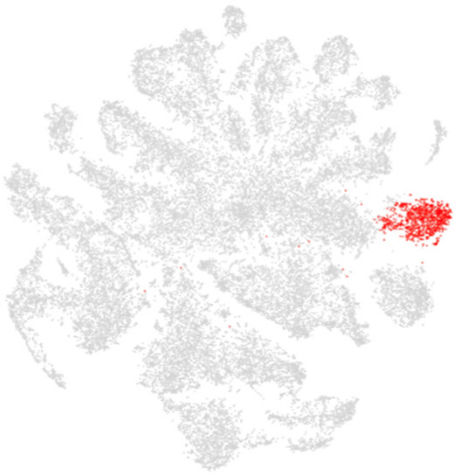

### Top 20 enriched genes

| | p_val | avg_log2FC | pct.1 | pct.2 | p_val_adj |
|---|---|---|---|---|---|
| *ttn-1* | 0 | 4.71043392 | 0.997 | 0.454 | 0 |
| *ndnf-1* | 0 | 3.86345479 | 0.668 | 0.08 | 0 |
| *pde-4* | 0 | 3.65561682 | 0.843 | 0.171 | 0 |
| *unc-22* | 0 | 3.49590736 | 0.933 | 0.323 | 0 |
| *cpna-2* | 1.20E-307 | 3.16785947 | 0.926 | 0.38 | 5.63E-303 |
| *M04G7.3* | 0 | 3.00944881 | 0.54 | 0.052 | 0 |
| *C32E12.4* | 0 | 2.82390468 | 0.68 | 0.137 | 0 |
| *unc-68* | 3.90E-264 | 2.6685098 | 0.811 | 0.281 | 1.83E-259 |
| *Y47H10A.4* | 0 | 2.653438 | 0.411 | 0.033 | 0 |
| *him-4* | 1.78E-145 | 2.609877 | 0.657 | 0.271 | 8.37E-141 |
| *unc-89* | 6.19E-256 | 2.59746611 | 0.917 | 0.424 | 2.90E-251 |
| *unc-49* | 0 | 2.59683984 | 0.539 | 0.074 | 0 |
| *unc-103* | 0 | 2.56775832 | 0.509 | 0.06 | 0 |
| *T13H5.1* | 5.27E-286 | 2.55410525 | 0.608 | 0.126 | 2.47E-281 |
| *prk-1* | 9.63E-206 | 2.55329927 | 0.647 | 0.196 | 4.52E-201 |
| *tnt-3* | 2.38E-190 | 2.55167952 | 0.695 | 0.246 | 1.12E-185 |
| *kin-1* | 1.83E-235 | 2.52008987 | 0.676 | 0.192 | 8.60E-231 |
| *C30E1.9* | 1.38E-243 | 2.499007 | 0.943 | 0.47 | 6.46E-239 |
| *W05F2.4* | 2.27E-242 | 2.4930021 | 0.651 | 0.174 | 1.06E-237 |
| *ketn-1* | 1.23E-192 | 2.44618878 | 0.842 | 0.397 | 5.77E-188 |

### Top 20 depleted genes

| | p_val | avg_log2FC | pct.1 | pct.2 | p_val_adj |
|---|---|---|---|---|---|
| *ule-2* | 6.54E-07 | -2.0044753 | 0.215 | 0.296 | 0.03067617 |
| *ZK813.2* | 3.80E-09 | -1.6817148 | 0.193 | 0.292 | 0.00017805 |
| *T20G5.8* | 2.78E-13 | -1.6650378 | 0.064 | 0.175 | 1.30E-08 |
| *dod-6* | 7.60E-12 | -1.6223984 | 0.165 | 0.289 | 3.56E-07 |
| *fipr-2* | 1.83E-08 | -1.6200748 | 0.11 | 0.197 | 0.00085739 |
| *nlp-77* | 6.99E-38 | -1.6092246 | 0.573 | 0.707 | 3.28E-33 |
| *C49G7.3* | 3.11E-09 | -1.603107 | 0.095 | 0.188 | 0.00014586 |
| *T21C9.13* | 4.32E-09 | -1.5734879 | 0.069 | 0.155 | 0.00020264 |
| *clec-87* | 6.65E-14 | -1.5470366 | 0.239 | 0.37 | 3.12E-09 |
| *Y22D7AR.10* | 1.32E-10 | -1.5246077 | 0.179 | 0.297 | 6.20E-06 |
| *ttr-2* | 1.06E-10 | -1.5077988 | 0.356 | 0.466 | 4.96E-06 |
| *ZC513.7* | 6.16E-07 | -1.4791371 | 0.339 | 0.432 | 0.02887184 |
| *myo-1* | 2.34E-08 | -1.3284302 | 0.072 | 0.153 | 0.00109781 |
| *acp-6* | 2.67E-18 | -1.3073254 | 0.422 | 0.549 | 1.25E-13 |
| *fipr-1* | 8.82E-10 | -1.296676 | 0.052 | 0.138 | 4.14E-05 |
| *spn-4* | 4.41E-15 | -1.274259 | 0.091 | 0.225 | 2.07E-10 |
| *col-95* | 8.96E-12 | -1.2364756 | 0.509 | 0.591 | 4.20E-07 |
| *C45G9.6* | 1.08E-07 | -1.2274516 | 0.06 | 0.134 | 0.00504972 |
| *pos-1* | 3.89E-11 | -1.2127919 | 0.103 | 0.213 | 1.82E-06 |
| *C05C10.5* | 1.02E-08 | -1.1876856 | 0.093 | 0.18 | 0.00047721 |

### Tissue enrichment

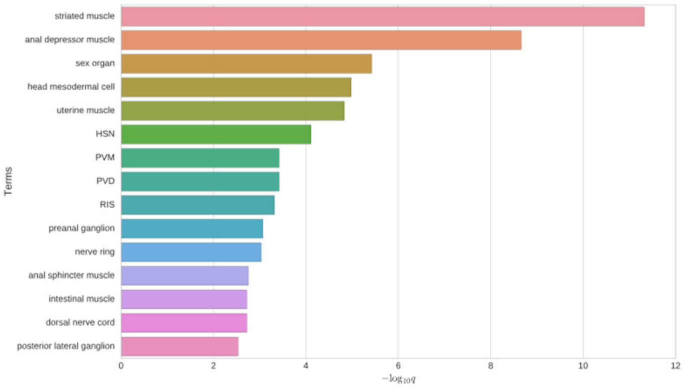


### GO enrichment

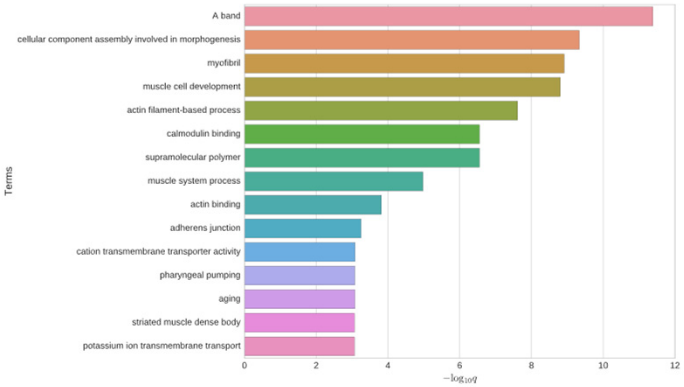


### Phenotype enrichment

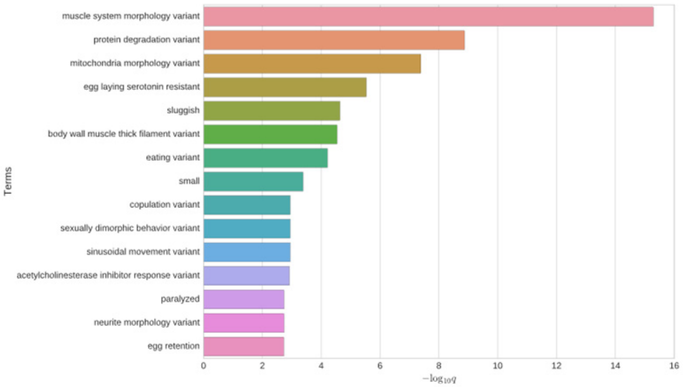


### *ndnf-1*

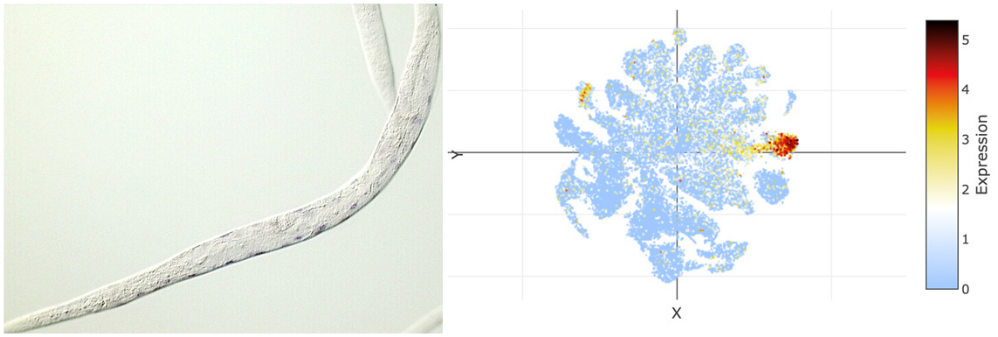


### *pde-4*

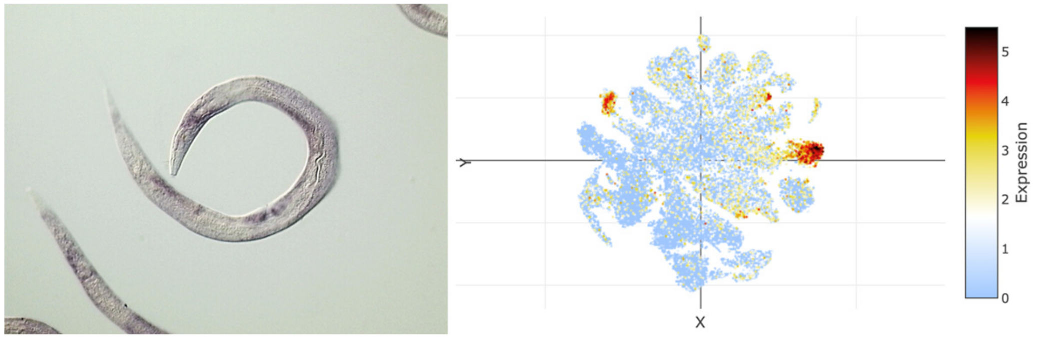

### t-SNE showing cell type 18

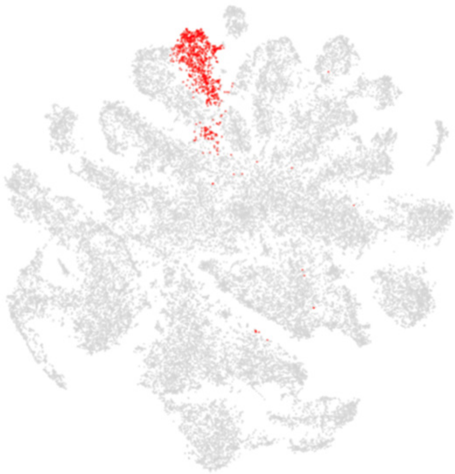

### Top 20 enriched genes

| | p_val | avg_log2FC | pct.1 | pct.2 | p_val_adj |
|---|---|---|---|---|---|
| *fipr-2* | 0 | 4.24298534 | 0.874 | 0.174 | 0 |
| *Y73F4A.2* | 0 | 4.08631311 | 0.779 | 0.083 | 0 |
| *myo-1* | 0 | 3.75760284 | 0.841 | 0.129 | 0 |
| *T03F1.11* | 0 | 3.7259048 | 0.886 | 0.143 | 0 |
| *myo-2* | 0 | 3.71661903 | 0.879 | 0.13 | 0 |
| *F41E6.15* | 0 | 3.64043414 | 0.818 | 0.122 | 0 |
| *tnt-4* | 0 | 3.38733634 | 0.739 | 0.072 | 0 |
| *C53B7.3* | 0 | 3.36929307 | 0.882 | 0.233 | 0 |
| *gpx-3* | 0 | 3.32405266 | 0.665 | 0.062 | 0 |
| *F35B12.3* | 0 | 3.19616538 | 0.775 | 0.089 | 0 |
| *hsp-12.2* | 0 | 3.19423322 | 0.787 | 0.13 | 0 |
| *tnc-2* | 0 | 3.13992367 | 0.76 | 0.104 | 0 |
| *tni-4* | 0 | 3.08797734 | 0.816 | 0.162 | 0 |
| *C45E5.4* | 0 | 3.02575579 | 0.464 | 0.035 | 0 |
| *abf-6* | 0 | 3.00805583 | 0.63 | 0.07 | 0 |
| *fipr-1* | 0 | 3.00647639 | 0.615 | 0.121 | 0 |
| *ttr-21* | 0 | 2.94070591 | 0.726 | 0.102 | 0 |
| *marg-1* | 0 | 2.90635956 | 0.51 | 0.033 | 0 |
| *ttr-27* | 0 | 2.90170531 | 0.66 | 0.077 | 0 |
| *myo-5* | 0 | 2.68708692 | 0.63 | 0.098 | 0 |

### Top 20 depleted genes

| | p_val | avg_log2FC | pct.1 | pct.2 | p_val_adj |
|---|---|---|---|---|---|
| *F53H2.3* | 3.64E-13 | -1.4527763 | 0.093 | 0.204 | 1.71E-08 |
| *afd-1* | 1.26E-14 | -1.2849631 | 0.211 | 0.34 | 5.92E-10 |
| *Y39B6A.1* | 8.86E-07 | -1.223081 | 0.354 | 0.432 | 0.04153365 |
| *Y105E8A.25* | 1.19E-14 | -1.2145742 | 0.222 | 0.349 | 5.60E-10 |
| *mix-1* | 2.12E-10 | -1.1284448 | 0.195 | 0.296 | 9.93E-06 |
| *F57F5.1* | 8.53E-16 | -1.1220463 | 0.382 | 0.503 | 4.00E-11 |
| *F53H1.4* | 2.90E-16 | -1.0922292 | 0.101 | 0.234 | 1.36E-11 |
| *vit-2* | 2.51E-11 | -1.0774401 | 0.726 | 0.724 | 1.18E-06 |
| *sos-1* | 5.85E-14 | -1.0709941 | 0.114 | 0.235 | 2.74E-09 |
| *daf-2* | 3.81E-08 | -1.0644315 | 0.124 | 0.203 | 0.00178525 |
| *spat-2* | 1.24E-15 | -1.0411305 | 0.193 | 0.325 | 5.80E-11 |
| *plg-1* | 6.50E-09 | -1.0209548 | 0.138 | 0.228 | 0.00030495 |
| *vit-5* | 5.91E-11 | -1.0201046 | 0.765 | 0.764 | 2.77E-06 |
| *crh-1* | 2.18E-09 | -1.013326 | 0.169 | 0.265 | 0.00010211 |
| *vit-6* | 3.36E-09 | -0.9862497 | 0.781 | 0.754 | 0.00015777 |
| *ced-1* | 2.84E-12 | -0.9835985 | 0.264 | 0.377 | 1.33E-07 |
| *lpd-3* | 3.27E-13 | -0.9772606 | 0.213 | 0.336 | 1.53E-08 |
| *apx-1* | 1.58E-10 | -0.960731 | 0.055 | 0.143 | 7.43E-06 |
| *npa-1* | 1.30E-16 | -0.9534892 | 0.502 | 0.596 | 6.09E-12 |
| *vit-3* | 5.24E-09 | -0.9480677 | 0.707 | 0.712 | 0.00024579 |

### Tissue enrichment

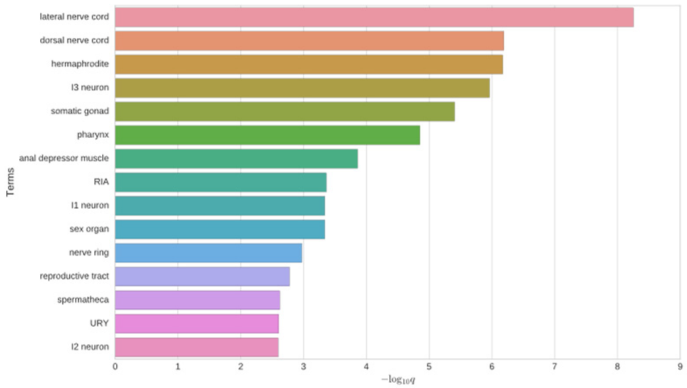


### GO enrichment

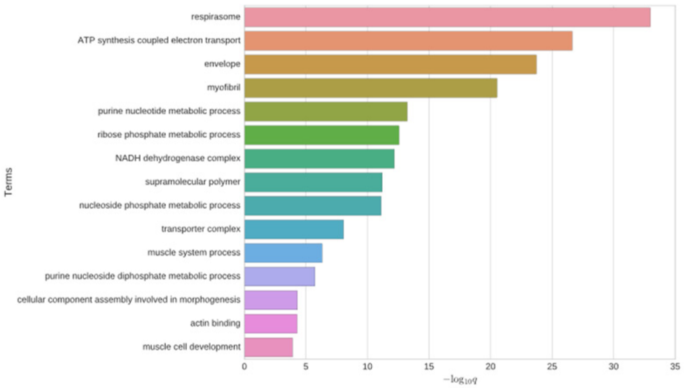


### Phenotype enrichment

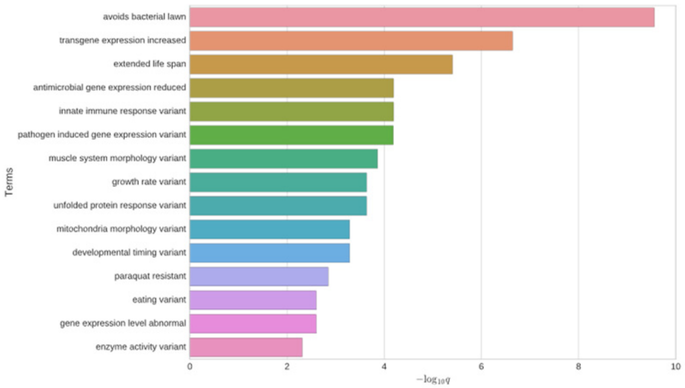


### *tnt-4*

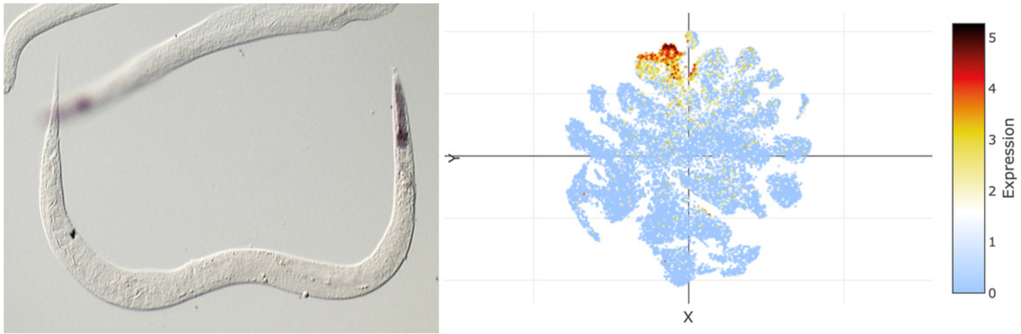


### *tni-4*

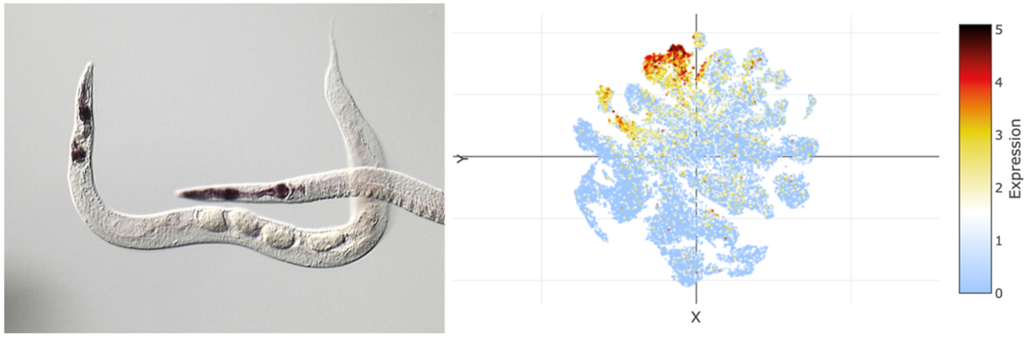

### t-SNE showing cell type 19

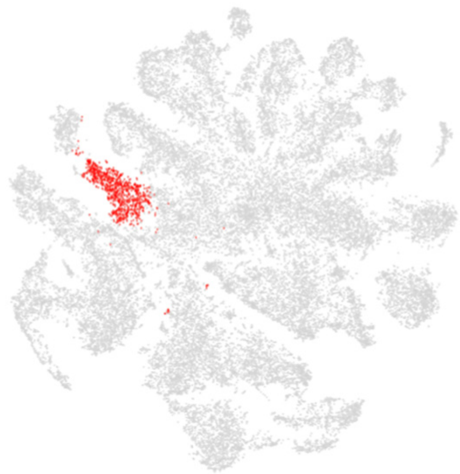

### Top 20 enriched genes

| | p_val | avg_log2FC | pct.1 | pct.2 | p_val_adj |
|---|---|---|---|---|---|
| *T04G9.7* | 0 | 3.89920734 | 0.971 | 0.365 | 0 |
| *perm-4* | 0 | 3.89342678 | 1 | 0.567 | 0 |
| *C39D10.7* | 0 | 3.73402906 | 0.977 | 0.264 | 0 |
| *tbh-1* | 0 | 3.70462896 | 0.923 | 0.199 | 0 |
| *C44B7.5* | 1.68E-157 | 3.59594683 | 0.71 | 0.296 | 7.90E-153 |
| *skpo-1* | 0 | 3.57449359 | 0.914 | 0.204 | 0 |
| *F48E3.4* | 0 | 3.54587511 | 0.908 | 0.228 | 0 |
| *B0513.4* | 0 | 3.3118748 | 0.847 | 0.207 | 0 |
| *perm-2* | 3.47E-285 | 3.17875693 | 0.998 | 0.687 | 1.63E-280 |
| *inx-8* | 0 | 3.17547075 | 0.784 | 0.149 | 0 |
| *sdz-27* | 0 | 3.16140111 | 0.86 | 0.245 | 0 |
| *ddo-3* | 0 | 2.78148639 | 0.692 | 0.125 | 0 |
| *fasn-1* | 2.90E-253 | 2.6233437 | 0.8 | 0.262 | 1.36E-248 |
| *F41C3.5* | 6.60E-220 | 2.57541678 | 0.941 | 0.524 | 3.10E-215 |
| *C17G1.2* | 0 | 2.54993391 | 0.644 | 0.081 | 0 |
| *cle-1* | 2.13E-252 | 2.46666479 | 0.674 | 0.168 | 9.99E-248 |
| *cht-3* | 1.97E-118 | 2.41511485 | 0.709 | 0.338 | 9.22E-114 |
| *W09C3.7* | 0 | 2.35509598 | 0.642 | 0.105 | 0 |
| *Y32F6A.5* | 6.30E-170 | 2.09865379 | 0.851 | 0.432 | 2.96E-165 |
| *Y38E10A.14* | 2.20E-129 | 2.08652542 | 0.628 | 0.232 | 1.03E-124 |

### Top 20 depleted genes

| | p_val | avg_log2FC | pct.1 | pct.2 | p_val_adj |
|---|---|---|---|---|---|
| *dig-1* | 9.17E-09 | -3.0593139 | 0.142 | 0.24 | 0.00043018 |
| *ttn-1* | 2.74E-10 | -1.7852621 | 0.387 | 0.471 | 1.29E-05 |
| *crh-1* | 3.58E-19 | -1.5974024 | 0.108 | 0.266 | 1.68E-14 |
| *spsb-1* | 1.01E-14 | -1.5258619 | 0.155 | 0.292 | 4.72E-10 |
| *C30E1.9* | 2.58E-15 | -1.4824084 | 0.347 | 0.487 | 1.21E-10 |
| *tos-1* | 2.42E-20 | -1.3574729 | 0.342 | 0.502 | 1.14E-15 |
| *ttr-2* | 1.64E-09 | -1.3464398 | 0.369 | 0.466 | 7.71E-05 |
| *col-12* | 4.00E-10 | -1.2316612 | 0.149 | 0.257 | 1.88E-05 |
| *ram-2* | 2.27E-09 | -1.2309732 | 0.158 | 0.261 | 0.00010626 |
| *acp-6* | 2.18E-13 | -1.2260347 | 0.451 | 0.548 | 1.02E-08 |
| *jun-1* | 6.83E-11 | -1.2147543 | 0.049 | 0.144 | 3.20E-06 |
| *daf-2* | 5.14E-11 | -1.1847657 | 0.099 | 0.203 | 2.41E-06 |
| *sos-1* | 5.17E-13 | -1.1688857 | 0.113 | 0.235 | 2.43E-08 |
| *fkh-7* | 3.06E-12 | -1.1639645 | 0.076 | 0.187 | 1.43E-07 |
| *T28D6.3* | 9.92E-08 | -1.1620953 | 0.151 | 0.238 | 0.00465501 |
| *F53H2.3* | 1.81E-07 | -1.1276952 | 0.121 | 0.203 | 0.00849378 |
| *nlp-77* | 5.26E-17 | -1.1017946 | 0.649 | 0.705 | 2.47E-12 |
| *ddo-2* | 4.36E-07 | -1.0950375 | 0.236 | 0.32 | 0.02043326 |
| *col-155* | 5.41E-07 | -1.0827854 | 0.122 | 0.201 | 0.0253796 |
| *T04F3.1* | 8.39E-12 | -1.0557274 | 0.183 | 0.312 | 3.93E-07 |

### Tissue enrichment

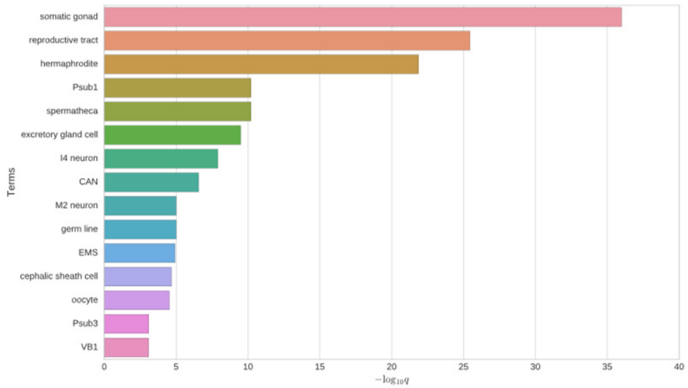


### GO enrichment

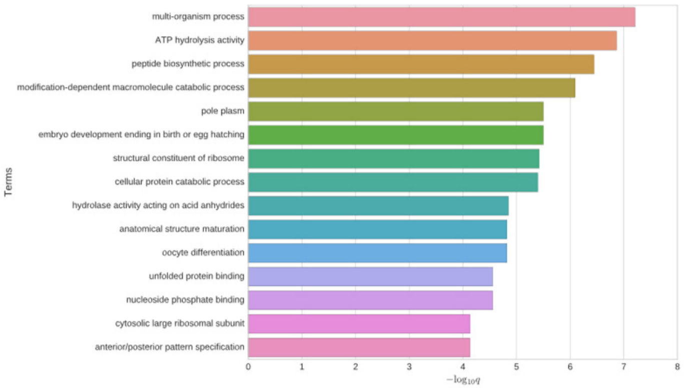


### Phenotype enrichment

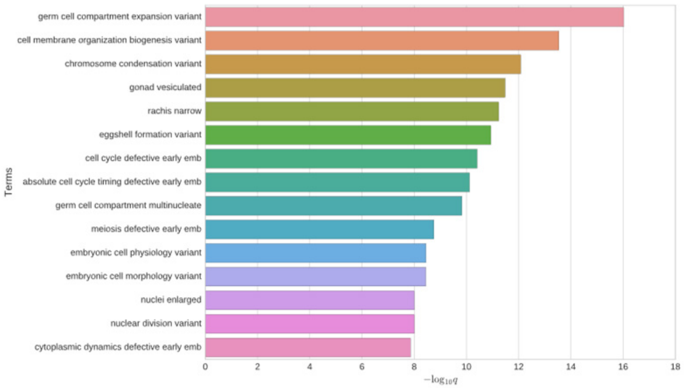


### *tbh-1*

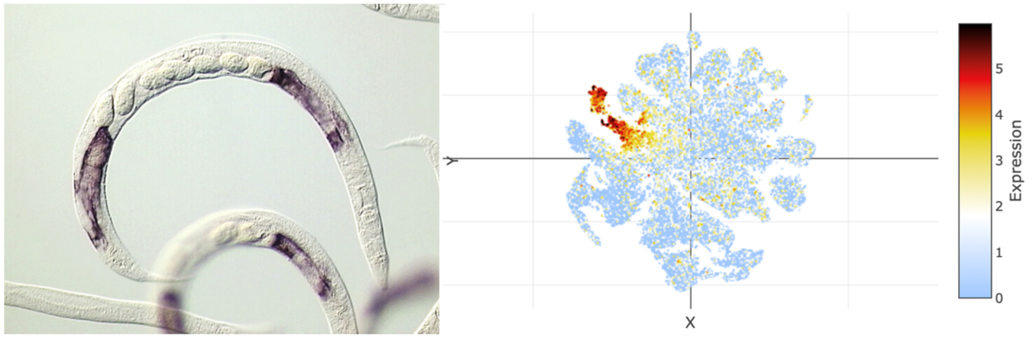


### *inx-8*

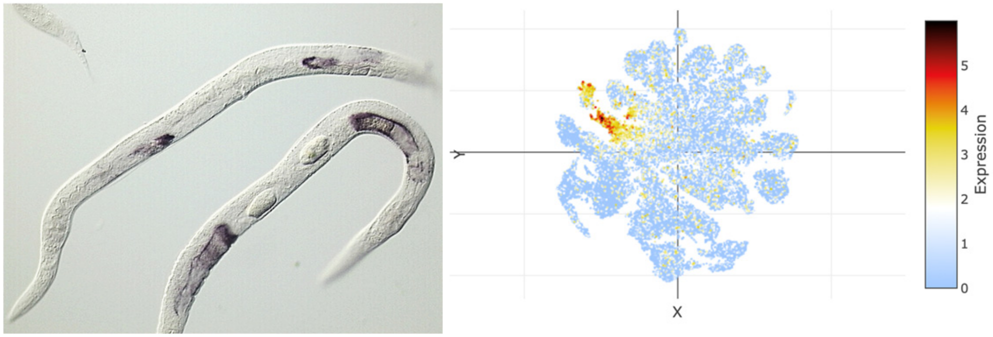

### t-SNE showing cell type 20

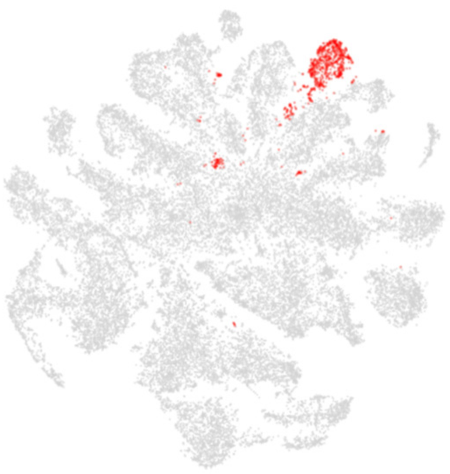

### Top 20 enriched genes

| | p_val | avg_log2FC | pct.1 | pct.2 | p_val_adj |
|---|---|---|---|---|---|
| *F53F4.13* | 0 | 4.0027797 | 0.922 | 0.182 | 0 |
| *F20A1.1* | 0 | 3.98221041 | 0.915 | 0.167 | 0 |
| *T02B11.3* | 0 | 3.82711022 | 0.899 | 0.131 | 0 |
| *far-8* | 0 | 3.53357811 | 0.813 | 0.103 | 0 |
| *F20A1.10* | 0 | 3.36043654 | 0.642 | 0.094 | 0 |
| *R11D1.3* | 0 | 3.25926067 | 0.732 | 0.065 | 0 |
| *F35B12.9* | 0 | 3.05471114 | 0.66 | 0.064 | 0 |
| *R102.2* | 6.63E-221 | 2.854265 | 0.547 | 0.109 | 3.11E-216 |
| *C33G8.4* | 0 | 2.80403476 | 0.668 | 0.076 | 0 |
| *F40F8.4* | 0 | 2.7879273 | 0.646 | 0.06 | 0 |
| *ZK822.4* | 0 | 2.59163118 | 0.62 | 0.057 | 0 |
| *Y69A2AR.22* | 0 | 2.58024445 | 0.606 | 0.052 | 0 |
| *F14D7.10* | 0 | 2.55719265 | 0.543 | 0.06 | 0 |
| *F59A7.2* | 0 | 2.52169293 | 0.547 | 0.055 | 0 |
| *tag-209* | 0 | 2.5049903 | 0.588 | 0.061 | 0 |
| *F14D7.7* | 2.28E-293 | 2.27685081 | 0.483 | 0.064 | 1.07E-288 |
| *K01A6.8* | 0 | 2.23840573 | 0.471 | 0.03 | 0 |
| *Y42G9A.2* | 0 | 2.16989085 | 0.487 | 0.04 | 0 |
| *F59A7.5* | 0 | 2.1248598 | 0.461 | 0.035 | 0 |
| *F07C6.3* | 0 | 2.08387698 | 0.479 | 0.031 | 0 |

### Top 20 depleted genes

| | p_val | avg_log2FC | pct.1 | pct.2 | p_val_adj |
|---|---|---|---|---|---|
| *dig-1* | 1.56E-10 | -1.9855564 | 0.368 | 0.234 | 7.31E-06 |
| *afd-1* | 1.95E-09 | -1.2546251 | 0.241 | 0.339 | 9.12E-05 |
| *Y105E8A.25* | 5.78E-08 | -1.1795545 | 0.264 | 0.347 | 0.00271109 |
| *mix-1* | 2.93E-09 | -1.1506557 | 0.193 | 0.295 | 0.00013747 |
| *sos-1* | 2.40E-09 | -1.1266752 | 0.133 | 0.234 | 0.0001126 |
| *F57F5.1* | 1.08E-10 | -1.1006425 | 0.404 | 0.502 | 5.08E-06 |
| *klp-12* | 4.49E-08 | -1.0225981 | 0.298 | 0.375 | 0.00210585 |
| *F53H1.4* | 3.64E-08 | -0.9829643 | 0.139 | 0.232 | 0.00170901 |
| *apx-1* | 1.23E-08 | -0.9389229 | 0.056 | 0.142 | 0.00057833 |
| *golg-4* | 5.56E-08 | -0.9294097 | 0.175 | 0.267 | 0.00260799 |
| *unc-40* | 8.38E-09 | -0.9106337 | 0.099 | 0.196 | 0.00039286 |
| *npp-8* | 2.10E-08 | -0.8990429 | 0.139 | 0.236 | 0.00098298 |
| *Y57A10A.31* | 3.38E-09 | -0.8771811 | 0.066 | 0.159 | 0.00015854 |
| *attf-6* | 1.69E-09 | -0.8768652 | 0.064 | 0.157 | 7.92E-05 |
| *xpc-1* | 3.91E-08 | -0.8762269 | 0.121 | 0.215 | 0.0018354 |
| *Y53F4B.21* | 3.61E-08 | -0.8582019 | 0.052 | 0.132 | 0.00169348 |
| *Y48G8AL.5* | 1.07E-07 | -0.8387539 | 0.074 | 0.156 | 0.00503567 |
| *ceh-100* | 5.20E-08 | -0.8343194 | 0.149 | 0.242 | 0.00243906 |
| *lea-1* | 6.27E-07 | -0.7774685 | 0.479 | 0.552 | 0.02942488 |
| *sqv-6* | 8.92E-07 | -0.7504222 | 0.038 | 0.102 | 0.04181791 |

### Tissue enrichment

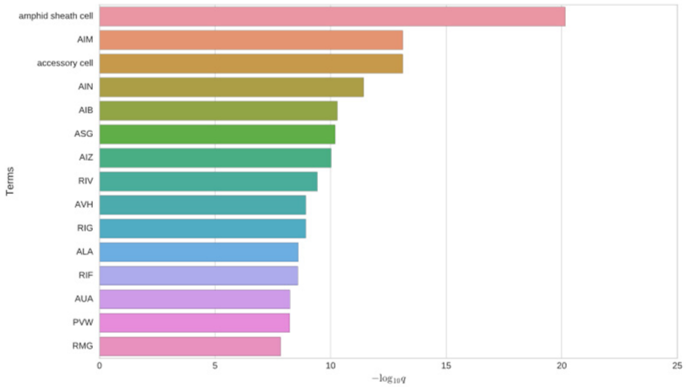


### GO enrichment

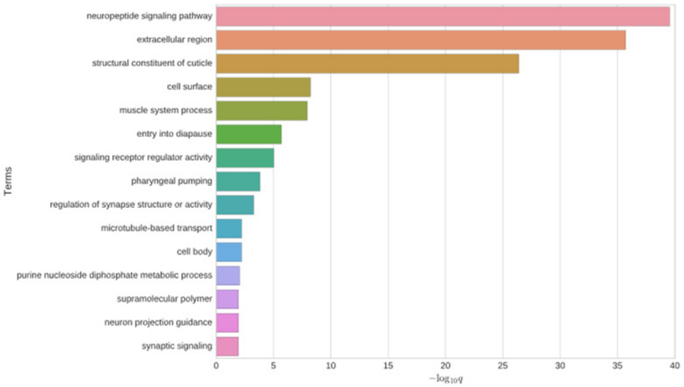


### Phenotype enrichment

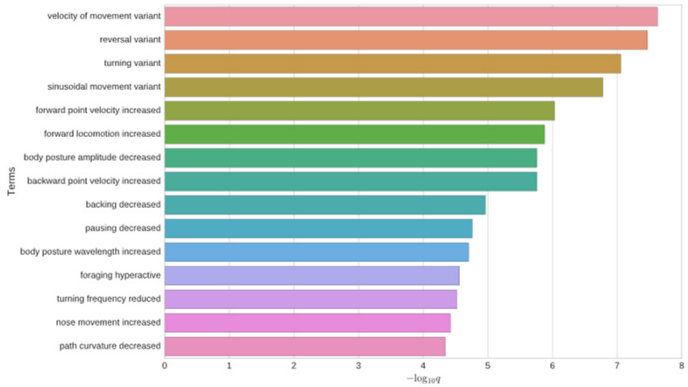


## *T02B11.3*

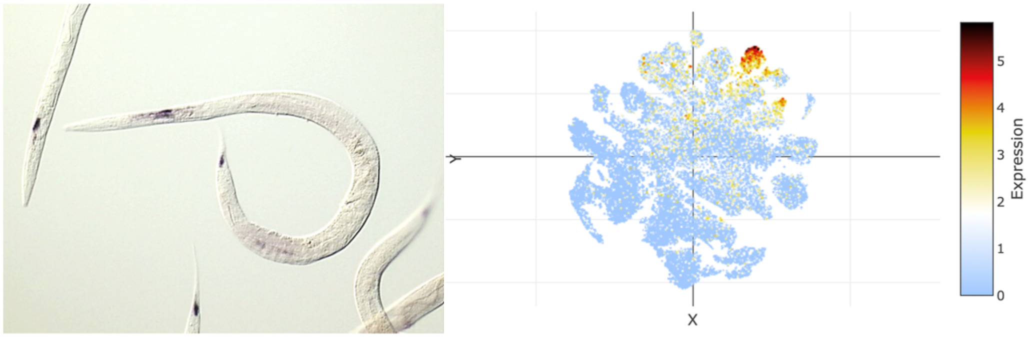


## *F53F4.13*

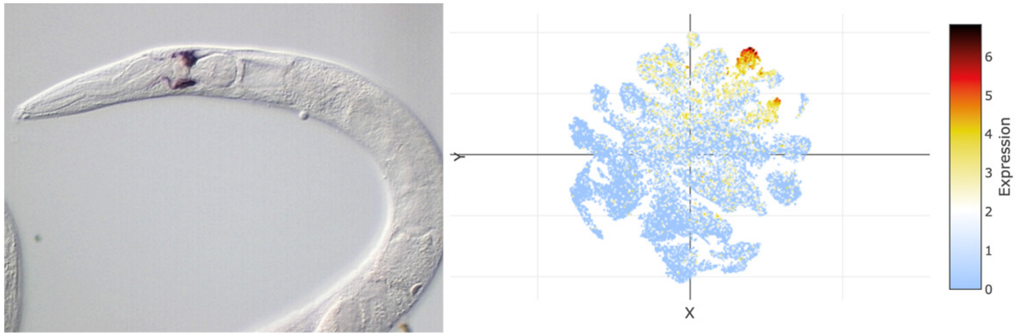

### t-SNE showing cell type 21

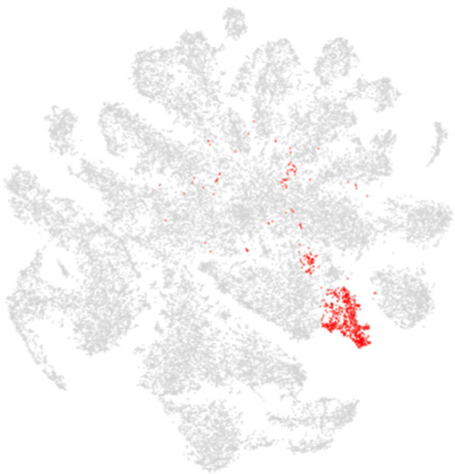

### Top 20 enriched genes

| | p_val | avg_log2FC | pct.1 | pct.2 | p_val_adj |
|---|---|---|---|---|---|
| *fipr-13* | 0 | 4.09801999 | 0.786 | 0.101 | 0 |
| *col-155* | 0 | 4.03167096 | 0.92 | 0.182 | 0 |
| *col-96* | 0 | 3.71423829 | 0.887 | 0.208 | 0 |
| *col-166* | 0 | 3.44090176 | 0.846 | 0.188 | 0 |
| *nas-37* | 0 | 3.30584965 | 0.639 | 0.058 | 0 |
| *col-118* | 0 | 3.26232385 | 0.743 | 0.142 | 0 |
| *hsp-43* | 1.90E-178 | 2.6812599 | 0.715 | 0.23 | 8.91E-174 |
| *Y105C5B.5* | 3.56E-173 | 2.53049404 | 0.893 | 0.434 | 1.67E-168 |
| *acp-6* | 4.27E-177 | 2.51770205 | 0.949 | 0.536 | 2.00E-172 |
| *F13D12.3* | 1.22E-184 | 2.35840308 | 0.63 | 0.17 | 5.72E-180 |
| *hsp-12.3* | 5.25E-106 | 2.12185877 | 0.439 | 0.123 | 2.46E-101 |
| *F42A8.1* | 6.73E-139 | 2.08036413 | 0.708 | 0.266 | 3.15E-134 |
| *Y43C5A.3* | 9.71E-129 | 2.05158344 | 0.528 | 0.155 | 4.55E-124 |
| *far-3* | 5.90E-82 | 2.00067362 | 0.563 | 0.227 | 2.77E-77 |
| *cht-3* | 1.23E-101 | 1.98297365 | 0.731 | 0.339 | 5.78E-97 |
| *F45E4.5* | 9.70E-164 | 1.96733219 | 0.357 | 0.058 | 4.55E-159 |
| *nlp-77* | 2.20E-160 | 1.91667965 | 0.99 | 0.696 | 1.03E-155 |
| *col-141* | 1.13E-263 | 1.9144458 | 0.31 | 0.027 | 5.31E-259 |
| *K07C11.7* | 1.65E-120 | 1.88810073 | 0.522 | 0.156 | 7.74E-116 |
| *col-142* | 2.24E-149 | 1.82949433 | 0.936 | 0.573 | 1.05E-144 |

### Top 20 depleted genes

| | p_val | avg_log2FC | pct.1 | pct.2 | p_val_adj |
|---|---|---|---|---|---|
| *dig-1* | 1.48E-07 | -2.1262216 | 0.353 | 0.234 | 0.00693907 |
| *F53H2.3* | 1.96E-07 | -1.2280449 | 0.115 | 0.203 | 0.00919183 |
| *mix-1* | 7.71E-10 | -1.1270753 | 0.183 | 0.296 | 3.62E-05 |
| *ipla-1* | 8.24E-08 | -0.9886066 | 0.257 | 0.352 | 0.00386426 |
| *cpr-1* | 1.49E-09 | -0.9807509 | 0.347 | 0.455 | 7.01E-05 |
| *dct-16* | 2.23E-15 | -0.9685919 | 0.715 | 0.763 | 1.05E-10 |
| *sos-1* | 4.81E-09 | -0.9529103 | 0.129 | 0.234 | 0.00022559 |
| *H06I04.3* | 3.81E-12 | -0.9406015 | 0.133 | 0.265 | 1.79E-07 |
| *npp-8* | 1.57E-09 | -0.935673 | 0.127 | 0.236 | 7.36E-05 |
| *lpd-3* | 4.23E-09 | -0.8951715 | 0.226 | 0.335 | 0.00019824 |
| *lys-2* | 5.79E-07 | -0.8929382 | 0.409 | 0.475 | 0.02716372 |
| *crh-1* | 3.12E-07 | -0.8856172 | 0.17 | 0.264 | 0.01464534 |
| *Y119D3B.21* | 2.87E-10 | -0.8745702 | 0.604 | 0.66 | 1.35E-05 |
| *sql-1* | 1.89E-08 | -0.8708044 | 0.133 | 0.232 | 0.0008851 |
| *vit-2* | 1.03E-08 | -0.8630216 | 0.678 | 0.725 | 0.00048083 |
| *mdt-26* | 6.32E-07 | -0.859684 | 0.158 | 0.244 | 0.02962771 |
| *Y79H2A.3* | 1.91E-07 | -0.8571863 | 0.144 | 0.239 | 0.00895628 |
| *asp-3* | 8.77E-09 | -0.8566837 | 0.515 | 0.594 | 0.00041158 |
| *F57F5.1* | 4.75E-08 | -0.8506876 | 0.4 | 0.502 | 0.00222874 |
| *fib-1* | 1.24E-09 | -0.8498251 | 0.185 | 0.299 | 5.84E-05 |

### Tissue enrichment

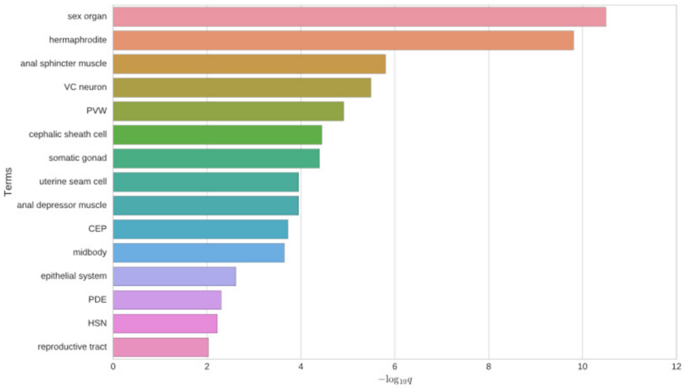


### GO enrichment

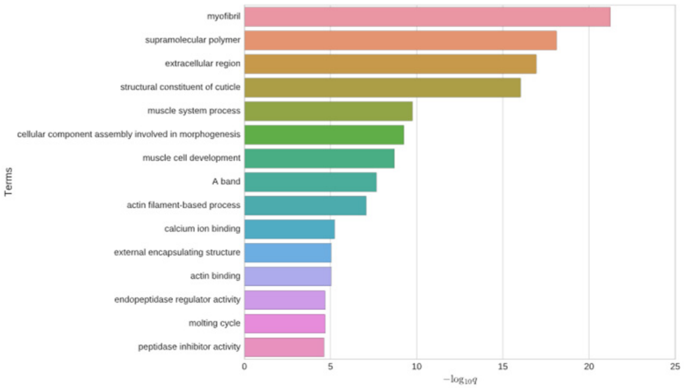


### Phenotype enrichment

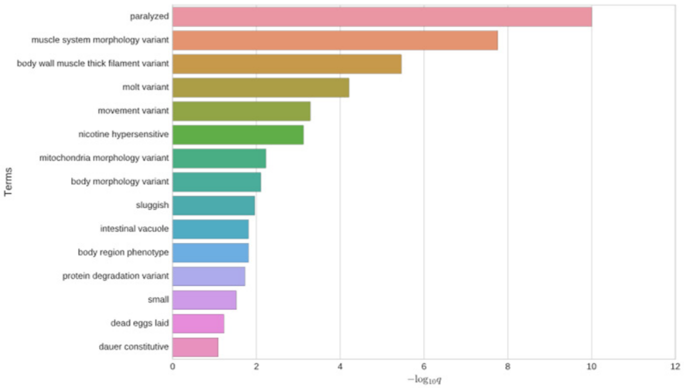


### *nas-37*

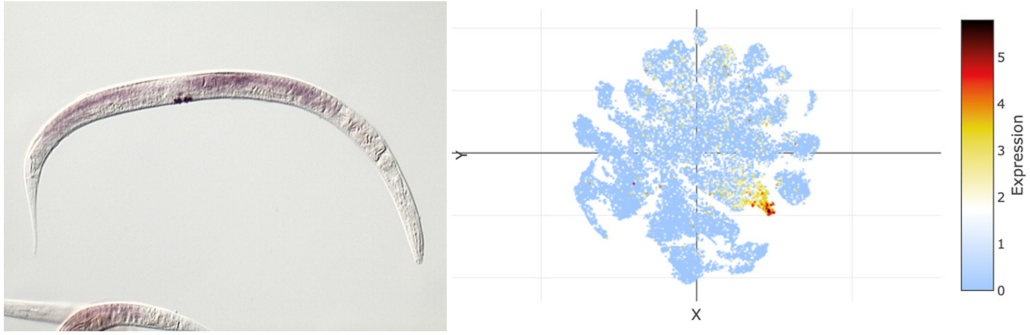


### *hsp-43*

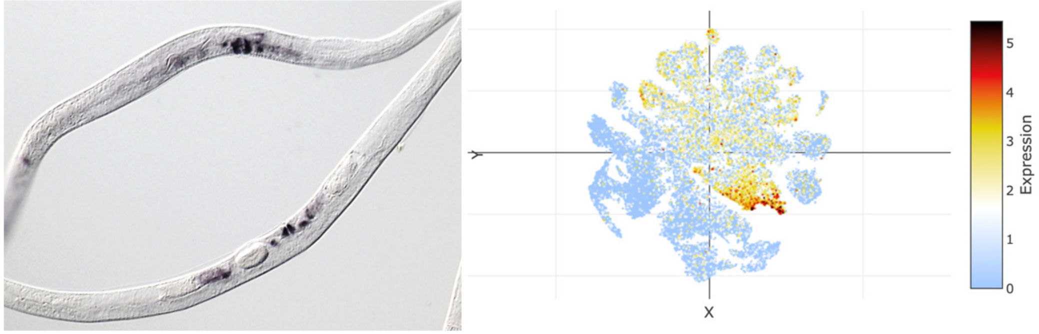

### t-SNE showing cell type 22

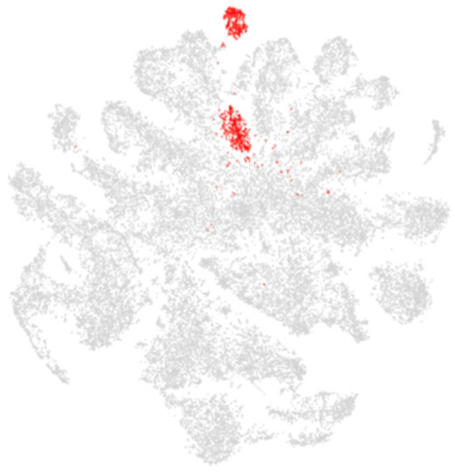

### Top 20 enriched genes

| | p_val | avg_log2FC | pct.1 | pct.2 | p_val_adj |
|---|---|---|---|---|---|
| *T28D6.3* | 0 | 3.91335063 | 0.936 | 0.22 | 0 |
| *ZK669.3* | 3.32E-285 | 2.66178642 | 0.691 | 0.136 | 1.56E-280 |
| *gst-41* | 9.84E-274 | 2.57465492 | 0.338 | 0.03 | 4.62E-269 |
| *B0238.12* | 1.14E-170 | 2.39212287 | 0.375 | 0.059 | 5.33E-166 |
| *abf-6* | 1.65E-204 | 2.30638666 | 0.471 | 0.078 | 7.72E-200 |
| *acp-6* | 2.74E-183 | 2.17380794 | 0.963 | 0.536 | 1.29E-178 |
| *ZC443.1* | 0 | 2.11871188 | 0.447 | 0.038 | 0 |
| *tni-3* | 3.53E-114 | 2.04566023 | 0.629 | 0.225 | 1.66E-109 |
| *F55H12.4* | 1.23E-97 | 2.03481537 | 0.765 | 0.39 | 5.77E-93 |
| *nlp-77* | 1.39E-190 | 2.010419 | 0.998 | 0.697 | 6.53E-186 |
| *acs-3* | 1.40E-102 | 1.97707573 | 0.18 | 0.022 | 6.59E-98 |
| *T26C5.4* | 1.74E-191 | 1.95778926 | 0.465 | 0.081 | 8.14E-187 |
| *D1086.1* | 1.70E-116 | 1.95346603 | 0.518 | 0.149 | 7.98E-112 |
| *vap-2* | 1.03E-219 | 1.93097021 | 0.522 | 0.091 | 4.85E-215 |
| *col-150* | 2.93E-167 | 1.8655416 | 0.572 | 0.136 | 1.37E-162 |
| *del-6* | 1.13E-121 | 1.84253629 | 0.735 | 0.316 | 5.30E-117 |
| *Y73F4A.2* | 1.29E-160 | 1.83960282 | 0.478 | 0.096 | 6.07E-156 |
| *tag-290* | 1.43E-147 | 1.82229644 | 0.373 | 0.065 | 6.70E-143 |
| *C45B2.1* | 7.11E-114 | 1.77727406 | 0.846 | 0.43 | 3.33E-109 |
| *Y113G7A.16* | 0 | 1.74637826 | 0.287 | 0.014 | 0 |

### Top 20 depleted genes

| | p_val | avg_log2FC | pct.1 | pct.2 | p_val_adj |
|---|---|---|---|---|---|
| *F53H2.3* | 1.51E-07 | -1.3168283 | 0.112 | 0.203 | 0.0070754 |
| *F57F5.1* | 7.75E-15 | -1.2806699 | 0.353 | 0.503 | 3.63E-10 |
| *vit-2* | 1.89E-15 | -1.2784977 | 0.673 | 0.725 | 8.86E-11 |
| *mix-1* | 1.44E-08 | -1.1876436 | 0.191 | 0.295 | 0.00067357 |
| *afd-1* | 2.74E-07 | -1.1640927 | 0.25 | 0.338 | 0.0128373 |
| *daf-2* | 7.47E-11 | -1.1464714 | 0.086 | 0.203 | 3.50E-06 |
| *elpc-2* | 1.31E-11 | -1.1291539 | 0.079 | 0.199 | 6.16E-07 |
| *vit-3* | 6.37E-13 | -1.1086278 | 0.638 | 0.714 | 2.99E-08 |
| *aakg-1* | 8.90E-08 | -1.1070908 | 0.118 | 0.21 | 0.00417221 |
| *Y79H2A.3* | 2.17E-08 | -1.0841757 | 0.14 | 0.239 | 0.00101978 |
| *F53H1.4* | 6.62E-10 | -1.080703 | 0.121 | 0.232 | 3.11E-05 |
| *vit-1* | 1.89E-10 | -1.0753422 | 0.61 | 0.677 | 8.87E-06 |
| *vit-6* | 1.89E-11 | -1.0661482 | 0.721 | 0.756 | 8.85E-07 |
| *cpr-1* | 8.06E-09 | -1.0544938 | 0.342 | 0.455 | 0.00037798 |
| *sos-1* | 5.31E-07 | -1.0541899 | 0.147 | 0.233 | 0.02492856 |
| *npp-8* | 9.75E-12 | -1.0412026 | 0.107 | 0.236 | 4.58E-07 |
| *vit-5* | 2.41E-09 | -1.0344991 | 0.741 | 0.764 | 0.00011301 |
| *lpd-3* | 1.84E-09 | -1.0333984 | 0.224 | 0.334 | 8.61E-05 |
| *C17F4.7* | 6.95E-08 | -1.0279658 | 0.702 | 0.712 | 0.00326114 |
| *cpr-6* | 8.78E-12 | -1.0100447 | 0.632 | 0.681 | 4.12E-07 |

### Tissue enrichment

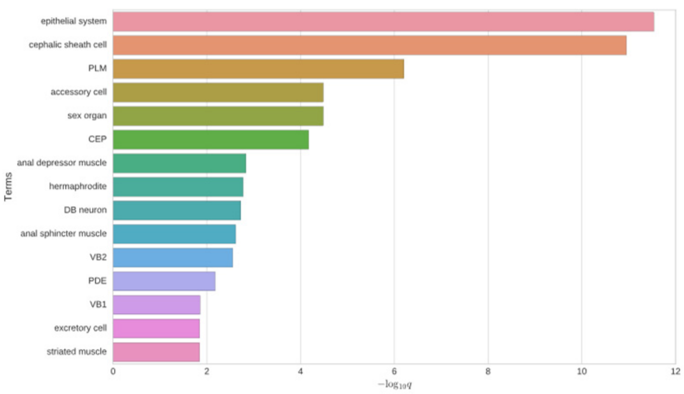


### GO enrichment

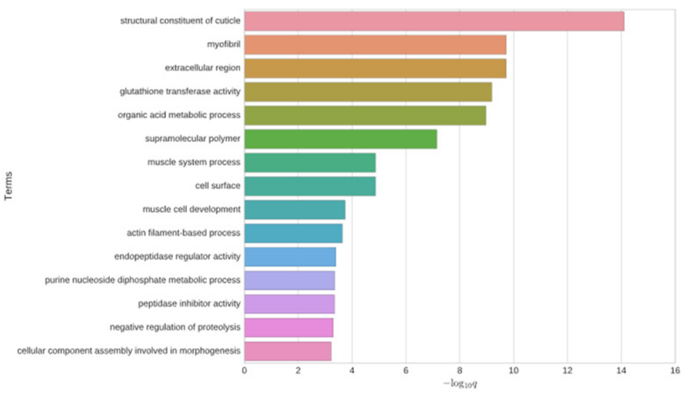


### Phenotype enrichment

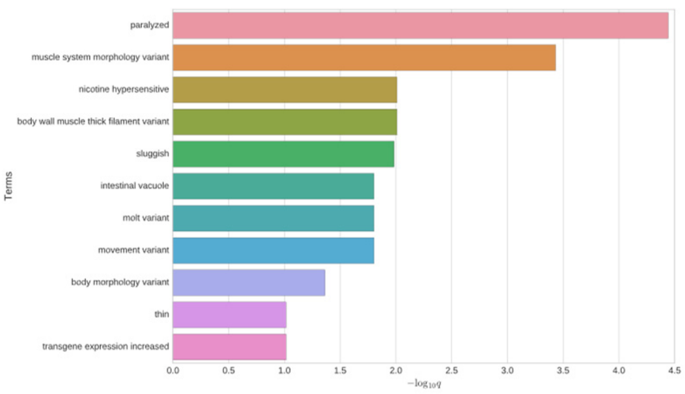


### *gst-41*

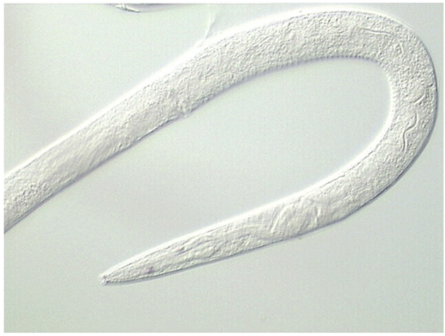

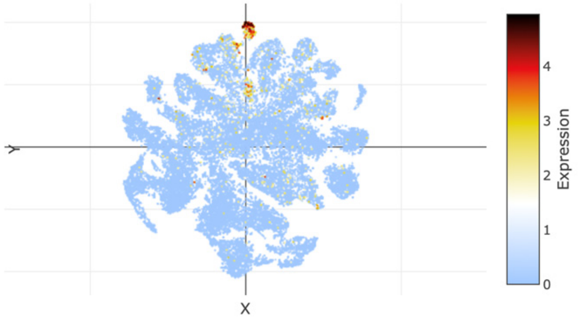


## *B0238.12*

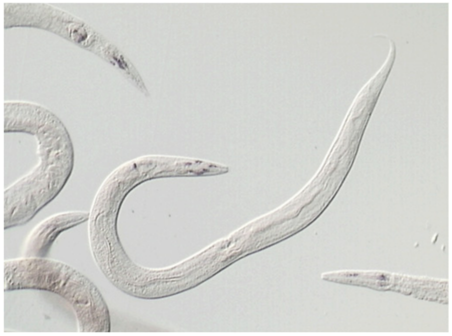

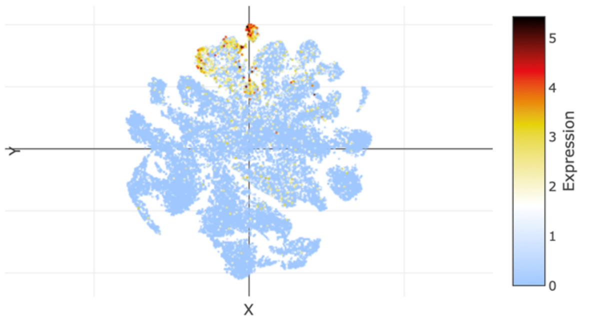

### t-SNE showing cell type 23

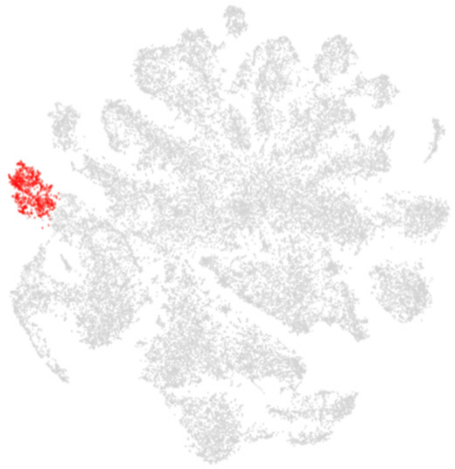

### Top 20 enriched genes

| | p_val | avg_log2FC | pct.1 | pct.2 | p_val_adj |
|---|---|---|---|---|---|
| *T21C9.13* | 0 | 5.07789238 | 0.964 | 0.132 | 0 |
| *F14H3.6* | 0 | 4.1271477 | 0.916 | 0.094 | 0 |
| *C05C10.5* | 0 | 4.11788199 | 0.918 | 0.158 | 0 |
| *pos-1* | 0 | 4.11616973 | 0.962 | 0.191 | 0 |
| *F08F3.6* | 0 | 4.06865345 | 0.89 | 0.085 | 0 |
| *spn-4* | 0 | 3.98897048 | 0.951 | 0.202 | 0 |
| *patr-1* | 0 | 3.90285022 | 0.922 | 0.183 | 0 |
| *mom-2* | 0 | 3.87843379 | 0.882 | 0.105 | 0 |
| *C37C3.9* | 0 | 3.86279559 | 0.894 | 0.114 | 0 |
| *T12G3.6* | 0 | 3.75388201 | 0.899 | 0.11 | 0 |
| *T10B11.8* | 0 | 3.71742132 | 0.753 | 0.072 | 0 |
| *mtk-1* | 0 | 3.71654975 | 0.798 | 0.151 | 0 |
| *tmem-131* | 0 | 3.7144092 | 0.869 | 0.197 | 0 |
| *rbr-2* | 0 | 3.70569234 | 0.816 | 0.185 | 0 |
| *clec-87* | 0 | 3.68817575 | 0.977 | 0.351 | 0 |
| *Y4C6A.3* | 0 | 3.6619247 | 0.802 | 0.056 | 0 |
| *neg-1* | 0 | 3.62277969 | 0.781 | 0.082 | 0 |
| *era-1* | 0 | 3.60817646 | 0.882 | 0.13 | 0 |
| *C13F10.7* | 0 | 3.59369398 | 0.791 | 0.081 | 0 |
| *zif-1* | 0 | 3.58539126 | 0.857 | 0.084 | 0 |

### Top 20 depleted genes

| | p_val | avg_log2FC | pct.1 | pct.2 | p_val_adj |
|---|---|---|---|---|---|
| *ule-4* | 1.64E-92 | -5.2174366 | 0.038 | 0.519 | 7.67E-88 |
| *far-2* | 2.67E-191 | -4.9787011 | 0.08 | 0.784 | 1.25E-186 |
| *col-122* | 1.08E-182 | -4.8207586 | 0.051 | 0.756 | 5.06E-178 |
| *col-119* | 9.43E-178 | -4.72203 | 0.053 | 0.747 | 4.42E-173 |
| *col-101* | 1.96E-170 | -4.6967788 | 0.067 | 0.733 | 9.18E-166 |
| *act-4* | 3.07E-175 | -4.6217209 | 0.048 | 0.736 | 1.44E-170 |
| *col-140* | 7.43E-180 | -4.6023819 | 0.059 | 0.755 | 3.49E-175 |
| *col-181* | 4.42E-176 | -4.5743292 | 0.049 | 0.742 | 2.08E-171 |
| *col-184* | 6.51E-169 | -4.4634474 | 0.04 | 0.719 | 3.05E-164 |
| *unc-54* | 8.33E-157 | -4.46105 | 0.051 | 0.699 | 3.91E-152 |
| *col-124* | 3.38E-166 | -4.3704093 | 0.048 | 0.723 | 1.59E-161 |
| *col-20* | 5.79E-171 | -4.332734 | 0.059 | 0.739 | 2.72E-166 |
| *lev-11* | 2.82E-178 | -4.3296425 | 0.042 | 0.744 | 1.32E-173 |
| *D1086.10* | 3.64E-63 | -4.3175522 | 0.03 | 0.402 | 1.71E-58 |
| *col-95* | 9.42E-119 | -4.2974024 | 0.051 | 0.602 | 4.42E-114 |
| *pat-10* | 1.09E-167 | -4.2782809 | 0.038 | 0.724 | 5.09E-163 |
| *col-80* | 3.57E-175 | -4.2730347 | 0.07 | 0.751 | 1.67E-170 |
| *nlp-77* | 4.94E-162 | -4.2670336 | 0.072 | 0.72 | 2.32E-157 |
| *cpn-3* | 1.19E-154 | -4.2597754 | 0.061 | 0.697 | 5.56E-150 |
| *col-143* | 2.31E-180 | -4.2557594 | 0.078 | 0.764 | 1.08E-175 |

### Tissue enrichment

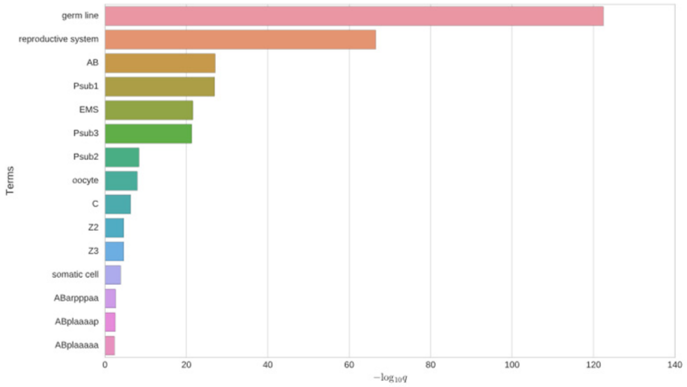


### GO enrichment

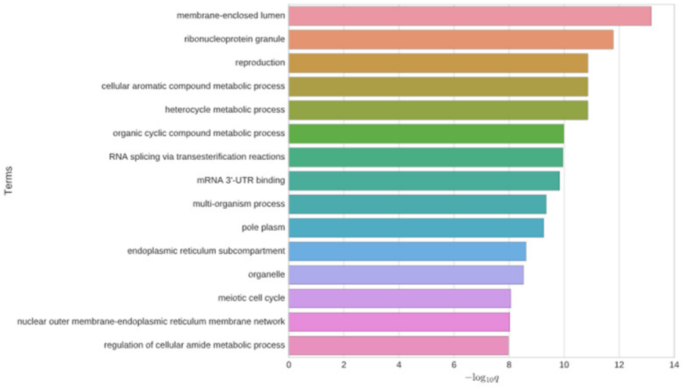


### Phenotype enrichment

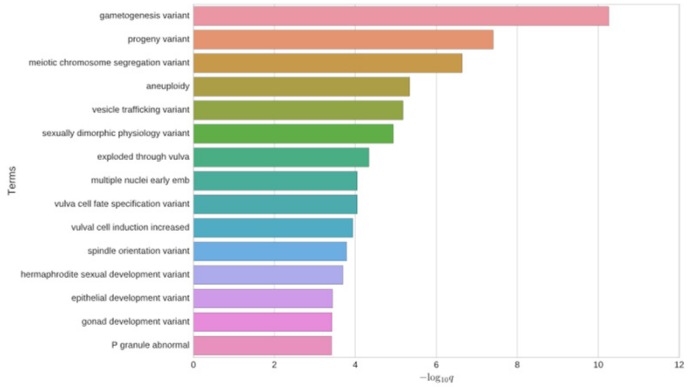


### *glp-1*

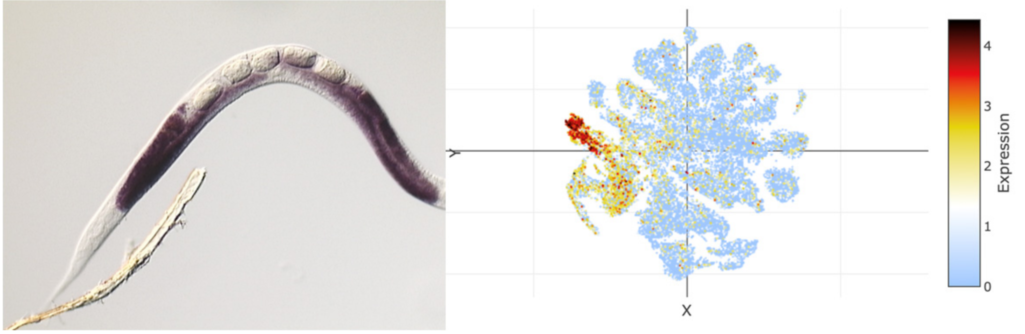


### *gld-1*

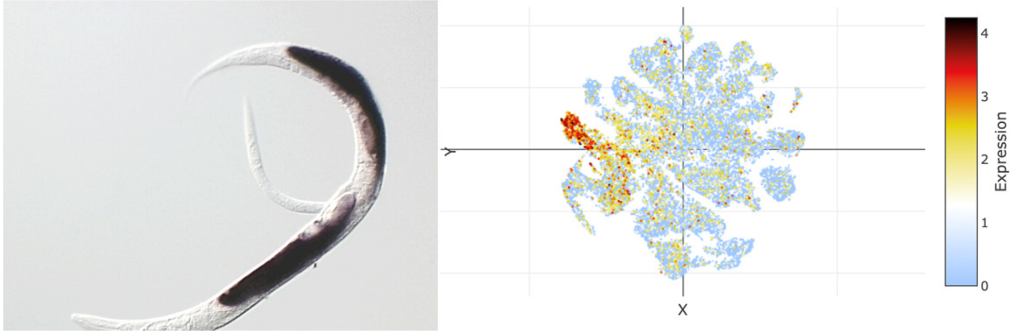

### t-SNE showing cell type 24

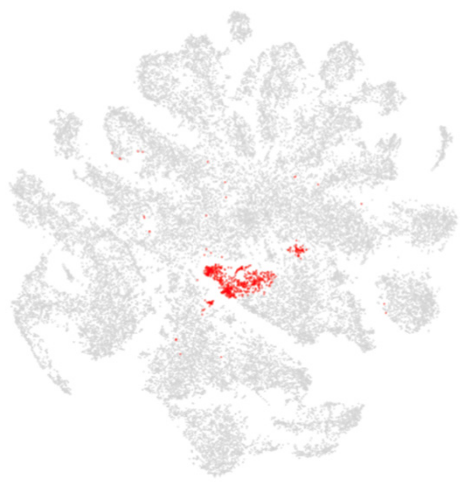

### Top 20 enriched genes

| | p_val | avg_log2FC | pct.1 | pct.2 | p_val_adj |
|---|---|---|---|---|---|
| *ule-1* | 0 | 5.06501145 | 0.874 | 0.25 | 0 |
| *ule-2* | 0 | 4.40636231 | 0.914 | 0.28 | 0 |
| *D1086.7* | 0 | 4.32192192 | 0.899 | 0.219 | 0 |
| *C08A9.10* | 0 | 3.75268209 | 0.804 | 0.136 | 0 |
| *K11D12.13* | 1.40E-209 | 3.5461727 | 0.626 | 0.143 | 6.54E-205 |
| *F09E10.1* | 0 | 3.50895164 | 0.599 | 0.081 | 0 |
| *Y48G1C.13* | 0 | 3.48193198 | 0.632 | 0.083 | 0 |
| *C46C2.5* | 1.08E-298 | 3.40809568 | 0.599 | 0.094 | 5.06E-294 |
| *F59A6.12* | 0 | 3.3948197 | 0.727 | 0.084 | 0 |
| *ule-4* | 6.68E-231 | 3.34343654 | 0.954 | 0.497 | 3.13E-226 |
| *C10G8.4* | 2.87E-122 | 3.32627956 | 0.744 | 0.334 | 1.35E-117 |
| *C30G7.4* | 0 | 3.27786374 | 0.698 | 0.071 | 0 |
| *Y106G6D.8* | 1.38E-238 | 3.08765547 | 0.762 | 0.208 | 6.48E-234 |
| *ttr-34* | 1.97E-295 | 2.92212669 | 0.705 | 0.135 | 9.24E-291 |
| *hex-1* | 4.38E-128 | 2.78020561 | 0.553 | 0.166 | 2.06E-123 |
| *Y51H7C.1* | 3.65E-251 | 2.67086809 | 0.518 | 0.08 | 1.71E-246 |
| *K10C2.8* | 7.48E-99 | 2.56086893 | 0.454 | 0.134 | 3.51E-94 |
| *F08F1.4* | 6.81E-164 | 2.55133084 | 0.612 | 0.173 | 3.19E-159 |
| *T25D3.3* | 0 | 2.54687445 | 0.474 | 0.037 | 0 |
| *ttr-2* | 3.28E-109 | 2.53731675 | 0.826 | 0.455 | 1.54E-104 |

### Top 20 depleted genes

| | p_val | avg_log2FC | pct.1 | pct.2 | p_val_adj |
|---|---|---|---|---|---|
| *crh-1* | 2.35E-09 | -1.152779 | 0.145 | 0.264 | 0.00011017 |
| *spsb-1* | 6.99E-08 | -1.1264596 | 0.181 | 0.291 | 0.0032786 |
| *F53E4.1* | 3.91E-09 | -1.0097246 | 0.161 | 0.28 | 0.0001832 |
| *daf-2* | 6.08E-09 | -0.947104 | 0.095 | 0.203 | 0.00028526 |
| *elpc-2* | 3.17E-08 | -0.9275282 | 0.099 | 0.199 | 0.00148886 |
| *lmd-3* | 2.44E-09 | -0.9272528 | 0.297 | 0.41 | 0.00011433 |
| *afd-1* | 6.38E-07 | -0.9037204 | 0.24 | 0.339 | 0.0299427 |
| *H06I04.3* | 2.63E-07 | -0.8065096 | 0.167 | 0.264 | 0.01235367 |
| *tni-4* | 3.82E-07 | -0.7926128 | 0.093 | 0.184 | 0.01790638 |
| *C35B1.2* | 8.25E-09 | -0.7859282 | 0.101 | 0.209 | 0.00038703 |
| *ceh-100* | 3.09E-07 | -0.7592855 | 0.145 | 0.242 | 0.01447461 |
| *rsks-1* | 1.47E-07 | -0.7558131 | 0.137 | 0.237 | 0.00688257 |
| *unc-40* | 3.42E-08 | -0.7512286 | 0.095 | 0.196 | 0.00160331 |
| *sql-1* | 8.52E-09 | -0.7492299 | 0.121 | 0.232 | 0.00039943 |
| *Y105C5A.15* | 3.70E-08 | -0.7432723 | 0.048 | 0.137 | 0.00173431 |
| *F58B3.6* | 1.48E-11 | -0.742568 | 0.11 | 0.24 | 6.92E-07 |
| *Y48G8AL.5* | 1.85E-07 | -0.738407 | 0.068 | 0.155 | 0.00869343 |
| *egl-4* | 8.59E-11 | -0.7326871 | 0.35 | 0.482 | 4.03E-06 |
| *F16A11.1* | 2.39E-07 | -0.730184 | 0.148 | 0.243 | 0.01119518 |
| *W07E11.1* | 1.70E-07 | -0.7196724 | 0.205 | 0.304 | 0.00798622 |

### Tissue enrichment

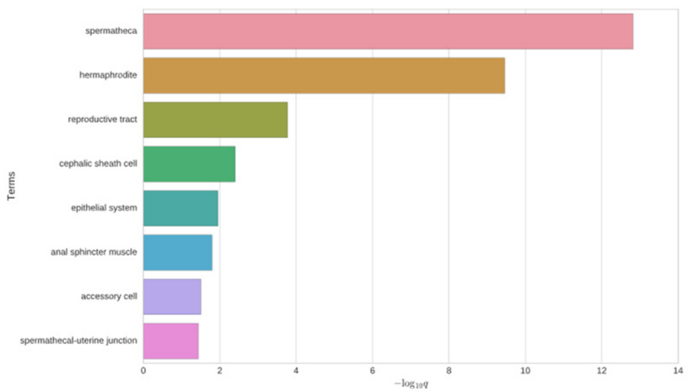


### GO enrichment

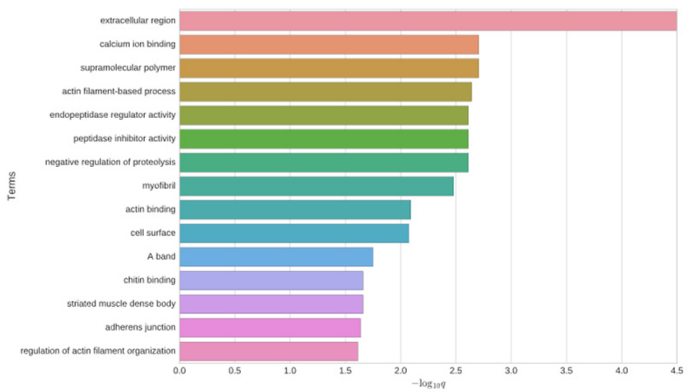


### Phenotype enrichment

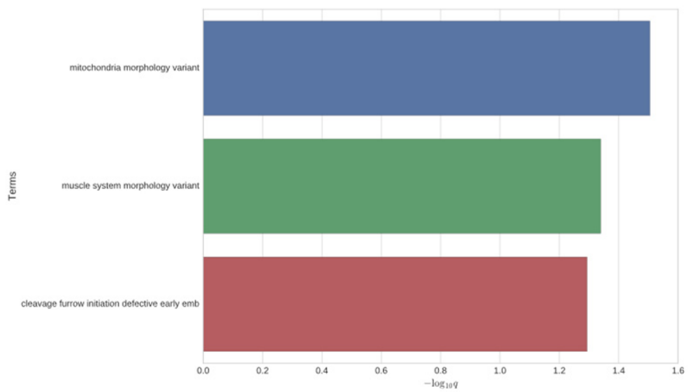


#### *pes-23*

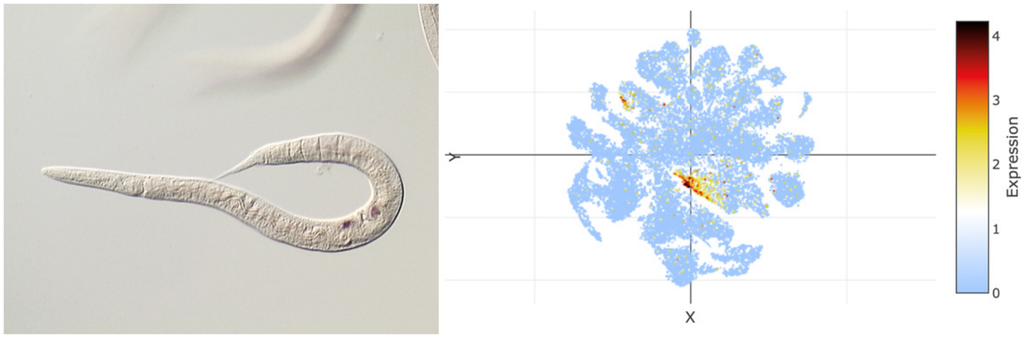


#### *ule-1*

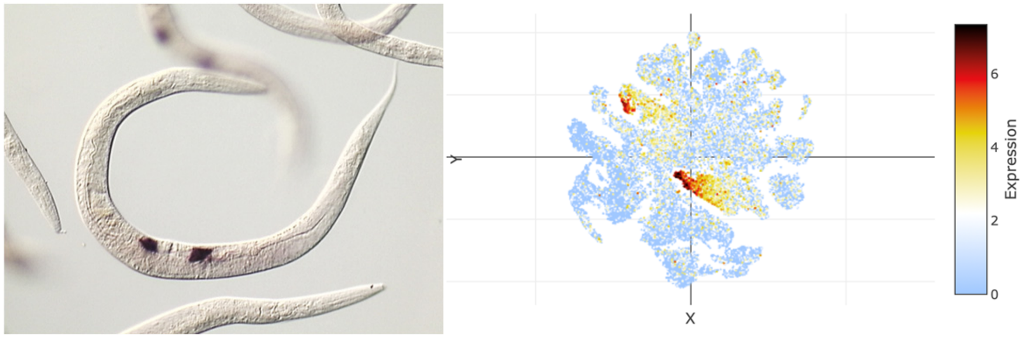

### t-SNE showing cell type 25

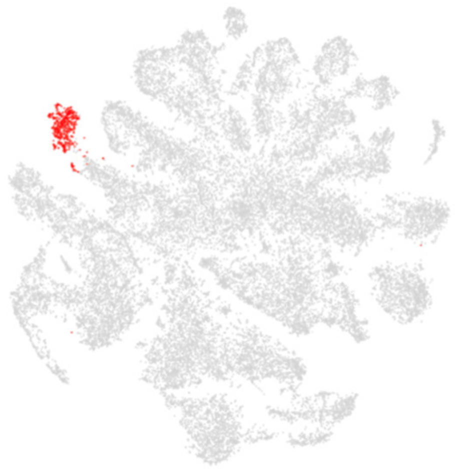

### Top 20 enriched genes

| | p_val | avg_log2FC | pct.1 | pct.2 | p_val_adj |
|---|---|---|---|---|---|
| *fasn-1* | 0 | 5.84270177 | 0.982 | 0.265 | 0 |
| *C39D10.7* | 8.09E-300 | 5.09991577 | 0.985 | 0.272 | 3.79E-295 |
| *ZC513.7* | 9.50E-223 | 4.72135562 | 0.982 | 0.42 | 4.46E-218 |
| *B0545.4* | 0 | 4.67893158 | 0.772 | 0.041 | 0 |
| *irk-1* | 0 | 4.18698545 | 0.723 | 0.014 | 0 |
| *abts-4* | 0 | 4.12604164 | 0.754 | 0.07 | 0 |
| *tbh-1* | 3.29E-273 | 3.99128796 | 0.888 | 0.207 | 1.54E-268 |
| *aak-1* | 0 | 3.86598673 | 0.86 | 0.155 | 0 |
| *skpo-1* | 4.83E-240 | 3.75765996 | 0.86 | 0.213 | 2.27E-235 |
| *frm-7* | 1.24E-174 | 3.75278549 | 0.936 | 0.436 | 5.82E-170 |
| *F45B8.5* | 0 | 3.54379944 | 0.663 | 0.015 | 0 |
| *Y105C5A.25* | 0 | 3.36408584 | 0.672 | 0.033 | 0 |
| *C44B7.5* | 1.76E-66 | 3.36197696 | 0.641 | 0.302 | 8.27E-62 |
| *W04A4.2* | 1.24E-293 | 3.33351245 | 0.535 | 0.056 | 5.82E-289 |
| *Y38E10A.14* | 1.04E-106 | 3.29254218 | 0.663 | 0.236 | 4.88E-102 |
| *W09C3.7* | 0 | 3.24707631 | 0.787 | 0.108 | 0 |
| *lim-7* | 0 | 3.22596324 | 0.578 | 0.024 | 0 |
| *C04G2.14* | 0 | 3.18178109 | 0.48 | 0.04 | 0 |
| *perm-4* | 7.25E-159 | 3.14982192 | 0.994 | 0.572 | 3.40E-154 |
| *anc-1* | 5.69E-170 | 3.08059504 | 0.994 | 0.685 | 2.67E-165 |

### Top 20 depleted genes

| | p_val | avg_log2FC | pct.1 | pct.2 | p_val_adj |
|---|---|---|---|---|---|
| *ule-4* | 5.03E-14 | -2.8176448 | 0.347 | 0.509 | 2.36E-09 |
| *D1086.10* | 4.79E-12 | -2.7534585 | 0.231 | 0.395 | 2.25E-07 |
| *nlp-77* | 1.86E-57 | -2.4992579 | 0.325 | 0.709 | 8.72E-53 |
| *lbp-2* | 1.99E-42 | -2.4948408 | 0.322 | 0.636 | 9.34E-38 |
| *ttr-16* | 3.24E-40 | -2.4426892 | 0.368 | 0.648 | 1.52E-35 |
| *ule-2* | 9.60E-08 | -2.359075 | 0.176 | 0.295 | 0.00450282 |
| *cpn-3* | 6.90E-41 | -2.2887364 | 0.419 | 0.685 | 3.24E-36 |
| *col-95* | 4.48E-30 | -2.2716506 | 0.298 | 0.593 | 2.10E-25 |
| *col-140* | 1.21E-54 | -2.2649531 | 0.419 | 0.742 | 5.65E-50 |
| *C35B1.4* | 1.44E-12 | -2.2265026 | 0.097 | 0.263 | 6.78E-08 |
| *C10G8.4* | 1.27E-07 | -2.2191065 | 0.231 | 0.345 | 0.00597648 |
| *col-119* | 1.91E-49 | -2.217853 | 0.438 | 0.734 | 8.97E-45 |
| *col-122* | 8.48E-52 | -2.1939011 | 0.426 | 0.743 | 3.98E-47 |
| *col-184* | 1.32E-46 | -2.1271967 | 0.413 | 0.706 | 6.20E-42 |
| *col-20* | 1.30E-49 | -2.1167704 | 0.377 | 0.728 | 6.12E-45 |
| *col-181* | 2.34E-46 | -2.1116332 | 0.453 | 0.729 | 1.10E-41 |
| *tnt-2* | 1.92E-36 | -2.105879 | 0.31 | 0.606 | 9.01E-32 |
| *col-101* | 1.10E-43 | -2.0968267 | 0.429 | 0.721 | 5.16E-39 |
| *fipr-2* | 2.66E-12 | -2.0924108 | 0.046 | 0.197 | 1.25E-07 |
| *col-80* | 1.81E-50 | -2.079209 | 0.416 | 0.739 | 8.48E-46 |

### Tissue enrichment

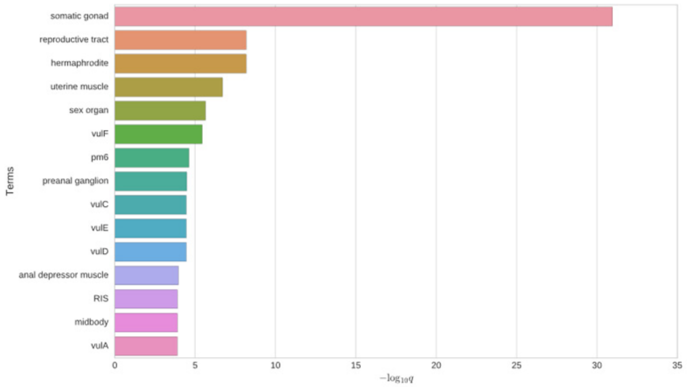


### GO enrichment

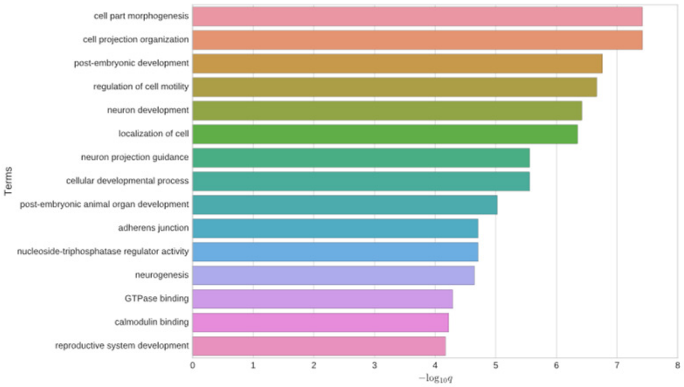


### Phenotype enrichment

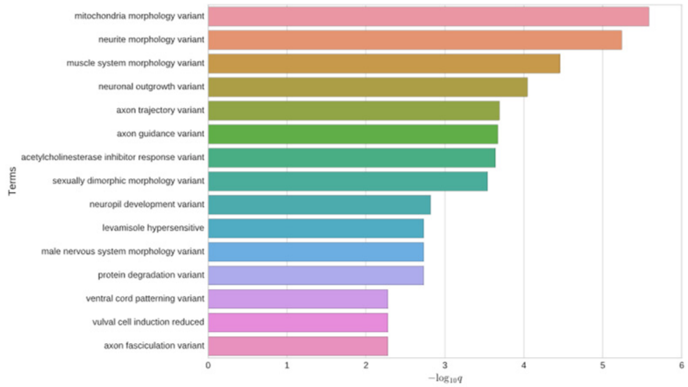


## *C39D10.7*

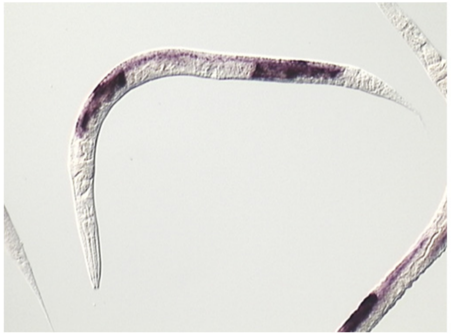

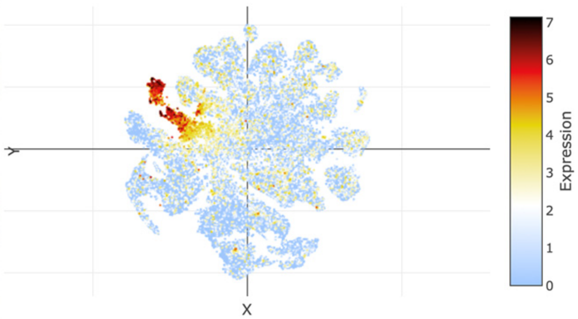


### *fasn-1*

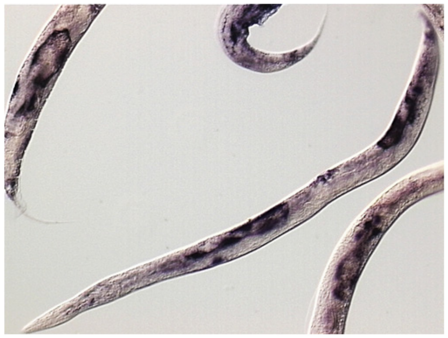

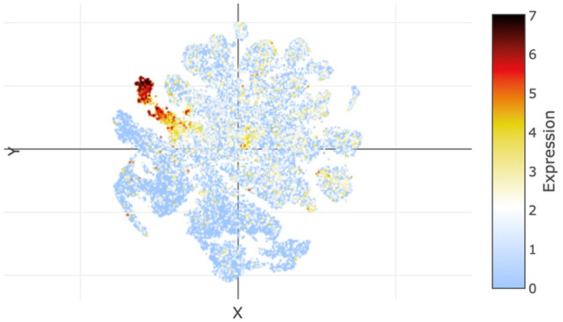

### t-SNE showing cell type 26

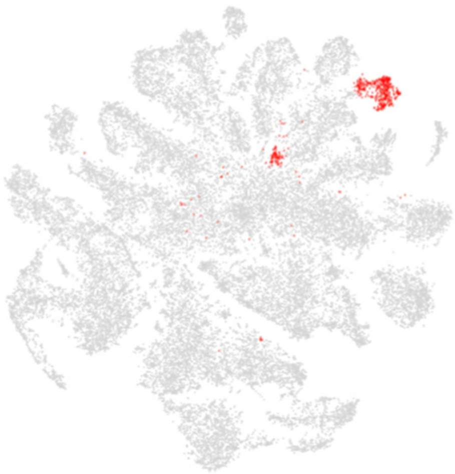

### Top 20 enriched genes

| | p_val | avg_log2FC | pct.1 | pct.2 | p_val_adj |
|---|---|---|---|---|---|
| *nspc-4* | 0 | 4.83083585 | 0.897 | 0.111 | 0 |
| *nspc-7* | 0 | 4.48240695 | 0.836 | 0.086 | 0 |
| *nspc-14* | 0 | 4.31236431 | 0.805 | 0.071 | 0 |
| *nspc-20* | 0 | 4.21284414 | 0.775 | 0.07 | 0 |
| *nspc-9* | 0 | 3.91972667 | 0.763 | 0.059 | 0 |
| *nspc-10* | 0 | 3.5758342 | 0.69 | 0.047 | 0 |
| *nspc-1* | 0 | 3.55081748 | 0.684 | 0.038 | 0 |
| *nspc-13* | 0 | 3.29167112 | 0.626 | 0.03 | 0 |
| *nspc-3* | 0 | 2.23152161 | 0.45 | 0.019 | 0 |
| *flp-1* | 1.87E-52 | 2.13844162 | 0.225 | 0.046 | 8.77E-48 |
| *nlp-49* | 7.24E-81 | 1.88416721 | 0.286 | 0.05 | 3.40E-76 |
| *lips-15* | 6.71E-65 | 1.69913496 | 0.465 | 0.147 | 3.15E-60 |
| *nspc-5* | 0 | 1.55568815 | 0.249 | 0.008 | 0 |
| *D1086.3* | 2.43E-50 | 1.47985337 | 0.696 | 0.348 | 1.14E-45 |
| *flp-11* | 3.56E-26 | 1.43407009 | 0.188 | 0.055 | 1.67E-21 |
| *F56D6.8* | 3.06E-28 | 1.41220853 | 0.441 | 0.209 | 1.44E-23 |
| *F36D1.7* | 1.39E-139 | 1.38164486 | 0.292 | 0.033 | 6.54E-135 |
| *hmit-1.2* | 4.51E-76 | 1.34085027 | 0.207 | 0.029 | 2.11E-71 |
| *F56D6.9* | 1.73E-22 | 1.23794153 | 0.474 | 0.254 | 8.13E-18 |
| *dct-8* | 1.15E-32 | 1.19908386 | 0.407 | 0.168 | 5.41E-28 |

### Top 20 depleted genes

| | p_val | avg_log2FC | pct.1 | pct.2 | p_val_adj |
|---|---|---|---|---|---|
| *H06I04.3* | 6.44E-08 | -1.0927223 | 0.149 | 0.263 | 0.00301887 |

### Tissue enrichment

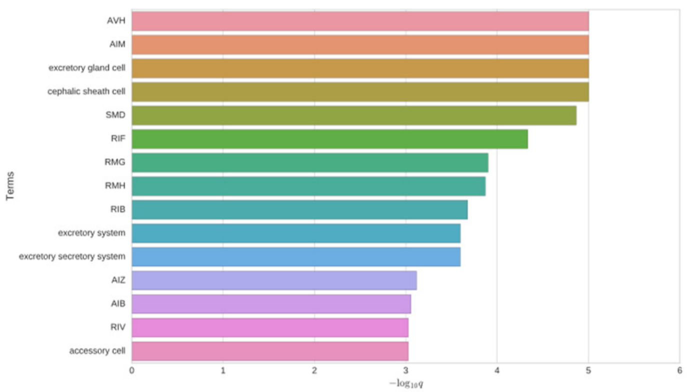


### GO enrichment

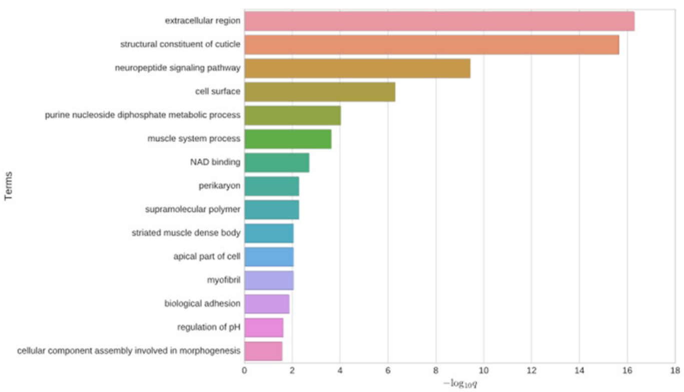


### Phenotype enrichment

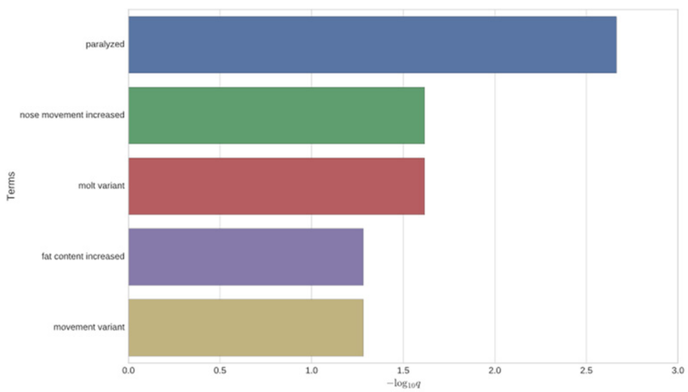


### *nspc-7*

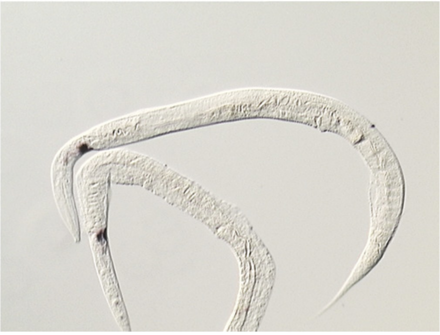

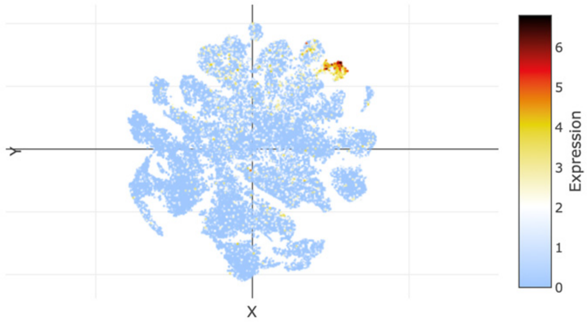


### *nspc-9*

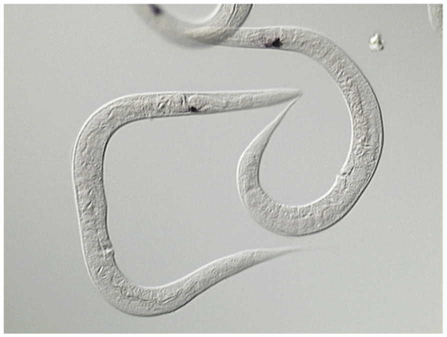

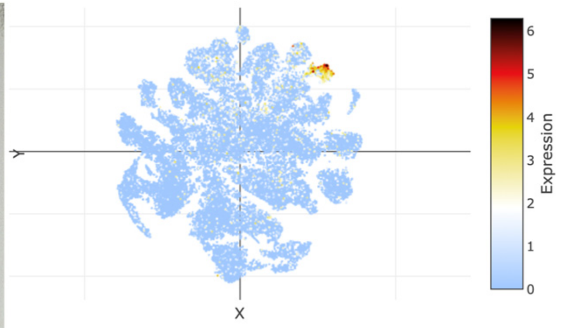

### t-SNE showing cell type 27

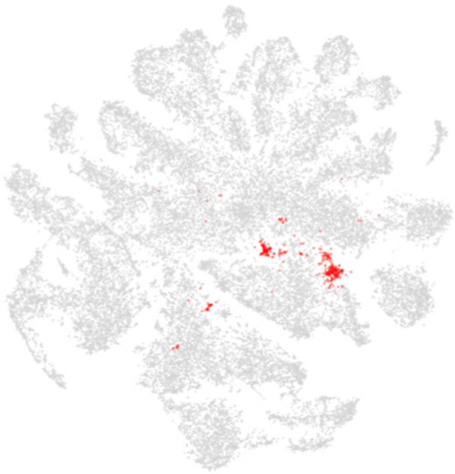

### Top 20 enriched genes

| | p_val | avg_log2FC | pct.1 | pct.2 | p_val_adj |
|---|---|---|---|---|---|
| *pks-1* | 8.89E-89 | 4.0228231 | 0.342 | 0.052 | 4.17E-84 |
| *acbp-6* | 1.03E-86 | 3.74026067 | 0.406 | 0.077 | 4.84E-82 |
| *F22F4.5* | 2.99E-167 | 3.71888228 | 0.41 | 0.042 | 1.40E-162 |
| *C53B7.2* | 2.65E-102 | 3.71641946 | 0.603 | 0.146 | 1.24E-97 |
| *cht-3* | 3.36E-67 | 3.25922289 | 0.761 | 0.344 | 1.58E-62 |
| *Y71H2B.1* | 1.10E-151 | 3.07514599 | 0.397 | 0.043 | 5.15E-147 |
| *C53B7.3* | 4.34E-58 | 2.84432466 | 0.645 | 0.248 | 2.03E-53 |
| *lbp-3* | 1.76E-60 | 2.68080243 | 0.679 | 0.288 | 8.24E-56 |
| *K10C2.12* | 2.94E-107 | 2.37296349 | 0.303 | 0.034 | 1.38E-102 |
| *bgal-1* | 6.02E-61 | 2.32577319 | 0.509 | 0.152 | 2.82E-56 |
| *pod-2* | 5.45E-11 | 2.31473403 | 0.679 | 0.547 | 2.56E-06 |
| *C09D4.2* | 1.72E-71 | 2.22813074 | 0.59 | 0.182 | 8.08E-67 |
| *hsp-12.3* | 2.43E-46 | 2.19630957 | 0.423 | 0.127 | 1.14E-41 |
| *mig-6* | 5.25E-66 | 2.11583586 | 0.786 | 0.362 | 2.46E-61 |
| *ost-1* | 6.55E-40 | 1.96370231 | 0.863 | 0.616 | 3.07E-35 |
| *lbp-1* | 5.88E-38 | 1.94751109 | 0.62 | 0.293 | 2.76E-33 |
| *ZC412.3* | 4.07E-41 | 1.90713406 | 0.521 | 0.203 | 1.91E-36 |
| *let-2* | 3.73E-41 | 1.90557376 | 0.838 | 0.572 | 1.75E-36 |
| *timp-1* | 7.54E-56 | 1.85244889 | 0.5 | 0.152 | 3.54E-51 |
| *acs-1* | 6.59E-19 | 1.78451102 | 0.632 | 0.404 | 3.09E-14 |

### Top 20 depleted genes

| | p_val | avg_log2FC | pct.1 | pct.2 | p_val_adj |
|---|---|---|---|---|---|
| *dig-1* | 1.12E-13 | -1.268791 | 0.436 | 0.235 | 5.26E-09 |
| *ule-4* | 1.93E-16 | -0.3599361 | 0.752 | 0.504 | 9.04E-12 |
| *ZK813.2* | 8.17E-07 | -0.3408854 | 0.449 | 0.288 | 0.03832419 |
| *Y22D7AR.10* | 2.80E-07 | -0.3006009 | 0.457 | 0.291 | 0.01313691 |

### Tissue enrichment

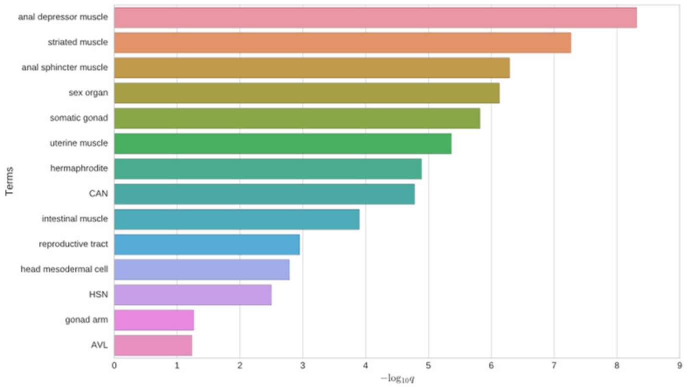


### GO enrichment

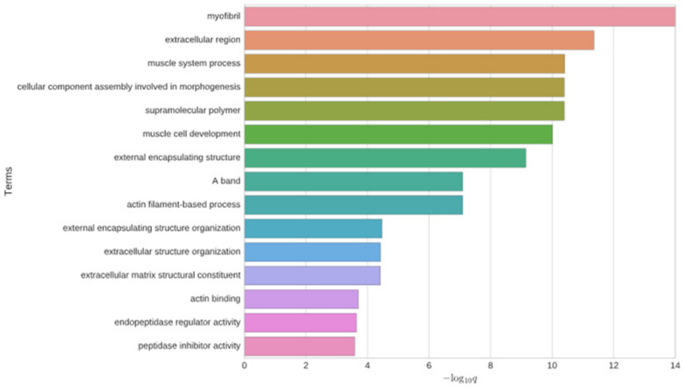


### Phenotype enrichment

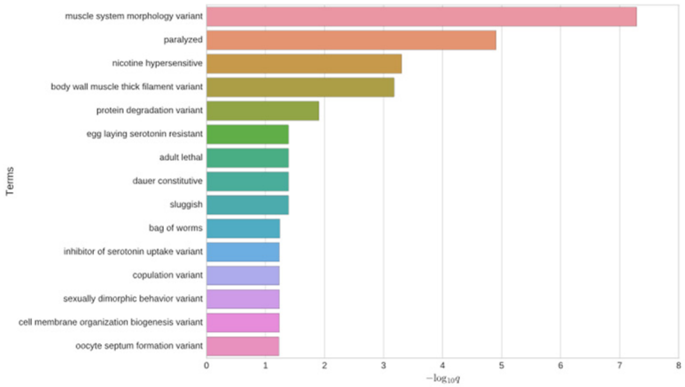


## *Y59C2A.1*

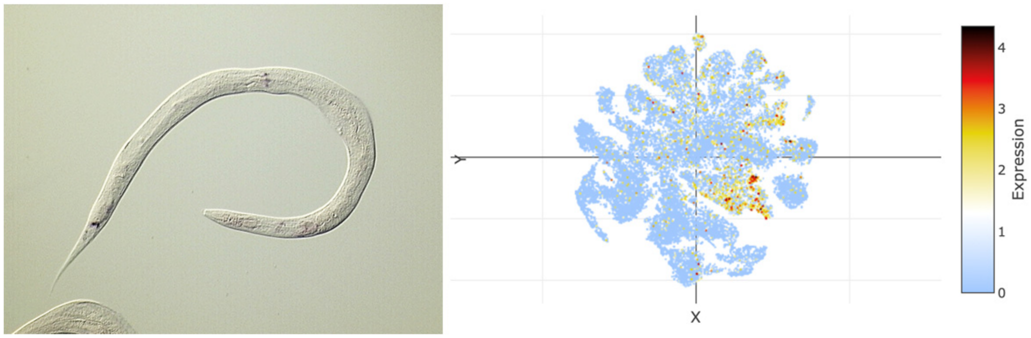


## *F22E12.1*

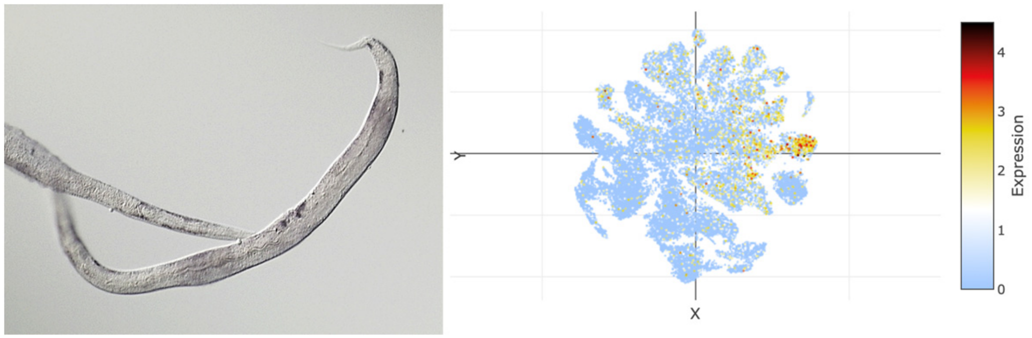

### t-SNE showing cell type 28

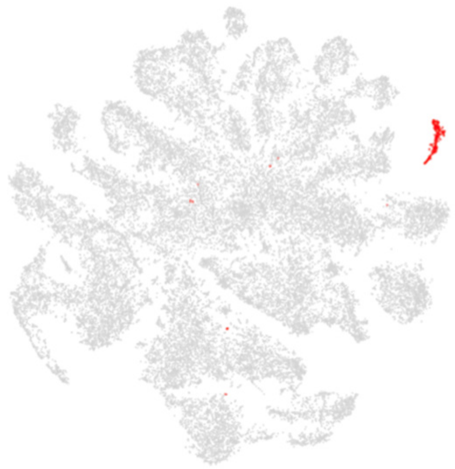

### Top 20 enriched genes

| | p_val | avg_log2FC | pct.1 | pct.2 | p_val_adj |
|---|---|---|---|---|---|
| *dig-1* | 3.23E-202 | 8.57695228 | 0.959 | 0.23 | 1.51E-197 |
| *aman-1* | 0 | 7.8349216 | 0.933 | 0.031 | 0 |
| *ZC116.3* | 0 | 7.38400593 | 0.846 | 0.012 | 0 |
| *inos-1* | 8.18E-294 | 6.72525712 | 0.903 | 0.121 | 3.84E-289 |
| *T19C3.5* | 0 | 6.33563494 | 0.81 | 0.016 | 0 |
| *Y75B7AL.2* | 0 | 6.10029465 | 0.738 | 0.011 | 0 |
| *Y73F4A.1* | 0 | 5.89322697 | 0.918 | 0.025 | 0 |
| *mig-6* | 1.45E-149 | 5.81111125 | 0.974 | 0.361 | 6.78E-145 |
| *ttr-1* | 0 | 5.67612335 | 0.933 | 0.04 | 0 |
| *cup-4* | 0 | 5.49147265 | 0.867 | 0.021 | 0 |
| *clec-145* | 0 | 5.39558576 | 0.795 | 0.011 | 0 |
| *far-1* | 2.08E-126 | 5.11466402 | 0.995 | 0.628 | 9.77E-122 |
| *Y116A8C.3* | 0 | 4.90522801 | 0.882 | 0.022 | 0 |
| *lgc-26* | 0 | 4.87721062 | 0.795 | 0.008 | 0 |
| *clec-178* | 0 | 4.75455192 | 0.867 | 0.015 | 0 |
| *ZC513.7* | 7.55E-129 | 4.58470428 | 0.974 | 0.424 | 3.54E-124 |
| *F35C12.3* | 0 | 4.47929833 | 0.882 | 0.027 | 0 |
| *unc-122* | 0 | 4.32605415 | 0.723 | 0.008 | 0 |
| *wrk-1* | 5.55E-271 | 4.30351118 | 0.851 | 0.112 | 2.60E-266 |
| *lgc-25* | 0 | 4.15864043 | 0.687 | 0.004 | 0 |

### Top 20 depleted genes

| | p_val | avg_log2FC | pct.1 | pct.2 | p_val_adj |
|---|---|---|---|---|---|
| *ule-4* | 2.25E-10 | -2.6926624 | 0.297 | 0.509 | 1.06E-05 |
| *D1086.10* | 1.55E-08 | -2.507027 | 0.205 | 0.394 | 0.00072772 |
| *ttn-1* | 9.59E-11 | -2.1883218 | 0.251 | 0.471 | 4.50E-06 |
| *vit-4* | 1.01E-18 | -1.5936053 | 0.379 | 0.684 | 4.75E-14 |
| *F57F5.1* | 2.85E-11 | -1.5338714 | 0.297 | 0.501 | 1.34E-06 |
| *dod-6* | 3.10E-08 | -1.5220349 | 0.108 | 0.287 | 0.00145238 |
| *perm-4* | 1.21E-10 | -1.5159118 | 0.359 | 0.581 | 5.66E-06 |
| *vit-3* | 1.40E-22 | -1.5103616 | 0.374 | 0.715 | 6.58E-18 |
| *ttr-51* | 2.49E-20 | -1.5058136 | 0.385 | 0.683 | 1.17E-15 |
| *C23H5.8* | 1.15E-11 | -1.5027429 | 0.226 | 0.454 | 5.41E-07 |
| *vit-1* | 1.11E-14 | -1.4145753 | 0.431 | 0.677 | 5.23E-10 |
| *lys-1* | 5.25E-15 | -1.4116604 | 0.308 | 0.573 | 2.46E-10 |
| *unc-54* | 2.97E-16 | -1.3933393 | 0.395 | 0.686 | 1.39E-11 |
| *act-3* | 2.48E-21 | -1.3773647 | 0.467 | 0.776 | 1.16E-16 |
| *mlc-3* | 8.12E-19 | -1.3579575 | 0.385 | 0.685 | 3.81E-14 |
| *vit-5* | 1.04E-19 | -1.3515589 | 0.482 | 0.766 | 4.86E-15 |
| *ketn-1* | 2.78E-11 | -1.343711 | 0.19 | 0.411 | 1.30E-06 |
| *ftn-2* | 9.77E-18 | -1.3416391 | 0.318 | 0.602 | 4.58E-13 |
| *C17F4.7* | 3.81E-17 | -1.3296595 | 0.426 | 0.714 | 1.79E-12 |
| *mup-2* | 2.50E-11 | -1.3293507 | 0.364 | 0.562 | 1.17E-06 |

### Tissue enrichment

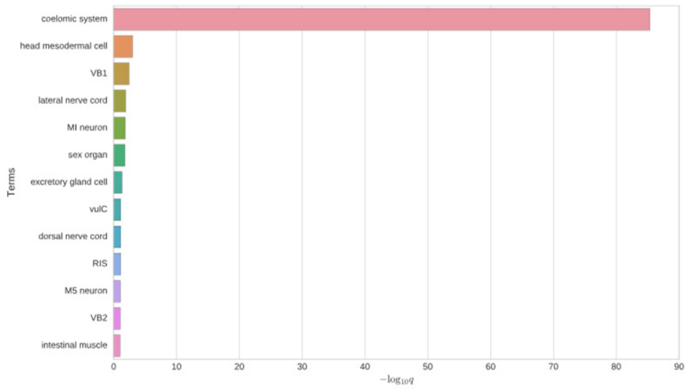


### GO enrichment

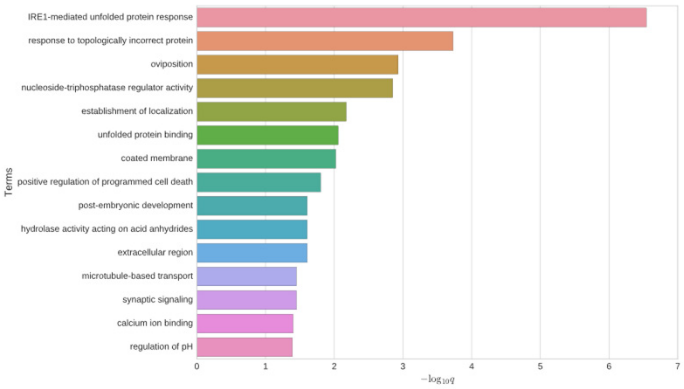


### Phenotype enrichment

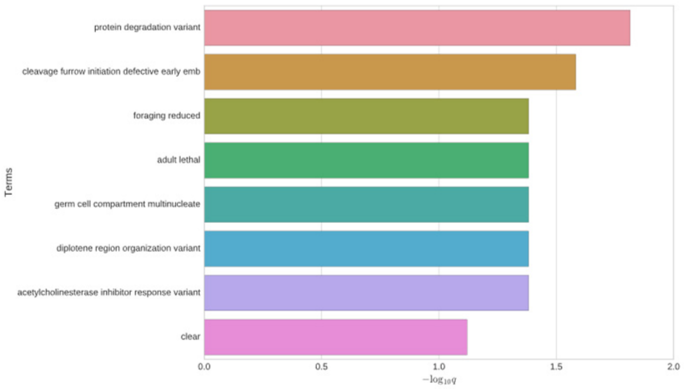


### *aman-1*

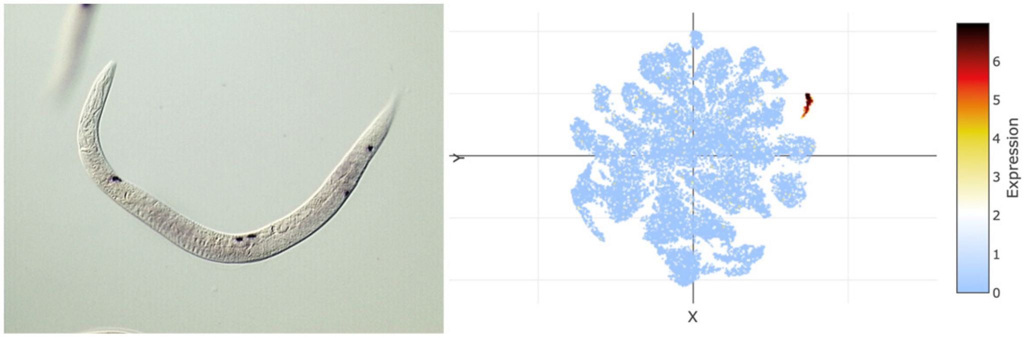


### *dig-1*

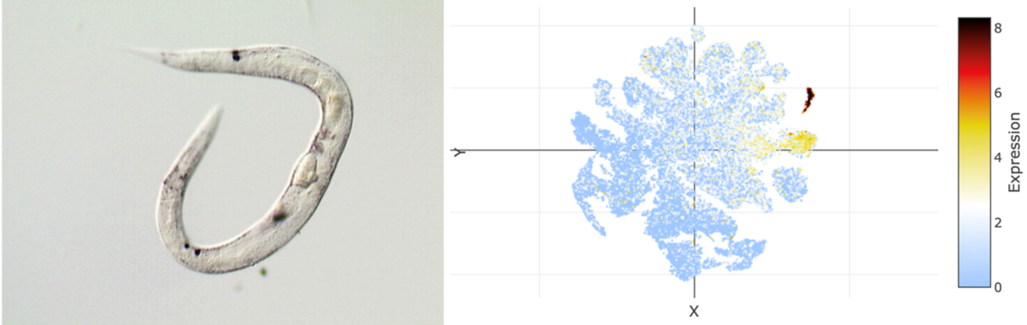

### t-SNE showing cell type 29

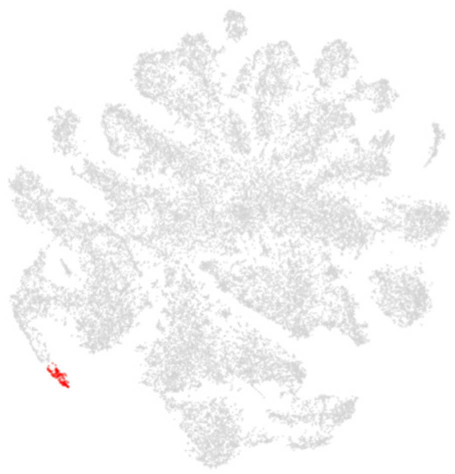

### Top 20 enriched genes

| | p_val | avg_log2FC | pct.1 | pct.2 | p_val_adj |
|---|---|---|---|---|---|
| *C35E7.5* | 0 | 4.46295818 | 0.811 | 0.037 | 0 |
| *sdz-29* | 0 | 4.46012328 | 0.396 | 0.007 | 0 |
| *odc-1* | 3.61E-68 | 4.32442019 | 0.523 | 0.084 | 1.69E-63 |
| *clec-266* | 1.13E-267 | 3.85284467 | 0.676 | 0.037 | 5.29E-263 |
| *pop-1* | 8.28E-75 | 3.53076343 | 0.55 | 0.083 | 3.88E-70 |
| *sdc-2* | 6.25E-167 | 3.40919513 | 0.739 | 0.074 | 2.93E-162 |
| *epg-2* | 7.35E-210 | 3.39618951 | 0.685 | 0.049 | 3.45E-205 |
| *cav-1* | 1.54E-121 | 3.38624962 | 0.649 | 0.075 | 7.21E-117 |
| *str-81* | 0 | 3.31258949 | 0.568 | 0.007 | 0 |
| *Y46H3C.7* | 8.18E-127 | 3.30984004 | 0.82 | 0.12 | 3.84E-122 |
| *C47D12.4* | 0 | 3.30398414 | 0.613 | 0.009 | 0 |
| *cht-1* | 6.75E-190 | 3.2026164 | 0.514 | 0.029 | 3.17E-185 |
| *Y82E9BR.17* | 0 | 3.18881549 | 0.649 | 0.01 | 0 |
| *T27A1.3* | 0 | 3.15315264 | 0.523 | 0.009 | 0 |
| *Y51H7C.15* | 4.24E-227 | 3.12242129 | 0.739 | 0.051 | 1.99E-222 |
| *his-24* | 5.65E-38 | 3.10260996 | 0.604 | 0.181 | 2.65E-33 |
| *Y45G5AM.5* | 0 | 3.09952927 | 0.441 | 0.006 | 0 |
| *chd-3* | 4.04E-80 | 3.09024482 | 0.703 | 0.138 | 1.89E-75 |
| *din-1* | 2.74E-61 | 3.07443022 | 0.784 | 0.224 | 1.28E-56 |
| *F53H1.4* | 1.14E-57 | 3.03436232 | 0.775 | 0.227 | 5.37E-53 |

### Top 20 depleted genes

| | p_val | avg_log2FC | pct.1 | pct.2 | p_val_adj |
|---|---|---|---|---|---|
| *D1086.10* | 1.84E-08 | -3.2291334 | 0.135 | 0.394 | 0.00086521 |
| *ule-4* | 9.19E-09 | -3.1411727 | 0.243 | 0.508 | 0.00043083 |
| *lev-11* | 5.65E-29 | -3.1115638 | 0.216 | 0.729 | 2.65E-24 |
| *far-2* | 2.01E-26 | -2.9362263 | 0.36 | 0.768 | 9.45E-22 |
| *col-139* | 6.05E-12 | -2.8962295 | 0.117 | 0.431 | 2.84E-07 |
| *col-124* | 4.31E-24 | -2.8686186 | 0.27 | 0.708 | 2.02E-19 |
| *col-119* | 8.24E-25 | -2.8547338 | 0.297 | 0.732 | 3.87E-20 |
| *unc-54* | 2.66E-24 | -2.833011 | 0.18 | 0.685 | 1.25E-19 |
| *ttr-16* | 1.88E-19 | -2.8327523 | 0.27 | 0.646 | 8.80E-15 |
| *col-93* | 1.21E-26 | -2.8078882 | 0.351 | 0.768 | 5.67E-22 |
| *cpn-3* | 3.41E-23 | -2.7989457 | 0.225 | 0.684 | 1.60E-18 |
| *K08D12.6* | 6.24E-21 | -2.7922375 | 0.153 | 0.604 | 2.93E-16 |
| *col-20* | 2.77E-25 | -2.7716992 | 0.252 | 0.725 | 1.30E-20 |
| *unc-27* | 2.47E-23 | -2.7640237 | 0.207 | 0.668 | 1.16E-18 |
| *unc-15* | 5.13E-23 | -2.7502167 | 0.162 | 0.649 | 2.40E-18 |
| *far-1* | 4.00E-21 | -2.72896 | 0.171 | 0.634 | 1.87E-16 |
| *act-3* | 2.61E-25 | -2.707055 | 0.387 | 0.775 | 1.22E-20 |
| *C39D10.7* | 7.01E-07 | -2.7036447 | 0.072 | 0.284 | 0.03287574 |
| *mup-2* | 2.70E-21 | -2.6529847 | 0.09 | 0.563 | 1.27E-16 |
| *col-122* | 1.98E-24 | -2.6381536 | 0.324 | 0.74 | 9.28E-20 |

### Tissue enrichment

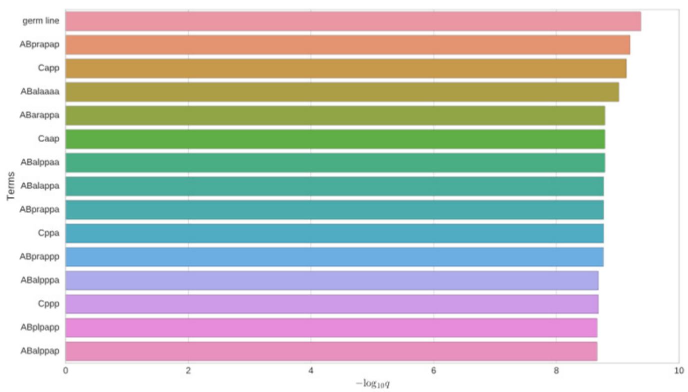


### GO enrichment

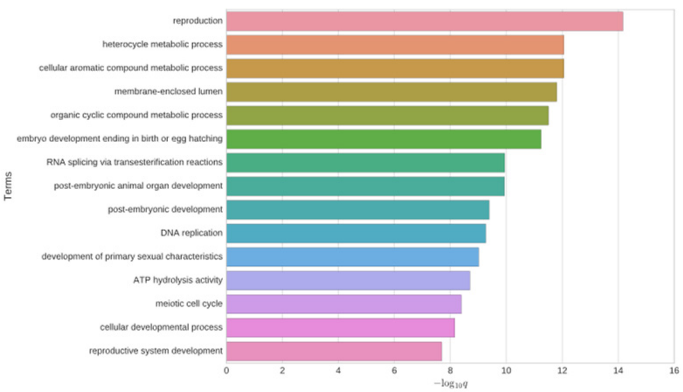


### Phenotype enrichment

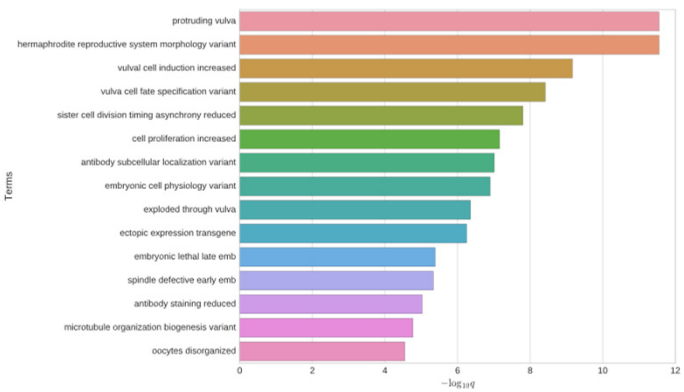


### *clec-266*

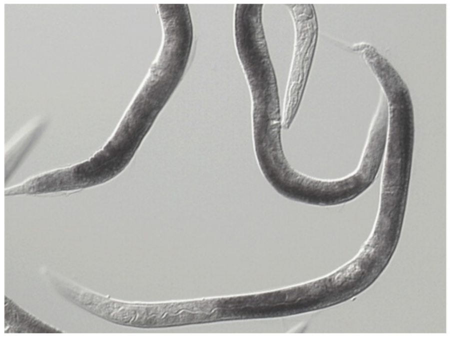

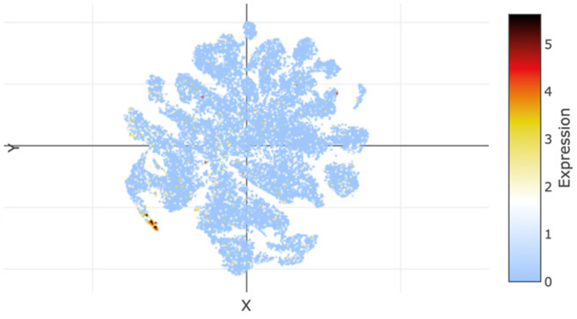


## *C35E7.5*

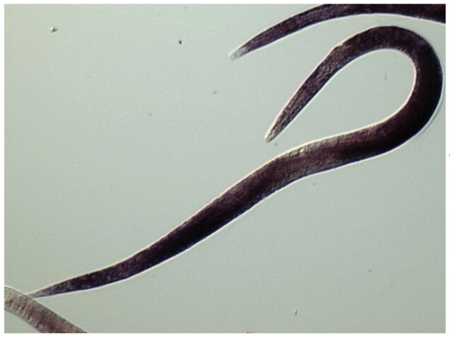

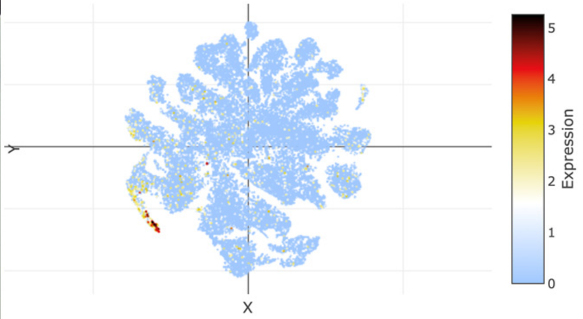

### t-SNE showing cell type 30

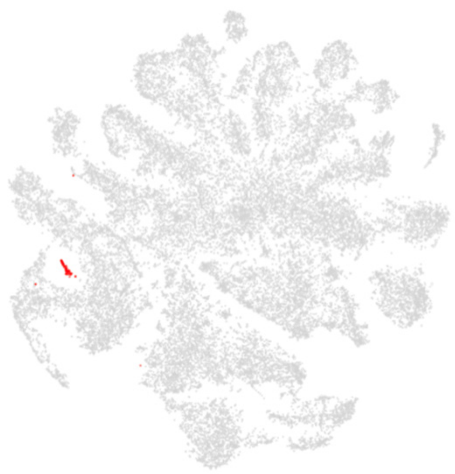

### Top 20 enriched genes

| | p_val | avg_log2FC | pct.1 | pct.2 | p_val_adj |
|---|---|---|---|---|---|
| *MTCE.33* | 4.23E-20 | 6.41200122 | 0.763 | 0.266 | 1.98E-15 |
| *ctc-1* | 9.78E-21 | 5.15495749 | 0.895 | 0.437 | 4.59E-16 |
| *ctb-1* | 3.18E-18 | 3.8397139 | 0.816 | 0.383 | 1.49E-13 |
| *nduo-5* | 2.26E-22 | 3.60902475 | 0.868 | 0.372 | 1.06E-17 |
| *ctc-3* | 1.32E-26 | 3.35728548 | 1 | 0.575 | 6.19E-22 |
| *ctc-2* | 6.22E-24 | 3.32558884 | 0.947 | 0.439 | 2.92E-19 |
| *nduo-4* | 1.60E-24 | 3.14952748 | 0.921 | 0.344 | 7.49E-20 |
| *atp-6* | 3.31E-09 | 2.8378901 | 0.763 | 0.546 | 0.00015531 |
| *nduo-1* | 7.46E-16 | 2.79468754 | 0.816 | 0.362 | 3.50E-11 |
| *fln-1* | 3.01E-10 | 2.72370835 | 0.763 | 0.537 | 1.41E-05 |
| *nduo-6* | 5.71E-15 | 2.68922005 | 0.921 | 0.676 | 2.68E-10 |
| *rla-0* | 8.48E-08 | 2.46660321 | 0.789 | 0.572 | 0.00397621 |
| *rpl-11.1* | 1.60E-10 | 2.39478235 | 0.684 | 0.315 | 7.52E-06 |
| *nduo-2* | 6.93E-10 | 2.30895059 | 0.526 | 0.182 | 3.25E-05 |
| *MTCE.7* | 7.96E-07 | 2.29792725 | 0.632 | 0.345 | 0.03733506 |
| *ppw-2* | 3.85E-13 | 2.28241747 | 0.474 | 0.124 | 1.81E-08 |
| *rpoa-1* | 2.24E-15 | 2.15638609 | 0.605 | 0.176 | 1.05E-10 |
| *rla-1* | 3.65E-09 | 2.13590711 | 0.737 | 0.402 | 0.00017135 |
| *ndfl-4* | 1.72E-07 | 2.1061077 | 0.711 | 0.473 | 0.00805659 |
| *eea-1* | 4.01E-13 | 2.04993501 | 0.658 | 0.232 | 1.88E-08 |

### Top 20 depleted genes

| | p_val | avg_log2FC | pct.1 | pct.2 | p_val_adj |
|---|---|---|---|---|---|
| *col-93* | 1.68E-15 | -5.126399 | 0.079 | 0.767 | 7.87E-11 |
| *col-119* | 2.43E-13 | -4.994781 | 0.132 | 0.731 | 1.14E-08 |
| *nlp-77* | 2.61E-14 | -4.8577085 | 0 | 0.705 | 1.22E-09 |
| *col-140* | 2.25E-14 | -4.8169383 | 0.053 | 0.738 | 1.05E-09 |
| *col-95* | 2.03E-10 | -4.7165161 | 0.026 | 0.589 | 9.51E-06 |
| *F46H5.3* | 3.08E-16 | -4.4789981 | 0.105 | 0.802 | 1.45E-11 |
| *col-20* | 1.80E-13 | -4.4578894 | 0.079 | 0.723 | 8.43E-09 |
| *pat-10* | 3.77E-13 | -4.4451127 | 0.053 | 0.708 | 1.77E-08 |
| *col-106* | 2.72E-13 | -4.4186801 | 0.026 | 0.686 | 1.28E-08 |
| *col-181* | 1.83E-13 | -4.3287726 | 0.079 | 0.726 | 8.57E-09 |
| *lev-11* | 1.40E-12 | -4.1562704 | 0.158 | 0.728 | 6.55E-08 |
| *col-19* | 1.34E-11 | -4.1228499 | 0.079 | 0.663 | 6.29E-07 |
| *act-4* | 5.58E-13 | -4.0803911 | 0.079 | 0.719 | 2.62E-08 |
| *col-122* | 1.60E-13 | -4.0798741 | 0.079 | 0.739 | 7.51E-09 |
| *unc-27* | 2.52E-11 | -4.0473375 | 0.105 | 0.667 | 1.18E-06 |
| *mlc-3* | 4.32E-12 | -3.9836271 | 0.079 | 0.683 | 2.03E-07 |
| *col-160* | 1.32E-14 | -3.9245871 | 0.053 | 0.755 | 6.19E-10 |
| *F11E6.3* | 3.90E-12 | -3.8870397 | 0.105 | 0.688 | 1.83E-07 |
| *col-8* | 9.44E-12 | -3.8131409 | 0.053 | 0.646 | 4.43E-07 |
| *col-124* | 2.08E-11 | -3.7723947 | 0.158 | 0.707 | 9.74E-07 |

### Tissue enrichment

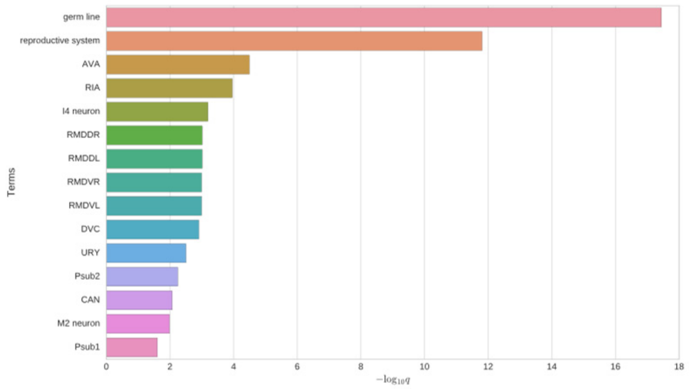


### GO enrichment

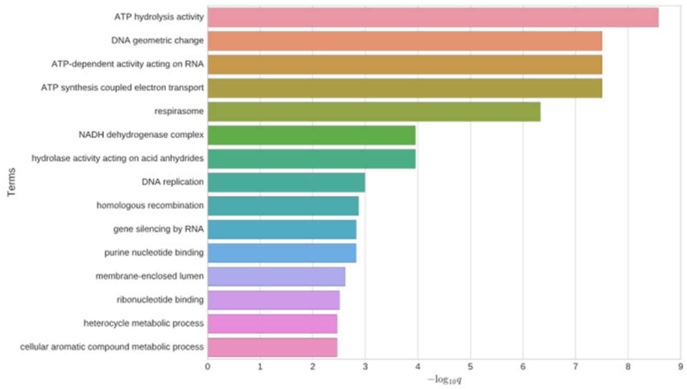


### Phenotype enrichment

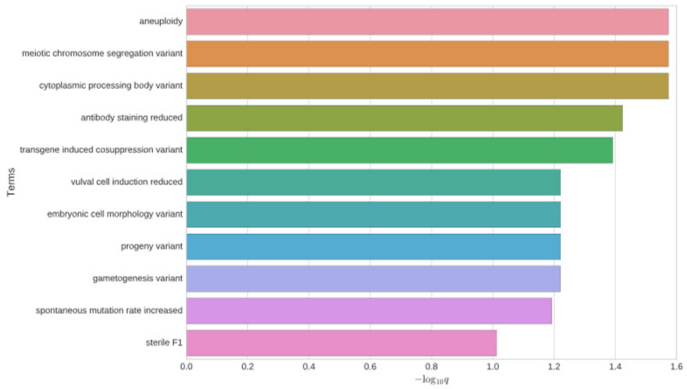


### *ppw-2*

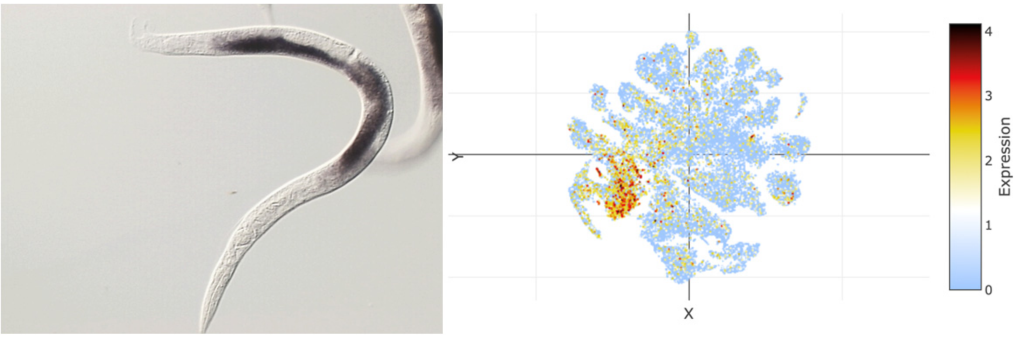


### *set-9*

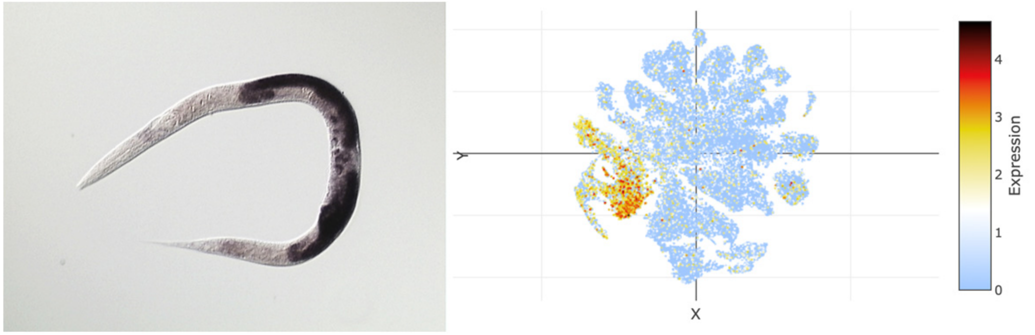
